# CYP76BK1 orthologs catalyze furan and lactone ring formation in clerodane diterpenoids across the mint family

**DOI:** 10.1101/2024.08.28.609960

**Authors:** Nicholas J. Schlecht, Emily R. Lanier, Trine B. Andersen, Julia Brose, Daniel Holmes, Björn R. Hamberger

**Affiliations:** Department of Biochemistry and Molecular Biology, Michigan State University, East Lansing, MI, USA; DOE Great Lakes Bioenergy Research Center, Michigan State University, East Lansing, MI, USA; Department of Plant Biology, Michigan State University, East Lansing, MI, USA; Department of Chemistry, Michigan State University, East Lansing MI, USA

## Abstract

The Lamiaceae (mint family) is the largest known source of furanoclerodanes, a subset of clerodane diterpenoids with broad bioactivities including insect antifeedant properties. The *Ajugoideae* subfamily, in particular, accumulates significant numbers of structurally related furanoclerodanes. The biosynthetic capacity for formation of these diterpenoids is retained across most Lamiaceae subfamilies, including the early-diverging *Callicarpoideae* which forms a sister clade to the rest of Lamiaceae. *VacCYP76BK1*, a cytochrome P450 monooxygenase from *Vitex agnus-castus*, was previously found to catalyze the formation of the proposed precursor to furan and lactone-containing labdane diterpenoids. Through transcriptome-guided pathway exploration, we identified orthologs of *VacCYP76BK1* in *Ajuga reptans* and *Callicarpa americana.* Functional characterization demonstrated that both could catalyze the oxidative cyclization of clerodane backbones to yield a furan ring. Subsequent investigation revealed a total of ten *CYP76BK1* orthologs across six Lamiaceae subfamilies. Through analysis of available chromosome-scale genomes, we identified four *CYP76BK1* members as syntelogs within a conserved syntenic block across divergent subfamilies. This suggests an evolutionary lineage that predates the speciation of the Lamiaceae. Functional characterization of the *CYP76BK1* orthologs affirmed conservation of function, as all catalyzed furan ring formation. Additionally, some orthologs yielded two novel lactone ring moieties. The presence of the *CYP76BK1* orthologs across Lamiaceae subfamilies closely overlaps with the distribution of reported furanoclerodanes. Together, the activities and distribution of the *CYP76BK1* orthologs identified here support their central role in furanoclerodane biosynthesis within the Lamiaceae family. Our findings lay the groundwork for biotechnological applications to harness the economic potential of this promising class of compounds.

**Significance Statement:** The discovery and functional characterization of *CYP76BK1* orthologs across diverse Lamiaceae subfamilies revealed novel chemistry and their central role in furanoclerodane biosynthesis, providing insights into the metabolic landscape and dynamic evolution of this plant family over approximately 50 million years. These findings pave the way for targeted biosynthetic engineering efforts and the sustainable production of furanoclerodane compounds, offering promising prospects for agricultural and pharmaceutical applications.

## INTRODUCTION

The past century has seen a tremendous explosion in the use of chemicals to treat biological threats, both in medicine and in agriculture. However, evolving challenges such as climate change and chemical resistance underscore a critical need for continued development of novel and effective biological control agents (Garcia-Solache and Casadevall, 2010; Yi *et al*., 2014; MacGowan and Macnaughton, 2017). Modern advances in genomics and metabolomics have enabled new opportunities to leverage plant natural products and their semi-synthetic derivatives to address these challenges (Han *et al*., 2024).

Diterpenoids are a structurally diverse class of specialized metabolites prevalent in plants. A limited set of diterpene backbones gives rise to thousands of uniquely oxidatively decorated diterpenoids with a broad spectrum of bioactivities. The modular nature of the biosynthetic route through action of diterpene synthases (diTPSs) and cytochrome P450 monooxygenases (P450s) is attractive from both a natural and bioengineering perspective, enabling rapid diversification of chemistries from just a few initial structures. From an evolutionary perspective, diterpenoids can serve as molecular signatures across taxa, reflecting the history of plant families, speciation, and adaptation. They are key contributors to abiotic stress responses, microbial symbioses, developmental signaling, and defense against herbivores and pathogens (Cheng *et al*., 2007). For humans, their economic importance is based on applications such as therapeutics, nutraceuticals, and in agriculture. One of the most prolific sources of diterpenoids are the Lamiaceae, also known as the mint family, recognized for highly aromatic plants. Species within this family contribute roughly 16% of the over 25,000 distinct diterpenoids reported in the Dictionary of Natural Products (DNP) (Dictionary of Natural Products v30.2).

Clerodanes, one of the most prevalent classes of diterpenoids, are of considerable economic interest due to their bioactivities. Within the Lamiaceae, numerous clerodane diterpenoids have been reported with a broad spectrum of activities including powerful opioid receptor agonists from the psychedelic plant *Salvia divinorum*, potent insect antifeedant and insecticidal compounds from the Scutellarioideae and Ajugoideae subfamilies, and MRSA-active antibacterial clerodanes in *Callicarpa americana* (Gebbinck *et al*., 2002; Roth *et al*., 2002; Dettweiler *et al*., 2020). Additional reported bioactivities of clerodanes include anti-cancer and antimicrobial therapeutics (Li *et al*., 2016). There are seven common skeletal configurations of clerodanes with modifications of carbons 11-16 typically featuring either a furan or lactone moiety— collectively called “furanoclerodanes” (Figure 1) (Li *et al*., 2016). In rare cases, other heteroatoms are incorporated in sidechains or as pyrrolidine rings (Kobayashi *et al*., 2000). Based on available structure-activity relationship data, the widespread presence of furan and lactone moieties and their variations appear to be the main drivers of the biological activities of furanoclerodanes (Gebbinck *et al*., 2002; Li *et al*., 2016; Enriz *et al*., 2000). While the furan and lactone sidechain modifications are often observed with clerodane backbones, other labdane backbones with these same features (hereafter “furanolabdanes”) have also been reported from diverse plant taxa (Kiuchi *et al*., 2004; H., Wu *et al*., 2013; X., De Wu *et al*., 2013; Li *et al*., 2016; Zhou *et al*., 2023).

**Figure 1.**
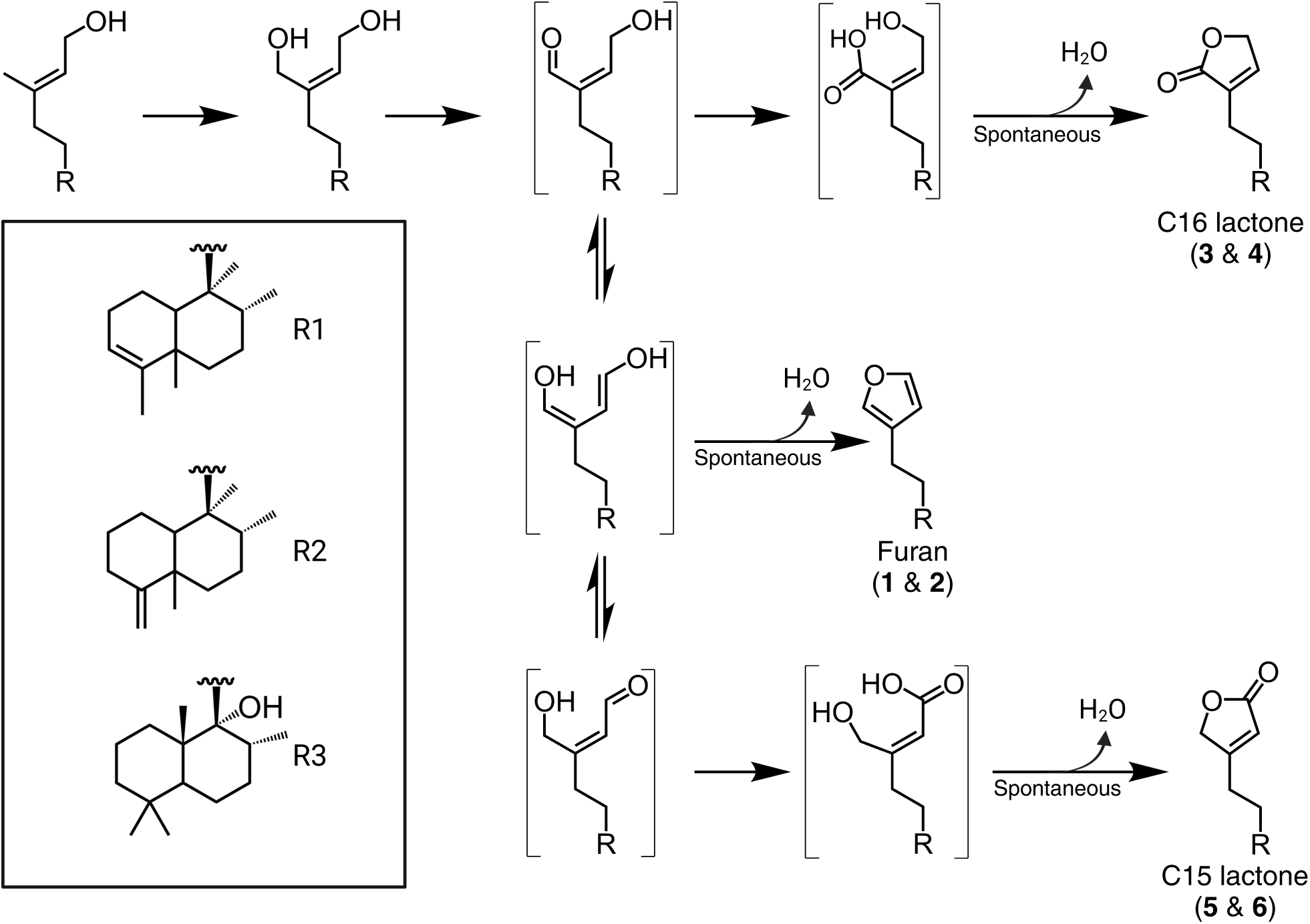
Structures of relevant clerodanes and furanoditerpenoids. A. Inset, neo-clerodane backbone with C15 and C16 highlighted for their role in furan and lactone cyclization and lactone carbonyl positioning. **I.**-**VII.**, seven types of clerodane backbones (Li *et al*., 2016). B. Salvinorin A from *S. divinorum*; Ajugarin I from *A. reptans;* 12(*S*),16ξ-dihydroxycleroda-3,13-dien-15,16-olide is a furanoclerodane from *C. americana;* Vitexilactone, example of a furanolabdane from *V. agnus-castus* (Roth *et al*., 2002; Nishina *et al*., 2017; Khan *et al*., 2023)

Recent investigations have begun to elucidate the enzymes involved in furanoclerodane biosynthesis in select species. Several diTPSs have been identified that form the key diterpene diphosphate precursors to various clerodane backbones. Within the Lamiaceae, it has been reported that *S. divinorum* (SdKPS1), *Vitex agnus-castus* (VacTPS5), *C. americana* (CamTPS2), *Scutellaria baicalensis* (SbdiTPS2.8) and *Salvia splendens* (SspdiTPS2.1) all have (-)-kolavenyl diphosphate (KPP) synthases (Pelot *et al*., 2017; Heskes *et al*., 2018; Hamilton *et al*., 2020; Li *et al*., 2023). The double bond isomer of KPP, isokolavenyl diphosphate (IKPP), is the major product of ArTPS2 from *Ajuga reptans* and two diTPSs from *Scutellaria* species (SbdiTPS2.7 and SbbdiTPS2.1, 2.3) (Johnson *et al*., 2019; Li *et al*., 2023; Qiu *et al*., 2023). More recently, a few P450s catalyzing formation of the furan moiety have been identified as well. In *S. divinorum*, *CYP76AH39* was found to modify kolavenol (dephosphorylated KPP) with a dihydrofuran moiety (Kwon *et al*., 2021). In switchgrass (*Panicum virgatum*), several P450s in the monocot-specific CYP71Z family were shown to catalyze furan ring formation on both clerodane and labdane backbones (Muchlinski *et al*., 2021). In *V. agnus-castus*, *CYP76BK1*was shown to hydroxylate C16 of the labdane peregrinol, which was suggested to be the first step in the formation of the furan ring apparent in vitexilactone (Figure 1) (Heskes *et al*., 2018).

In this work, we sought to further elucidate biosynthetic routes leading to bioactive furanoclerodanes within the Lamiaceae. We initially focused on two distinct and potentially economically significant compound classes: the Ajugarins, potent insect antifeedant furanoclerodanes produced by Ajugoideae species, and the MRSA-active furanoclerodane isolated from *C. americana* (12(*S*),16ξ-dihydroxycleroda-3,13-dien-15,16-olide) (Gebbinck *et al*., 2002; Dettweiler *et al*., 2020). These pathways, originating from evolutionarily distinct subfamilies, provided a strategic foundation for discovering a broadly connected landscape of furanoclerodane and furanolabdane biosynthesis across the Lamiaceae.

## RESULTS

### Lamiaceae species produce most plant furanoclerodanes

To gain understanding of furanoclerodane distribution patterns across all organisms, the DNP was mined for reported diterpenoids. Compounds annotated as labdanes or clerodanes bearing sidechain moieties characteristic of furanoclerodanes (i.e., structures II, III, and VI in Figure 1) were extracted. This analysis revealed that furanoclerodanes and furanolabdanes occur across 16 distinct clades across plants and some marine organisms (Figure 2). These include 10 dicotyledon families, 2 monocotyledon families, a family of ferns, a family of liverworts, and two different orders of demosponges. The majority of furanoclerodanes are found in the Lamiaceae family. Notably, there are over five times more furanoclerodanes than furanolabdanes across all families, revealing a strong bias towards the clerodane backbone.

**Figure 2.**
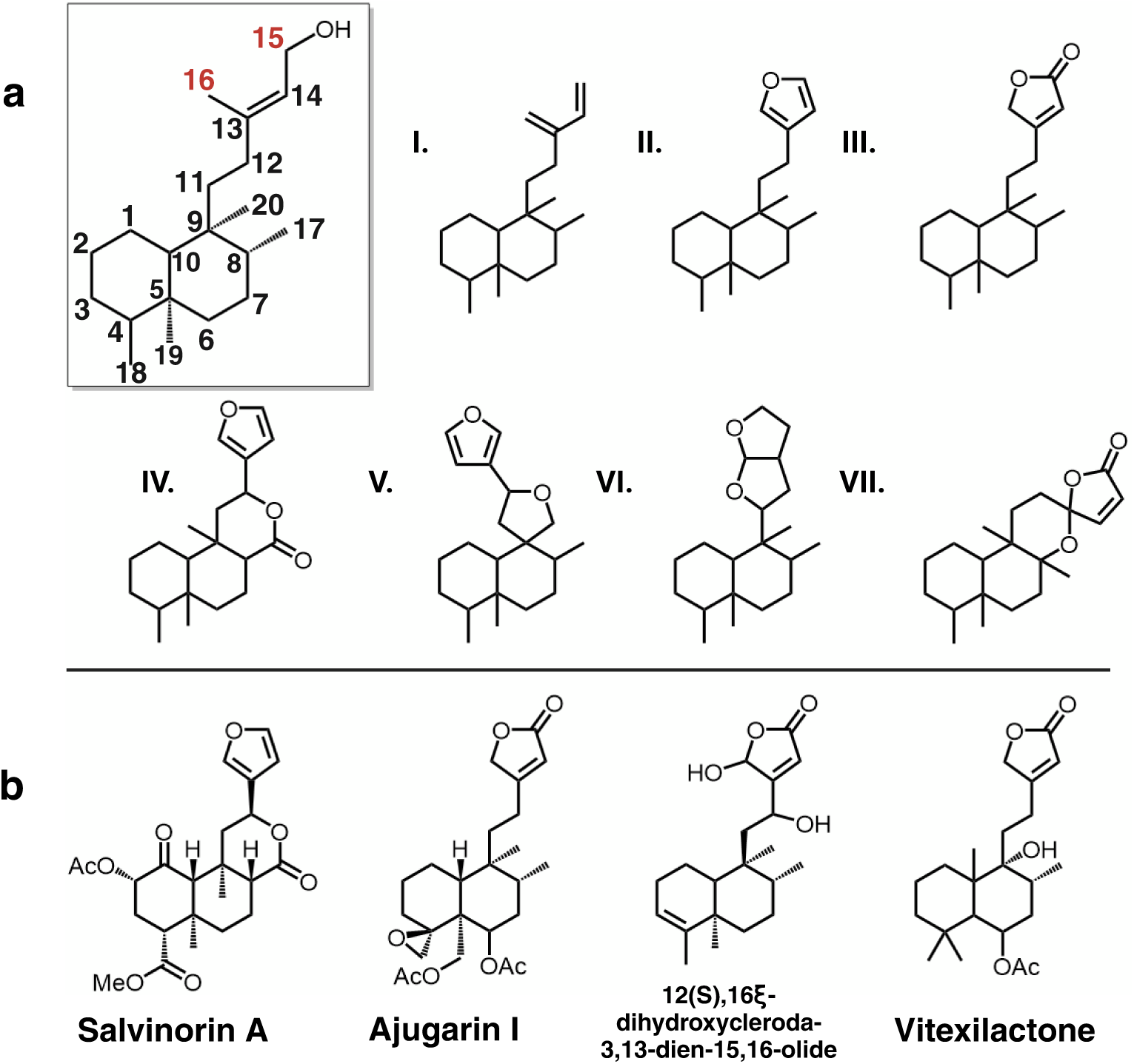
Furanoclerodane and furanolabdane abundances. Furanoclerodanes, blue; furanolabdanes, gray. Bars represent the number of unique compounds in the respective clade. The panel on the right shows the enrichment in the Lamiaceae. The percentages next to subfamilies compare the furanoclerodanes/furanolabdanes found in that subfamily to all recorded diterpenoids of that clade. The genus level bar plot was filtered to 3 and above compounds.

Within Lamiaceae, furanoclerodane/labdanes are found in eight of eleven subfamilies but vary in the abundances of uniquely decorated structures (Figures S1 and S2).

Callicarpoideae contains roughly a dozen unique furanoclerodanes while the Ajugoideae has several hundred. While the Nepetoideae subfamily contains approximately half of all Lamiaceae species, its *Salvia* genus is the only one with reported furanoclerodanes. The asymmetric diversity paired with presence of furanoclerodanes in evolutionarily distant subfamilies suggests there may be a basal, conserved metabolic pathway within the Lamiaceae.

### Identification of two P450s catalyzing production of furanoclerodanes in A. reptans and C. americana

To investigate whether a furanoclerodane biosynthetic pathway is conserved across Lamiaceae, we analyzed two taxonomically divergent species, *C. americana* from the early-diverging Callicarpoideae subfamily, as well as *A*. *reptan*s, member of the Ajugoideae. Cytochrome P450 monooxygenases, particularly members of the CYP71 clan, can catalyze oxidative tailoring reactions in plant diterpenoid biosynthesis. Protein blast homology with a set of reference CYP71s was used to identify candidates from transcriptomic (*A. reptans*) and genomic (*C. americana*) data, yielding roughly 250 candidates. Candidate genes in *C. americana* were selected using tissue-specific expression data to correlate expression of candidates with that of the KPP synthase *CamTPS2* (Figure S3). In *A. reptans* only expression data from leaf tissue was available, so candidates were chosen based on strength of expression and clustering with the CYP76 subfamily, which is prominent in Lamiaceae diterpenoid metabolism (Bathe and Tissier, 2019). This approach yielded eight candidates from each species for functional characterization.

Candidate P450 transcripts were cloned and transiently expressed in *Nicotiana benthamiana* along with upstream terpene precursor genes D-xylulose-5-phosphate synthase (DXS) and geranylgeranyl diphosphate synthase (GGPPS) along with either CamTPS2 or ArTPS2. Consistent with previous studies, endogenous non-specific phosphatase activities were sufficient to provide the substrate for P450 oxidation (Pelot *et al*., 2017; Johnson *et al*., 2019; Muchlinski *et al*., 2021; Kwon *et al*., 2021). Most of the initial candidates did not convert the clerodane substrates. However, the orthologs *ArCYP76BK1* and *CamCYP76BK1* were found to catalyze formation of two new products (Figure S4), identified by NMR as 15,16-epoxy-4,18,(16),14-clerodatriene (**1**) and 15,16-epoxy-3,13(16),14-clerodatriene (**2**). These result from furan ring cyclization of the isokolavenol and kolavenol backbone, respectively (Figures S5-S8). These compounds are likely intermediates in the pathway toward the bioactive clerodanes (Figure 1). This transformation echoes the activity of previously identified CYP71Zs from switchgrass and *CYP76AH39* in *S. divinorum* (Kwon *et al*., 2021). *ArCYP76BK1* and *CamCYP76BK1* exhibit high protein sequence identity (71-74%) as well as functional similarity to *VacCYP76BK1*, despite approximately 50M years of evolutionary distance between the Callicarpoideae, Viticoideae, and Ajugoideae subfamilies (Kumar *et al*., 2022). This discovery prompted further investigation into the significance of *CYP76BK1* orthologs in furanoclerodane production across other Lamiaceae subfamilies.

### Exploration of the CYP76BK family across the mint family

The set of 48 transcriptomes generated by the Mint Genome Project, which provides widespread coverage across subfamilies, was investigated for the presence of CYP76 family members (Figure 3a) (Boachon *et al*., 2018). An additional transcriptome was generated for *Teucrium chamaedrys*. Phylogenetic analysis identified a monophyletic CYP76BK clade containing sequences from species in the Premnoideae, Ajudoideae, Peronematoideae, Scutellarioideae, Viticoideae, and Callicarpoideae (Figure S9, Figure 3b). This revealed an additional seven species with *CYP76BK1* orthologs, named according to species *CpCYP76BK1* (*Cornutia pyramidata*), *PbCYP76BK1* (*Petraeovitex bambusetorum*), *HsCYP76BK1* (*Holmskioldia sanguinea*), *SbCYP76BK1* (*S. baicalensis*), *CbCYP76BK1* (*Clerodendrum bungei*), *Teucrium canadense* (*TcaCYP76BK1*) and *T. chamaedrys* (*TchCYP76BK1)*.

**Figure 3.**
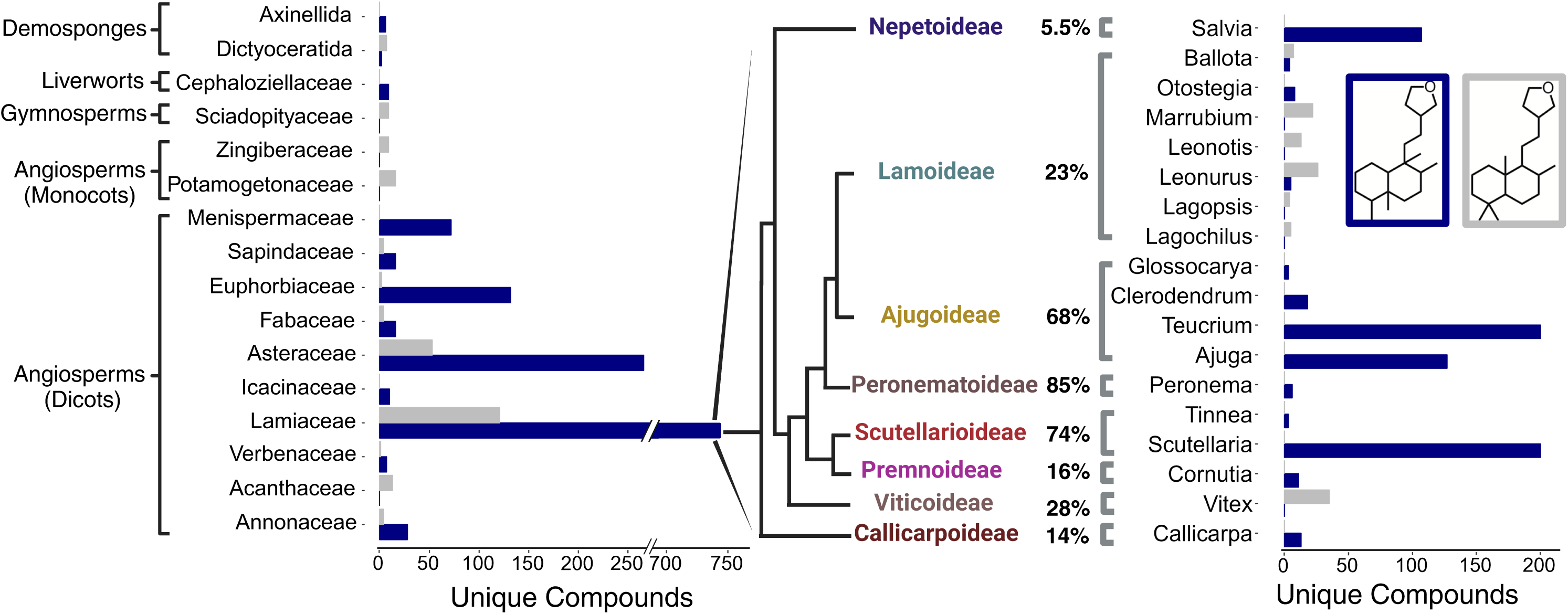
Identification of CYP76BK1 orthologs from the Lamiaceae family. (a) Phylogeny of the 48 Lamiaceae representatives screened for *CYP76BK1* orthologs. (*) Indicates subfamilies with at least one CYP76BK1 ortholog (Figure adapted from Boachon *et al*., 2018). (b) Maximum likelihood tree of CYP76BK1 orthologs along with reference sequences from the CYP76 family (in black). Black dots indicate a bootstrap value of 70% or greater (1000 bootstraps). CYP76AH39 is implicated in furanoclerodane biosynthesis in *Salvia.* Reference P450s can be found in Table S1.

### Functional characterization of CYP76BK1 orthologs reveals alternative cyclization products

Newly identified *CYP76BK1* transcripts were cloned from leaf tissue cDNA and evaluated using transient expression in *N. benthamiana.* Each ortholog was co-expressed with a diTPS to assess activity with both kolavenol and isokolavenol (Figure 4). GC-MS analysis of the resulting enzyme products revealed that all CYP76BK1 enzymes exhibited similar activity as ArCYP76BK1 and CamCYP76BK1, generating peaks corresponding to retention times and fragmentation patterns of **1** and **2**. CamCYP76BK1 and ArCYP76BK1 product profiles revealed additional small products insufficient for purification and structural elucidation. These peaks were far more abundant with expression of other orthologs, allowing more complete analysis. Compounds **3**, **4**, and **5** were elucidated by NMR analysis (Figures S5-8, S10-15), revealing a C16 lactone moiety on **3** and **4** while **5** presents a C15 lactone arrangement. These were formally assigned as 4(18)- clerodadien-16,15-olide (**3**), 3,13-clerodadiene-16,15-olide (**4**) and 4(18)-clerodadien-15,16-olide (**5**). Compound **6** was tentatively characterized as 3,13-clerodadien-15,16-olide based on analogous mass fragmentation patterns and retention time to **5.**

**Figure 4.**
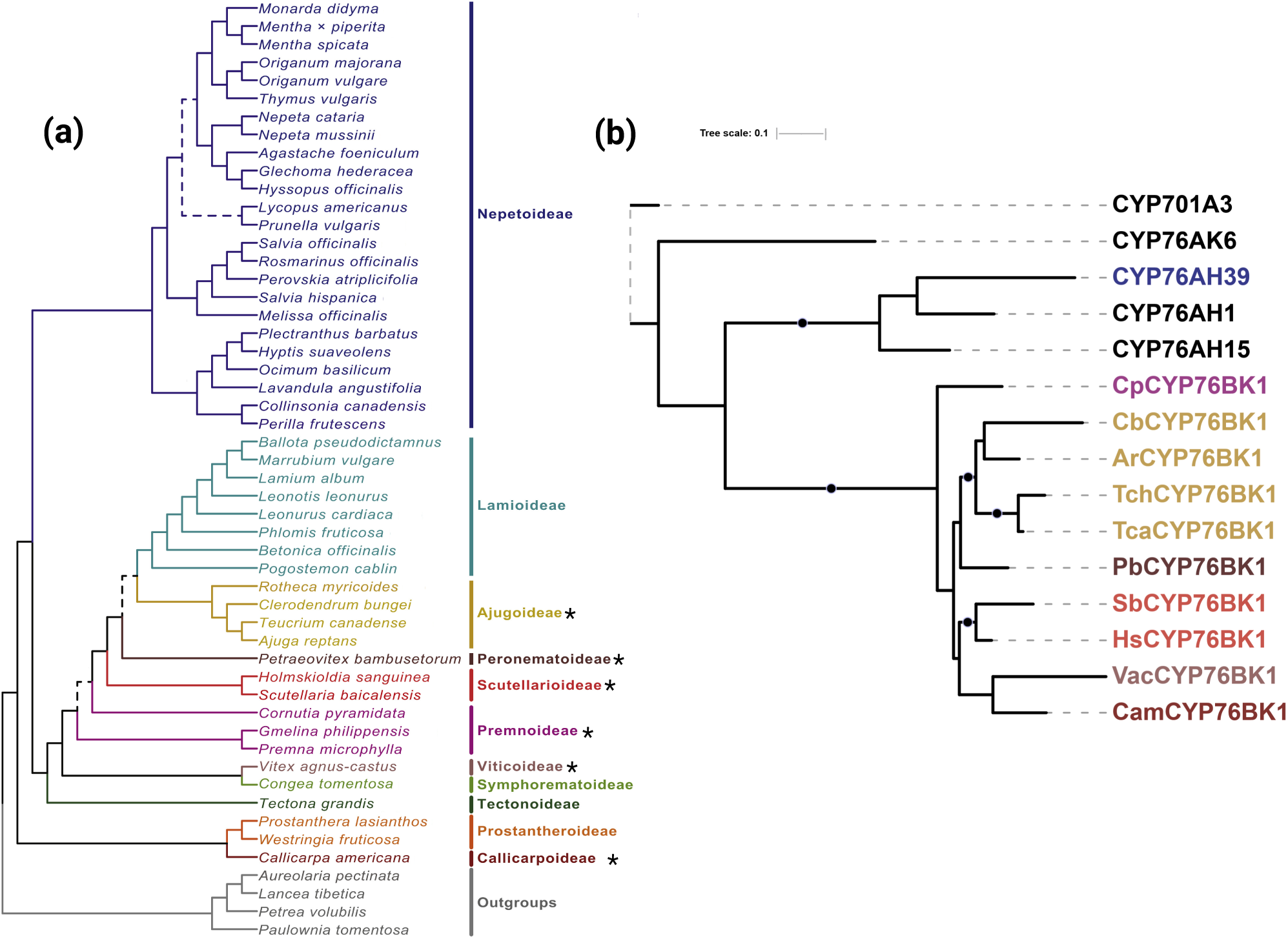
Functional characterization of CYP76BK1 orthologs. Total ion chromatograms with absolute intensities from extracts of *N. benthamiana* leaves transiently expressing each P450 candidate with the two different clerodane diTPS. Chromatograms are shown from minutes 12-18, offset by 0.1 minutes and are scaled to the highest peak in each set. Each infiltration also included a separate construct for DXS + GGPPS to increase yields. Mass spectra of compounds **1-6** can be found in Figures S16 and S17. (a) ArTPS2, an isokolavenol synthase, expressed with each ortholog. (b) CamTPS2, a kolavenol synthase, expressed with each ortholog. Compounds **1-5** have NMR data to support their structures (Figures S5-8, S10-15). The structure of compound **6** is proposed based on similar spectra and retention time to **5**.

*VacCYP76BK1* was earlier shown to generate the C16 hydroxylation of peregrinol. This was suggested as precursor to the furanolabdanes rotundifuran and vitexilactone found in *V. agnus-castus* g. We co-expressed *VacCYP76BK1* and *CYP76BK1* orthologs with the peregrinol diphosphate synthase LlTPS1 from *Leonotis leonurus* (Johnson *et al*., 2019) and could detect traces of products with mass spectra consistent with the furan and lactone derivatives of peregrinol (Figure S18), but products were not in sufficient quantities for NMR analysis. The putative peregrinol furan was also detectable with all orthologs except *CamCYP76BK1,* while the putative C15 and C16 lactones were only detectable with expression of *VacCYP76BK1, HsCYP76BK1, CbCYP76BK1,* and *ArCYP76BK1*.

Notable differences in activity can be observed among the orthologs. HsCYP76BK1 afforded the best conversion of kolavenol to **1**, while CbCYP76BK1 was most active against isokolavenol with the best yield of **4** and SbCYP76BK1 afforded the highest yield of **6**. There is apparently some bias among orthologs in which lactone variation dominates the product profile, with CbCYP76BK1, CpCYP76BK1, and HsCYP76BK1 preferentially catalyzing formation of the C16 lactone while expression of PbCYP76BK1, SbCYP76BK1, and TchCYP76BK1 led to higher accumulation of the C15 lactone. Relative activities of ArCYP76BK1 and CamCYP76BK1 were more difficult to distinguish due to lower overall activity in lactone formation. Overall, there appears to be flexibility in the acceptance of both kolavenol and isokolavenol substrates (Figure S19). Neither CamCYP76BK1 nor VacCYP76BK1 appear capable of converting isokolavenol to either the furan or lactone products, but both show moderate activity with kolavenol.

### Analysis of plant extracts

We analyzed leaf extracts of species with *CYP76BK1* orthologs by GC-MS for evidence of the furan and lactone containing intermediates. However, only the leaf extract of *C. americana* had peaks with the same mass spectrum and retention time as **2**, **4**, and **6** (Figure 5). Other extracts showed peaks which did not match the CYP76BK1 products but may represent further modified furanoclerodane products, based on mass fragmentation patterns (Figure S20).

**Figure 5.**
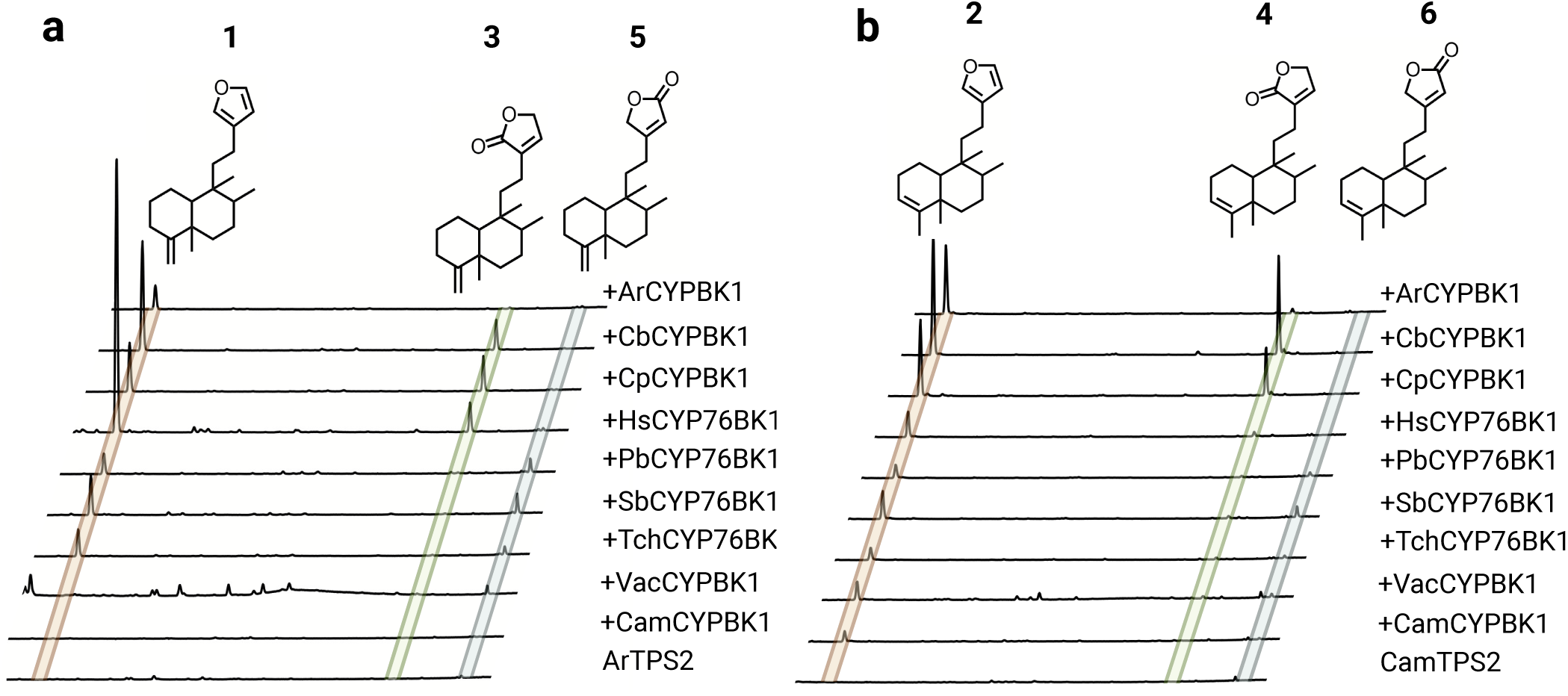
GC-MS analysis of plant extracts. (a) EIC showing the presence of the furan (m/z 286) and both lactone (m/z 302) moieties on the kolavenol backbone in the *C. americana* leaf extract. Purified products derived from *N. benthamiana* leaf extracts expressing either CamTPS2 and CbCYP76BK1 (**2** and **4**) or CamTPS2 and SbCYP76BK1 (**6**) were used for reference. (b) Deconvoluted mass spectra of **2**, **4**, and **6**. Top (black) from purified products, bottom (gray) corresponding peak from the *C. americana* leaf extract.

### Evaluating syntenic orthologs of CYP76BKs

To further understand the context and evolutionary origin of the *CYP76BK1* s, we examined available chromosome-scale genomes to determine syntenic relationships of *CYP76BK1* s in the Lamiaceae family. The genomes of *C. americana* (Hamilton *et al*., 2020), *Clerodendrum inerme* (He *et al*., 2022), *Pogostemon cablin* (Shen *et al*., 2022), *Salvia hispanica* (Brose *et al*., 2024)*, miltiorrhiza* (Pan *et al*., 2023), *Salvia officinalis* (Li *et al*., 2022), *Salvia splendens* (Dong *et al*., 2018), *S. baicalensis* (Q., Zhao *et al*., 2019), *Scutellaria barbata* (Xu *et al*., 2020), *Tectona grandis* (D., Zhao *et al*., 2019), *Thymus quinquecostatus* (Sun *et al*., 2022), and *Lavandula angustifolia* (Hamilton *et al*., 2023) were all selected based on their quality and subfamily membership: one Callicarpoideae, one Ajugoideae, one Lamioideae, two Scutellarioideae, one Tectonoideae, and six Nepetoideae.

Genomic investigations supported the transcriptomic analyses, as we found *CYP76BK1*members present in the Ajugoideae, Callicarpoideae, and Scutellarioideae subfamilies while absent in the Lamioideae, Nepetoideae, and Tectonoideae representatives. The *CYP76BK1s* in *C. americana, C. inerme, S. baicalensis,* and *S. barbata* were syntenic orthologs (syntelogs) (Figure 6). While the syntenic block is found throughout all the available Lamiaceae genomes, *CYP76BK1* homologs are not represented within the Lamioideae, Nepetoideae, and Tectonoideae.

**Figure 6.**
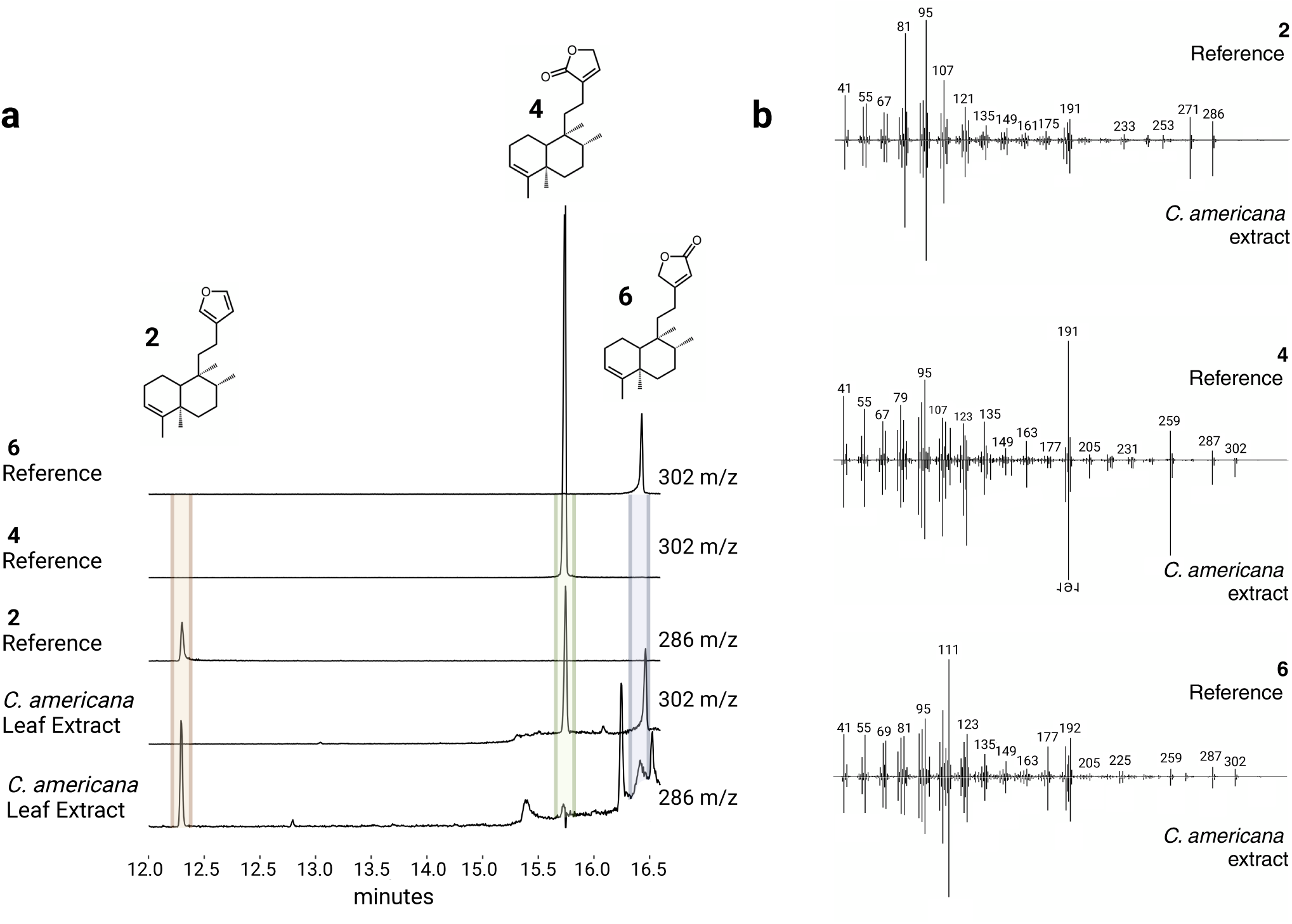
CYP76BK1 Syntenic Block Through the Lamiaceae. Species tree phylogeny of representative species of subfamilies across the Lamiaceae generated in (Brose *et al*., 2024). The red lines indicate syntenic orthologs of *CYP76BK1* present in *C. americana, S. baicalensis, S. barbata,* and *C. inerme*. Large syntenic blocks between the species are colored grey. Individual genes are colored black along the segments. The brackets highlight which syntenic blocks corresponding to each subfamily.

## Discussion

Furanoclerodanes have garnered particular interest due to their potent insect-antifeedant, antimicrobial, and other bioactivities, which likely arise from characteristic furan and lactone ring systems. Our findings establish CYP76BK1 as a key enzyme in the biosynthesis of furanoclerodanes across many distinct subfamilies of the Lamiaceae family, opening new avenues for bioengineering and biosynthetic use of furanoclerodanes.

According to compounds reported in the DNP, furanoclerodanes are found in 16 families including some sessile marine organisms as well as liverworts, monocots, and dicots. Despite the widespread occurrence and diversity, approximately 40% of all reported furanoclerodanes are from species that carry a *CYP76BK1* ortholog. The density of unique furanoclerodanes in a narrow set of species, particularly in the Ajugoideae and Scutellarioideae, highlights a rapid expansion into this metabolic niche. Across the Lamiaceae family, the distribution of *CYP76BK1* orthologs significantly overlaps with the occurrence of reported furanoclerodanes. We found orthologs in the Scutellaroideae, Ajugoideae, Callicarpoideae, Premnoideae, Peronematoideae, and Viticoideae subfamilies, all of which have species with reported furanoclerodanes. *H. sanguinea* (*HsCYP76BK1*) and *P. bambusetorum*(*PbCYP76BK1*) have no reported diterpenoids. However, with only limited phytochemical studies available in these species (Helfrich and Rimpler, 1999; Chaudhuri *et al*., 2004), the presence of *CYP76BK1* could indicate a potential source for new furanoclerodane structures.

In contrast, only a small proportion of reported furanoclerodanes from the Lamiaceae come from subfamilies without a *CYP76BK1* ortholog. Most notably, *Salvia* is the sole genus enriched in furanoclerodanes within the Nepetoideae. This can be explained by the presence of *CYP76AH39*, a dihydrofuran synthase from *S. divinorum* which has convergently evolved to perform a similar function to the *CYP76BK1* clade. We found in our phylogenetic analysis that putative *CYP76AH39* orthologs are limited to *Salvia* species (Figure S9). Moreover, recent work found through genomic and phylogenetic analysis of diterpene synthases at least two lineages leading to the appearance of clerodane biosynthesis in Lamiaceae, with the Nepetoideae lineage evolutionarily distinct from other subfamilies (Li *et al*., 2023). Together, the evolution of clerodane diterpene synthases and furan/lactone yielding P450s highlight convergent evolution of these multi-step biosynthetic pathways. Outside of *Salvia*, the Lamioideae subfamily also contains a small number of furanoclerodanes despite lacking CYP76BK1 orthologs in the transcriptomes we examined. It is plausible that the Lamioideae may have another enzyme or set of enzymes responsible for the furan ring formation. The multiple emergences of these compounds within and outside the Lamiaceae appears to underscore a selective advantage (Li *et al*., 2016; Tiedge *et al*., 2022).

Genomic analysis provided further context for the evolutionary history of CYP76BK1. The syntelogs are found in a syntenic block conserved across all available chromosome-scale genomes in the Lamiaceae, however *CYP76BK1* was lost multiple times including in the Tectonoideae, the Lamioideae and Nepetoideae subfamilies. This suggests that *CYP76BK1* emerged prior to the divergence of the major Lamiaceae subfamilies, thus constituting a foundational part of the diterpenoid diversity in Lamiaceae. Subsequent loss can be attributed to localized genomic deletions rather than larger rearrangements. Analysis of the syntenic block with PlantiSmash (Kautsar *et al*., 2017) found no additional terpene biosynthetic genes, in contrast to our previous discovery of an ancestral miltiradiene biosynthetic gene cluster present throughout the Lamiaceae (Bryson *et al*., 2023).

The functional capabilities of the *CYP76BK1* orthologs demonstrated here expand on the previous discovery of the founding member *VacCYP76BK1*. Initially, this ortholog was shown to catalyze hydroxylation of C16 of a labdane diterpenoid, peregrinol (Heskes *et al*., 2018). Utilizing GC-MS we found that this ortholog also can install both lactone and furan rings on clerodane and labdane substrates, albeit with low activity (Figure S18). The putative peregrinol furan was also detectable with all orthologs. We propose an updated mechanism for CYP76BK1 that accounts for the formation of the C15 and C16 lactone rings as well as the originally proposed furan ring (Figure 7). While a single oxidation results in the hydroxylated C15 product, a second oxidation allows spontaneous formation of the furan. A third oxidation then enables formation of the two lactones, depending on the equilibrium state. This mechanism aligns with the product profiles we observed, where the furan is the dominant product. It can also explain how different orthologs might quickly evolve the ability to synthesize one lactone over another by favoring a specific intermediate for the final oxidation.

**Figure 7.**
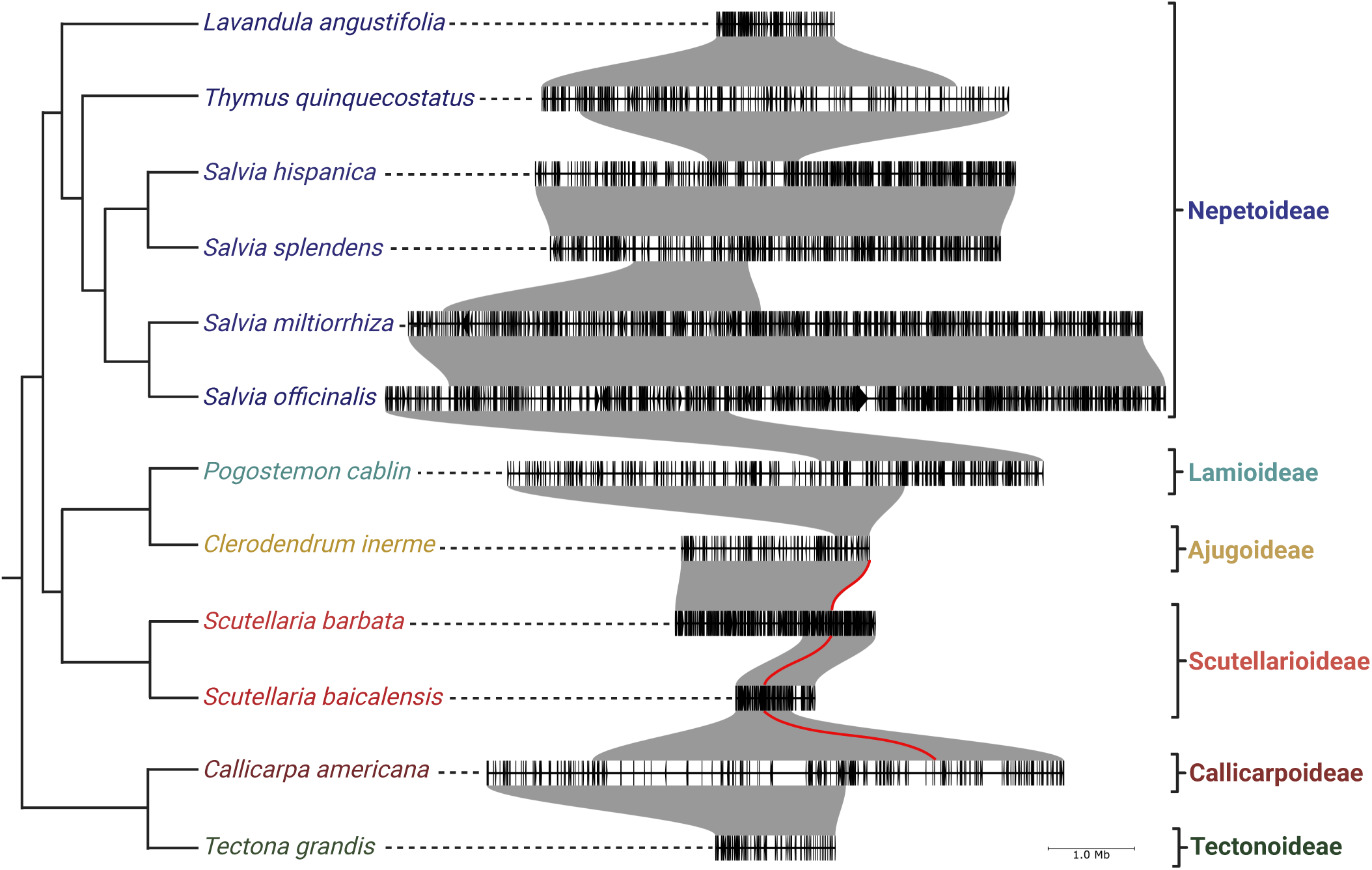
Proposed enzyme mechanism. The initial oxidation of C16 was observed with VacCYP76BK1 (Heskes *et al*., 2018). Bracketed structures represent intermediates. Formation of a carbonyl group helps delocalize the electrons, making further rearrangements feasible and permitting an opportunity for cyclization. A nucleophilic attack and dehydration can generate the furan or a subsequent oxidation on either aldehyde intermediate followed by a nucleophilic attack and dehydration will yield the respective lactone ring, leading to either the C15 or C16 lactone. (Solid arrows indicate enzymatic oxidation and dashed arrows indicate spontaneous rearrangement)

Comparison of the product profiles observed here with reported metabolites suggests that additional enzymes beyond CYP76BK1 likely govern final product profile *in planta*. The transient expression of *CYP76BK1* orthologs generally showed the furan as the dominant product, followed by the C16 lactone or the C15 lactone. Most of the structures reported in the Ajugoideae, Scutellaroideae, and Callicarpoideae contain the C15 lactone or furan, as seen in the Ajugarins, the MRSA-active clerodane in *C. americana*, and vitexilactone from *V. agnus-castus*. Type III and type VIII clerodanes appear to be derived from this C15 lactone intermediate. In contrast, only a very few reported structures contain the C16 lactone arrangement. However, the furanofuran or type VI clerodanes, which are highly abundant in *Clerodendrum*, could plausibly arise from the C16 lactone as an intermediate (Figure S1). The remaining furanoclerodane scaffolds, including types II, IV, and V appear to be derived from the furan intermediate. In metabolomic analysis of the plant extracts, we identified **2**, **4**, and **6** only in extracts from *C. americana*. This supports the biological relevance of these CYP76BK1 products, although it is interesting to note that CamCYP76BK1 itself had very low activity in the conversion to both **4** and **6**. Given the suggested multi-oxidation mechanism of CYP76BK1 enzymes, it is possible that other oxidases may be present *in planta* which are capable of steering product outcome towards the lactones. The lack of detectable CYP76BK1 products in the leaf extracts of other species may indicate that other tissues favor accumulation of these products, or that external stress is needed to initiate their biosynthesis. Alternatively, these compounds may represent intermediates in more complex pathways, potentially involving additional tailoring oxidases, shifting the product outcome towards more highly decorated species-specific diterpenoids.

The range of catalytic activities in the CYP76BK1 clade builds on recent discoveries of CYP76AH39 in *S. divinorum* and the CYP71Z clade in switchgrass (Muchlinski *et al*., 2021; Kwon *et al*., 2021). While CYP76AH39 can add a dihydrofuran to the kolavenol backbone, the CYP71Zs catalyze furan formation on multiple labdane and clerodane backbones. The CYP76BK1s further expand this biosynthetic toolbox with the ability to generate two lactone variants in addition to a furan on kolavenol, isokolavenol, and to a lesser degree some labdane substrates (Figure S18, Figure S21). Together these findings represent three distinct evolutionary trajectories towards furanoclerodane and furanolabdane biosynthesis. The substrate promiscuity demonstrated here among kolavenol, isokolavenol, and peregrinol substrates further supports the shared origin of the CYP76BK1 orthologs. For some species, such as those in the Ajugoideae, the reported furanoclerodanes appear largely isokolavenol-derived. Since *A. reptans* has only an iso-KPP synthase, while others such as C. americana have only a KPP synthase, we can surmise that the availability of these substrates likely drives product outcome more than the enzyme selectivity (Johnson *et al*., 2019; Hamilton *et al*., 2020).

A further observation from the DNP analysis is that across all plant families, furanoclerodanes outnumbered their labdane counterparts by over five-fold. Moreover, very few clerodanes are reported which lack a furan-containing moiety, while furanolabdanes comprise only a small proportion of all labdane structures reported. The pronounced skew towards furanoclerodanes may indicate selective pressures favoring the clerodane scaffold for furan and lactone ring installation, which would be consistent with the diverse bioactivities found in furanoclerodanes, most notably their insect antifeedants (Sosa *et al*., 1994; Gebbinck *et al*., 2002; Koul, 2008). We speculate that in addition to potential selective pressures from their corresponding bioactivities, the dominance of the furanoclerodanes could relate to the apparent lack of a clerodane-specific class I diTPS in most instances. Despite wide availability of transcriptomic and genomic resources, and numerous reports of class II diTPSs which form clerodienyl diphosphates, there is a general lack of corresponding class I diTPSs to facilitate diphosphate removal (Chen *et al*., 2017; Pelot *et al*., 2017; Johnson *et al*., 2019; Hamilton *et al*., 2020). It remains unclear why this pattern holds true across divergent species, but class I diTPSs typically introduce additional cyclizations in labdane pathways. In the absence of such a class I partner, plants may utilize alternative enzymes such as phosphatases or nudix hydrolases to dephosphorylate the diphosphate backbone. After dephosphorylation, P450s may evolve to oxidize these uncyclized sidechains, spontaneously forming furan and lactone scaffolds which confer a biological advantage.

Collectively, our work provides a genomic, functional, and metabolomic perspective on the central role of CYP76BK1 in furanoclerodane metabolism across Lamiaceae. These enzymes catalyze key oxidative cyclizations en route to a vast array of bioactive diterpenoid natural products. The conservation and phylogenetic distribution pattern of CYP76BK1 implicates it as an ancestral enzyme lineage facilitating the proliferation of furanoclerodanes throughout the evolution of the Lamiaceae family. Our findings establish a robust molecular toolbox for targeted engineering of furanoclerodane biosynthetic pathways, enabling sustainable production of these high-value compounds for pharmaceutical and agricultural applications

## Experimental procedures

### Survey of diterpenoids from the DNP

The Dictionary of Natural Products (v30.2) was mined for relevant diterpenoids using the following search criteria. The search category ‘Type of Compound’ with either V.S.55000 or V.S.54000, which correspond to clerodane and labdane diterpenoids respectively. Multiple subsets of the data were extracted by including specific substructures to isolate particular categories of furanoclerodanes. The substructures consisted of the side chain of the C15/C16 lactones (with and without the C14-C15 double bond), furan, and furanofuran, the commonly found VI substructure in Figure 1. CSV files were semi-automatically extracted to include the following data: Chemical Name, Molecular formula, Accurate mass, Type of Compound, Type of Organism, and Biological Source. A final control subset of data was included where the only search parameter was ‘Type of Compound words’ and the value was ‘*diterpen*’ to extract all compounds annotated as diterpene or diterpenoid.

The CSVs were imported into R (v4.2.2) and tidyverse 2.0.0 was used for varying analyses. Each dataset had 2 new columns appended to represent their backbone (furanoclerodane, furanolabdane, clerodane, labdane, or other diterpenoid) as well as which modification they have (furan, C15 lactone, C16 lactone, furanfuranofuran, non-furanoditerpenoid). The files were then concatenated. Duplicate lines were removed along with compounds lacking biological source or type of organism data. Duplicate chemical names placed in ‘other diterpenoid’ were also removed.

The biological source data was manually curated to remove tissue related data due to its nonuniform categorizations. The ‘Type of Organism’ category was divided into Kingdom, Phylum, Order, Class, and Family. Due to some DNP entries predating their species’ reclassification, various DNP entries were reclassified to contemporary clades, primarily moving various Verbenaceae to Lamiaceae. Data was grouped by either family or genus, with duplicate chemical name found within groups removed to ensure only one unique entry per group. The sum unique compounds were then plotted using different categorical compound descriptors respective of groupings. This includes comparing backbones in Lamiaceae (figure 2), which furano-moiety modifications are in Lamiaceae (Figure S1), and clerodane, labdane, furanoclerodanes/labdanes, and other diterpenoids that can be found in Lamiaceae (Figure S2).

### Candidate gene selection

Previously assembled genomic and tissue-specific expression data (Hamilton *et al*., 2020) were used to identify candidate genes in *C. americana*. The heatmap of candidate gene expression was generated using Heatmapper (Babicki *et al*., 2016). For all other species, previously assembled transcriptomic data were used (Evolutionary Genomics Consortium *et al*., 2018) (Table S2). Candidate P450s (Table S3) are initially filtered to based on 45% identity and an E value greater than 1E-5 using BLASTP against a set of reference sequences (Table S1). For *A. reptans* the candidates were further narrowed down to CYP76s exclusively. Finding orthologs of the other CYP76BKs followed the same process of *A. reptans* and were then confirmed via phylogenetic relationships.

### Phylogenetic trees

Reference sequences used in all protein phylogenies were obtained from GenBank (Table S2). Full-length peptide sequences were used. Multiple sequence alignments were generated using ClustalOmega (version 1.2.4; default parameters) and phylogenetic trees were generated by RAxML (version 8.2.12; Model = protgammaauto; Algorithm = a) with support from 1000 bootstrap replicates (Sievers *et al*., 2011; Stamatakis, 2014). Tree graphics were rendered using the Interactive Tree of Life (version 6.5.2) (Letunic and Bork, 2024).

### Plant material and cloning

Plants were obtained from commercial nurseries or botanical gardens (Table S4) and grown in a greenhouse under ambient photoperiod and 24 °C day/17 °C night temperatures.

Synthetic oligonucleotides for all enzymes used in this study are given in Table S5. Candidate enzymes were PCR-amplified from leaf cDNA, and coding sequences were cloned and sequence-verified with respective gene models. Constructs were then cloned into the plant expression vector pEAQ-HT and used in transient expression assays in *N. benthamiana*. One construct, VacCYP76BK1, was synthesized by Twist Bioscience before cloning into pEAQ-HT.

### Transcriptomic sequencing and assembly of **T. chamaedrys**

100 mg young leaf tissue of young leaves were harvested and frozen in liquid nitrogen. RNA was isolated using Spectrum Plant Total RNA kit (Sigma) with on-column DNAse digest. TruSeq stranded mRNA (polyA mRNA) libraries were constructed and sequenced on an Illumina Novaseq 6000 to 150 nt in paired-end mode. Sequencing was performed at the Research Technology Support Facility at Michigan State University.

Raw reads were evaluated with fastqc (v0.11.2) Babraham Bioinformatics), trimmed and corrected using trimmomatic (v0.39) (Bolger *et al*., 2014) and subsequently assembled using trinity (v 2.1.1) (Grabherr *et al*., 2011). Peptide sequences were predicted using Transdecoder (v. 5.5.0) Haas, BJ. https://github.com/TransDecoder/TransDecoder). The predicted peptides were blasted against *CamCYP76BK1*, *ArCYP76BK1*, and *VacCYP76BK1*. *TchCYP76BK1* was found to be 94.7% identical to *TcaCYP76BK1* identified and included in producing the final phylogenies and for cloning

### Transient expression for functional characterization in N. benthamiana

*N. benthamiana* plants were grown for 5 weeks in a controlled growth room under 12H light and 12H dark cycle at (22°C) before infiltration. Constructs for co-expression were separately transformed into *Agrobacterium tumefaciens* strain LBA4404. 20 mL cultures were grown overnight at 30□°C in LB with 50□µg/mL kanamycin and 50□µg/mL rifampicin. Cultures were collected by centrifugation and washed twice with 10□mL water. Cells were resuspended and diluted to an OD_600_ of 1.0 in 200 µM acetosyringone/water and incubated at 30□°C for 1–2 H. Separate cultures were mixed in a 1:1 ratio for each combination of enzymes, and 4- or 5-week-old plants were infiltrated with a 1□mL syringe into the underside (abaxial side) of *N. benthamiana* leaves. All gene constructs were co-infiltrated with two genes encoding rate-limiting steps in the upstream (MEP) pathway: *P. barbatus* 1-deoxy-D-xylulose-5-phosphate synthase (*PbDXS*) and GGPP synthase (*PbGGPPS*) to boost production of the diterpene precursor GGPP (Andersen-Ranberg *et al*., 2016). Plants were returned to the controlled growth room (22□°C, 12 H diurnal cycle) for 5 days. Approximately 200□mg fresh weight from infiltrated leaves was extracted with 1□mL hexane overnight at room temperature. Plant material was collected by centrifugation, and the organic phase was removed for GC-MS analysis.

### Plant extract metabolomics

Leaves from *C. pyramidata*, *P. bambusetorum*, *H. sanguinea*, *S. baicalensis*, *C. bungei*, *T. chamaedrys, A. reptans,* and *C. americana* were harvested for metabolite analysis. The leaves were frozen in liquid nitrogen, crushed, and extracted for three hours in ethyl acetate. Leaf material was collected by centrifugation and the organic phase was removed and concentrated for GC-MS analysis.

### GC-MS analysis

All GC-MS analyses were performed in Michigan State University’s Mass Spectrometry and Metabolomics Core Facility on an Agilent 7890⍰A GC with an Agilent VF-5ms column (30⍰m × 250⍰µm × 0.25⍰µm, with 10⍰m EZ-Guard) and an Agilent 5975⍰C detector. The inlet was set to 250⍰°C splitless injection of 1⍰µL and He carrier gas (1⍰mL/min), and the detector was activated following a 4⍰min solvent delay. All assays and tissue analysis used the following method: temperature ramp start 40⍰°C, hold 1⍰min, 40⍰°C/min to 200⍰°C, hold 4.5⍰min, 20⍰°C/min to 240⍰°C, 10⍰°C/min to 280⍰°C, 40⍰°C/min to 320⍰°C, and hold 5⍰min. MS scan range was set to 40–400. Deconvolution of spectra was done utilizing AMDIS (NIST. 2019).

### Product scale-up and NMR

For NMR analysis, production in the *N. benthamiana* system was scaled up to 1 L infection culture. A vacuum-infiltration system was used to infiltrate *A. tumefaciens* strains into whole *N. benthamiana* plants, with approximately 40 plants used for each enzyme combination. The furan and lactone derivatives of CamTPS2 were identified from the combination of CamTPS2 and CbCYP76BK1. The furan derivative of ArTPS2 was identified from the combination of ArTPS2 and ArCYP76BK1, while the C16 lactone derivative was identified from ArTPS2 with HsCYP76BK1 and the C15 lactone derivative utilized SbarbCYP76BK1. After 5 days, all leaf tissue was harvested and extracted overnight in 600□mL hexane at room temperature. The extract was concentrated by rotary evaporator. Each product was purified by silica gel flash column chromatography with a mobile phase of 98% hexane/2% ethyl acetate. NMR spectra were measured in Michigan State University’s Max T. Rogers NMR Facility on a Bruker Avance NEO 800 MHz or 600 MHz spectrometer equipped with a helium cooled TCl cryoprobe or a Prodigy TCI cryoprobe, respectively, using CDCl_3_ as the solvent. CDCl_3_ peaks were referenced to 7.26 and 77.00 ppm for ^1^H and ^13^C spectra, respectively.

### Syntenic Analysis of Lamiaceae

Publicly available chromosome-scale genomes of *C. armericana* (Hamilton *et al*., 2020), *Cleorodendrum inerme* (He *et al*., 2022), *Pogostemon cablin* (Shen *et al*., 2022), *Salvia hispanica* (Brose *et al*., 2024)*, Salvia miltiorrhiza* (Pan *et al*., 2023b), *Salvia officinalis* (Li *et al*., 2022), *Salvia splendens* (Dong *et al*., 2018), *Scutellaria baicalensis* (Q., Zhao *et al*., 2019), *Scutellaria barbata* Xu *et al*., 2020), *Tectona grandis* (D., Zhao *et al*., 2019), *Thymus quinquecostatus* (Sun *et al*., 2022), and *Lavandula angustifolia* (Hamilton *et al*., 2023) were obtained and quality assessed using BUSCO (v5.5.0; (Simao *et al*., 2015) embryophyta_odb10. In order to identify syntenic orthologs, only chromosome-scale assemblies with BUSCO (Basic Universal Single Copy Orthologs) scores for the genome greater than 90% and annotation scores greater than 80% (Table S6). Syntelogs through the Lamiaceae were obtained for the chromosome scale assemblies within the Lamiaceae with GENESPACE (v.1.1.10; (Lovell *et al*., 2022). The regions were then visualized using pyGenomeViz (Shimoyama, 2022). The phylogeny of the Lamiaceae species was generated by (Brose *et al*., 2024). The final figure was edited in BioRender.

## Accession numbers

Relevant accession numbers can be found on Table S1 and Table S3

## Data availability Statement

The raw sequence reads for the *Teucrium chamaedrys* transcriptome are available in the National Center for Biotechnology Information Sequence Read Archive under BioProject PRJNA1124528.

## Author contributions

NS, EL, TA and BH conceived the study. NS and EL wrote the manuscript with contributions from JB and TA. Pathways and constructs were designed by NS and EL. Terpene analysis in transient assays was performed by NS and EL. JB performed all genomic and syntenic analysis. DH assisted critically with NMR analysis.

## Acknowledgements

We would like to thank Britta Hamberger for assistance in maintaining plant material. P450 annotation was kindly provided by David Nelson (University of Tennessee). We would like to thank Drs. Cassandra Johnny and Anthony Schilmiller of Michigan State University’s Mass Spectrometry and Metabolomics Core Facility for their help in obtaining and interpreting GC-MS data, and the Max T. Rogers NMR Facility for their help in obtaining NMR data. We would also like to thank Dr. David Nelson for naming all CYP sequences presented in this work.

We collectively acknowledge that Michigan State University occupies the ancestral, traditional, and contemporary Lands of the Anishinaabeg – Three Fires Confederacy of Ojibwe, Odawa, and Potawatomi peoples. In particular, the University resides on Land ceded in the 1819 Treaty of Saginaw. We recognize, support, and advocate for the sovereignty of Michigan’s twelve federally-recognized Indian nations, for historic Indigenous communities in Michigan, for Indigenous individuals and communities who live here now, and for those who were forcibly removed from their Homelands. By offering this Land Acknowledgement, we affirm Indigenous sovereignty and will work to hold Michigan State University more accountable to the needs of American Indian and Indigenous peoples.

## Short Legend Titles for SI

- Table S1: Reference Sequences for phylogenies
- Table S2. List of mint transcriptomes analyzed along with species abbreviations and subfamily.
- Table S2. List of mint transcriptomes analyzed along with species abbreviations and subfamily.
- Table S3. GenBank accession numbers for genes cloned in this study.
- Table S4: Plant material sources
- Table S5. List primers used in this study.
- Table S6. Genomes used in Syntenic Analysis and BUSCO^a^ scores
- **Figure S1. P450 candidates selected for functional characterization**
- Figure S2: *C. americana* and *A. reptans* CYP76BK1 ortholog activity.
- Figure S3: NMR chemical shift assignments of compound 1
- Figure S4: NMR spectra of Compound 1
- Figure S5: NMR chemical shift assignments of compound 2
- Figure S6: NMR spectra of Compound 2
- Figure S7: CYP76 family phylogeny from the Mint Genome Project
- Figure S8: NMR chemical shift assignments of compound 3
- Figure S9: NMR spectra of 3
- Figure S10: NMR chemical shift assignments of compound 4
- Figure S11: NMR spectra of compound 4
- Figure S12: NMR chemical shift assignments of compound 5
- Figure S13: NMR spectra of compound 5
- Figure S14: kolavenol CYP76BK1 products
- Figure S15: isokolavenol CYP76BK1 products
- Figure S16: Extracted ion chromatograms of peregrinol backbone paired with CYP76BKs
- Figure S17: GCMS analysis of plant extracts
- Figure S18: Lamiaceae clerodane and labdane furan moiety distribution
- Figure S19: Combined kolavenol and isokolavenol CYP76BK1 total ion chromatograms.
- Figure S20: Lamiaceae diterpenoids by backbone type
- Figure S21:Extracted ion chromatograms (286) of ArCYP76BK1 against ent and (+)-copalyl diphosphate synthases

