## Supplemental Figures for "CYP76BK1 orthologs catalyze furan and lactone ring formation in clerodane diterpenoids across the mint family"

### Slide 1
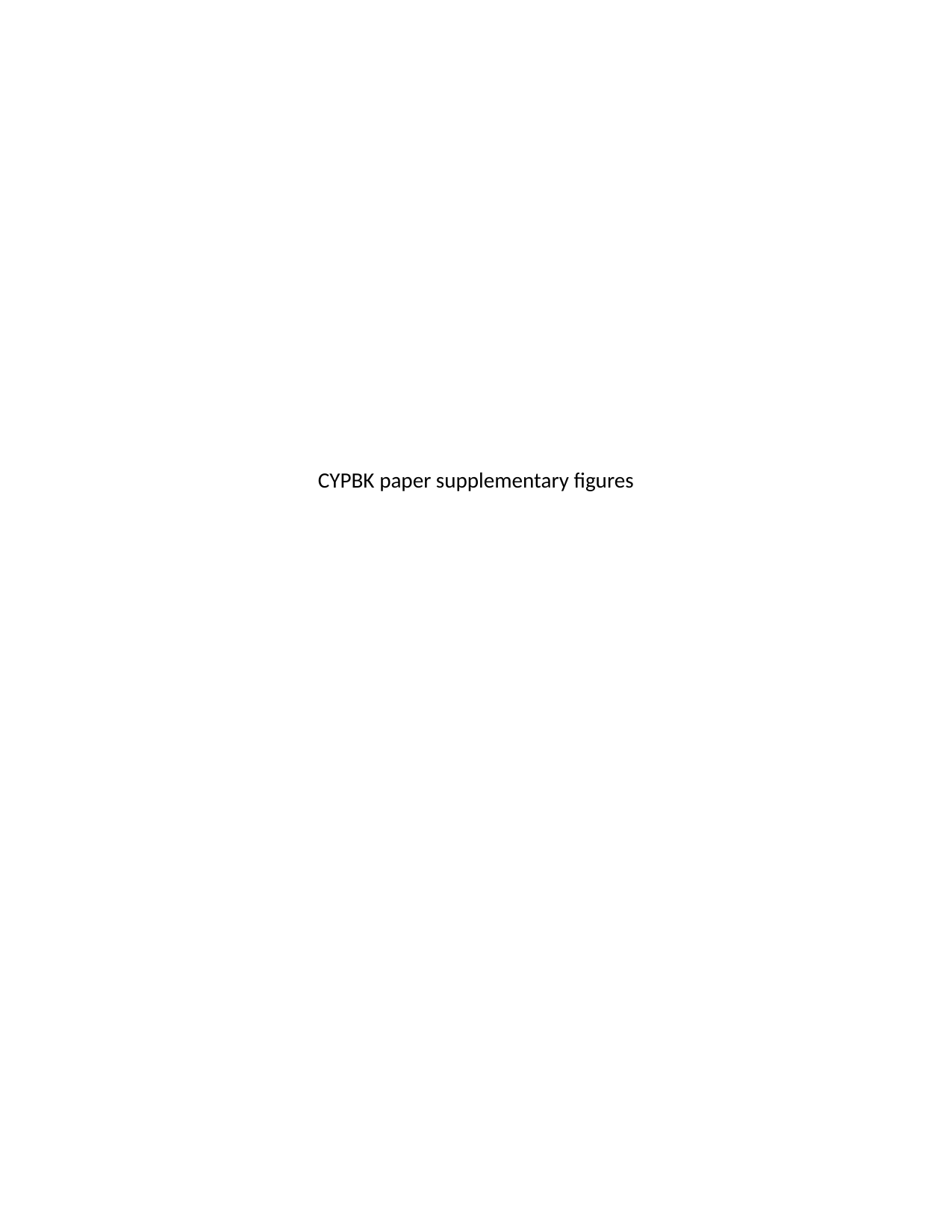

CYPBK paper supplementary figures

### Slide 2
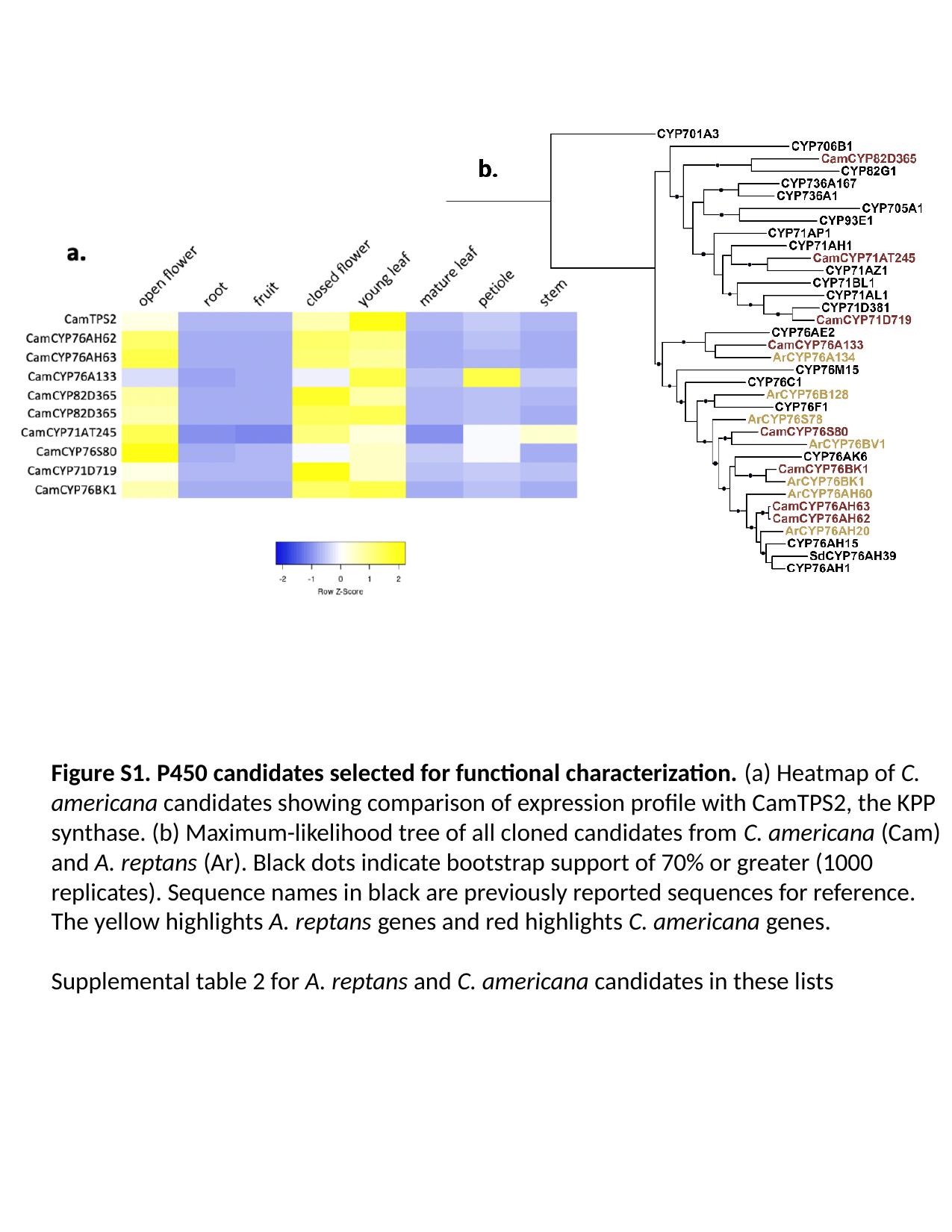

Figure S1. P450 candidates selected for functional characterization. (a) Heatmap of C. americana candidates showing comparison of expression profile with CamTPS2, the KPP synthase. (b) Maximum-likelihood tree of all cloned candidates from C. americana (Cam) and A. reptans (Ar). Black dots indicate bootstrap support of 70% or greater (1000 replicates). Sequence names in black are previously reported sequences for reference. The yellow highlights A. reptans genes and red highlights C. americana genes.
Supplemental table 2 for A. reptans and C. americana candidates in these lists

### Slide 3
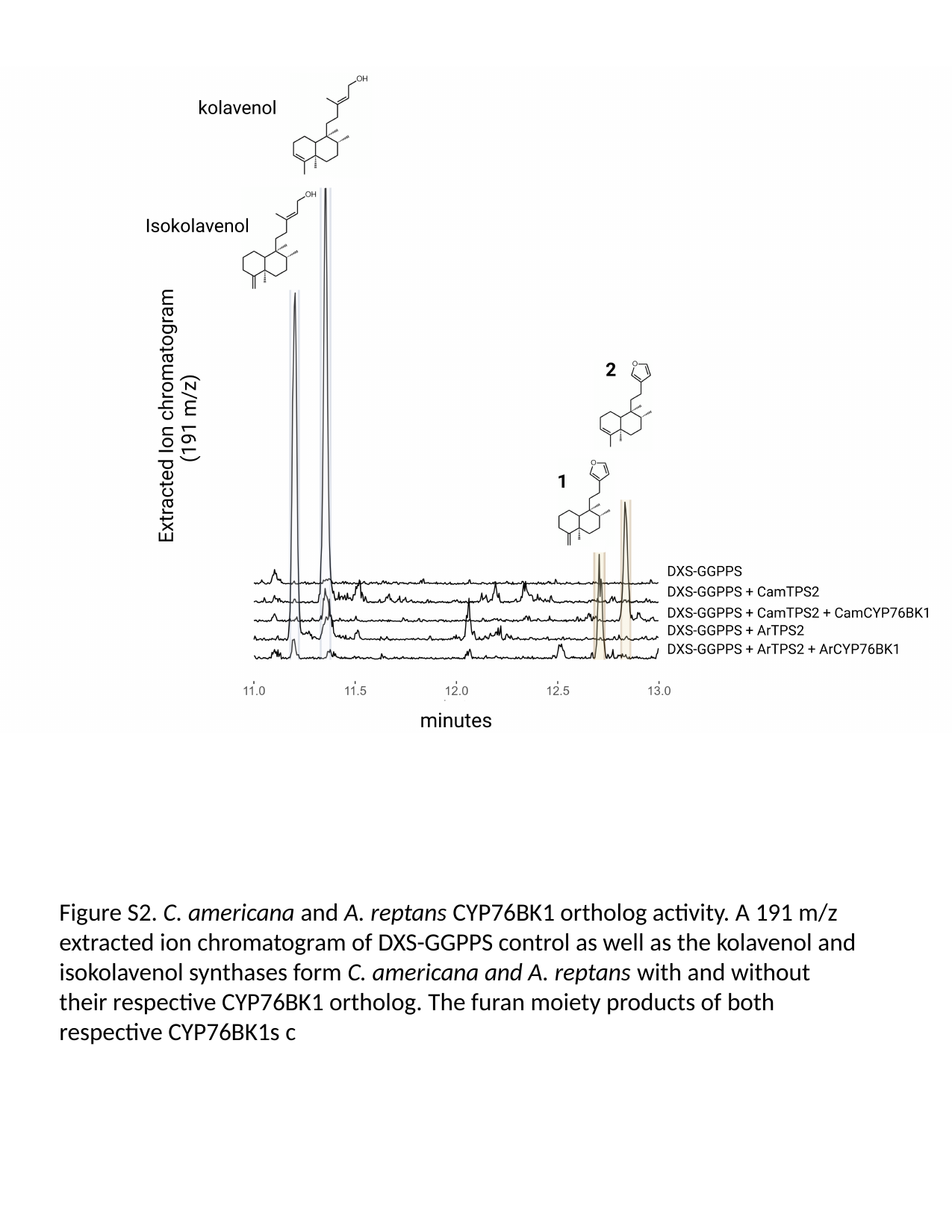

Figure S2. C. americana and A. reptans CYP76BK1 ortholog activity. A 191 m/z extracted ion chromatogram of DXS-GGPPS control as well as the kolavenol and isokolavenol synthases form C. americana and A. reptans with and without their respective CYP76BK1 ortholog. The furan moiety products of both respective CYP76BK1s c

### Slide 4
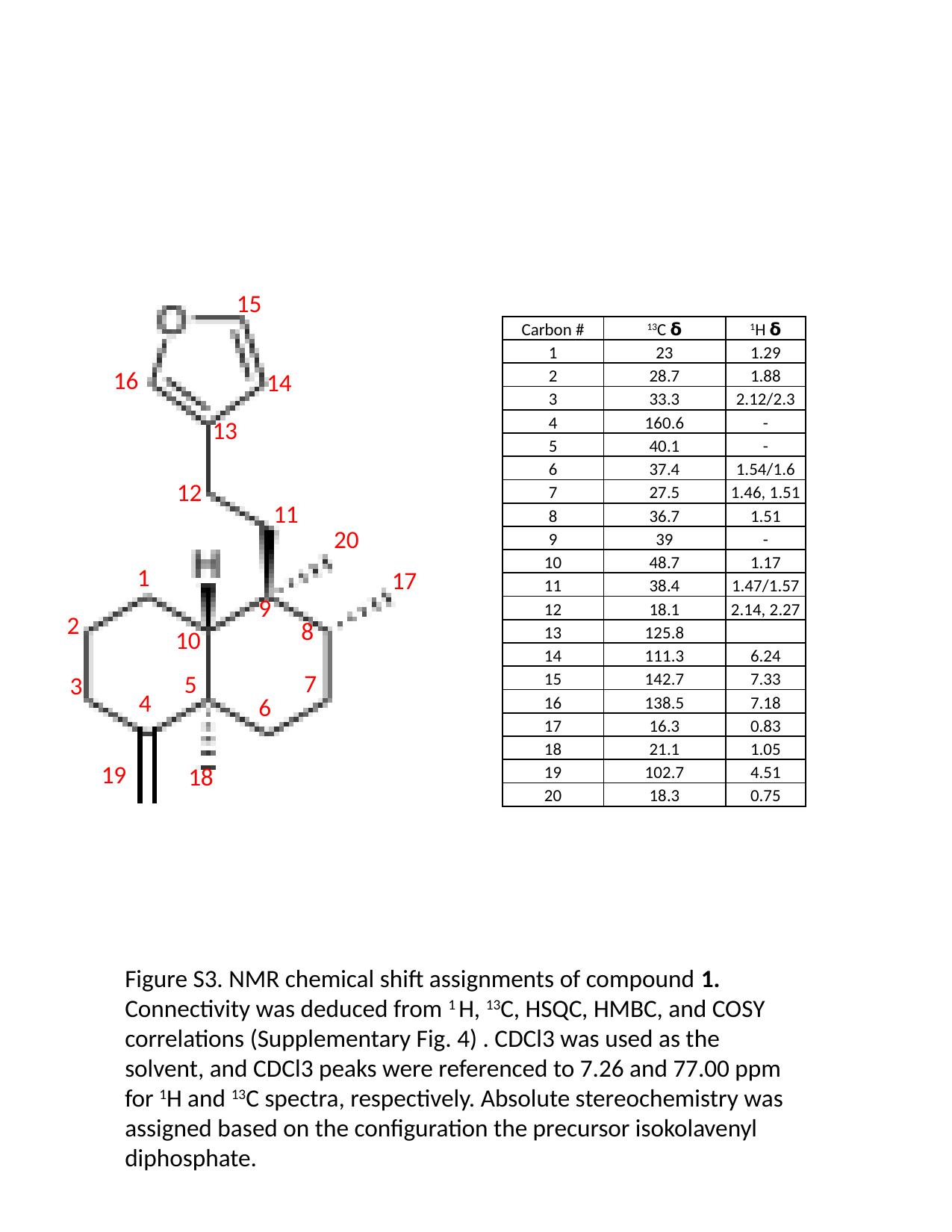

15
16
14
13
12
11
20
1
17
9
2
8
10
7
5
3
4
6
19
18
| Carbon # | ^13^C 𝝳 | ^1^H 𝝳 |
| --- | --- | --- |
| 1 | 23 | 1.29 |
| 2 | 28.7 | 1.88 |
| 3 | 33.3 | 2.12/2.3 |
| 4 | 160.6 | - |
| 5 | 40.1 | - |
| 6 | 37.4 | 1.54/1.6 |
| 7 | 27.5 | 1.46, 1.51 |
| 8 | 36.7 | 1.51 |
| 9 | 39 | - |
| 10 | 48.7 | 1.17 |
| 11 | 38.4 | 1.47/1.57 |
| 12 | 18.1 | 2.14, 2.27 |
| 13 | 125.8 | |
| 14 | 111.3 | 6.24 |
| 15 | 142.7 | 7.33 |
| 16 | 138.5 | 7.18 |
| 17 | 16.3 | 0.83 |
| 18 | 21.1 | 1.05 |
| 19 | 102.7 | 4.51 |
| 20 | 18.3 | 0.75 |
Figure S3. NMR chemical shift assignments of compound 1. Connectivity was deduced from ^1^ H, ^13^C, HSQC, HMBC, and COSY correlations (Supplementary Fig. 4) . CDCl3 was used as the solvent, and CDCl3 peaks were referenced to 7.26 and 77.00 ppm for ^1^H and ^13^C spectra, respectively. Absolute stereochemistry was assigned based on the configuration the precursor isokolavenyl diphosphate.

### Slide 5
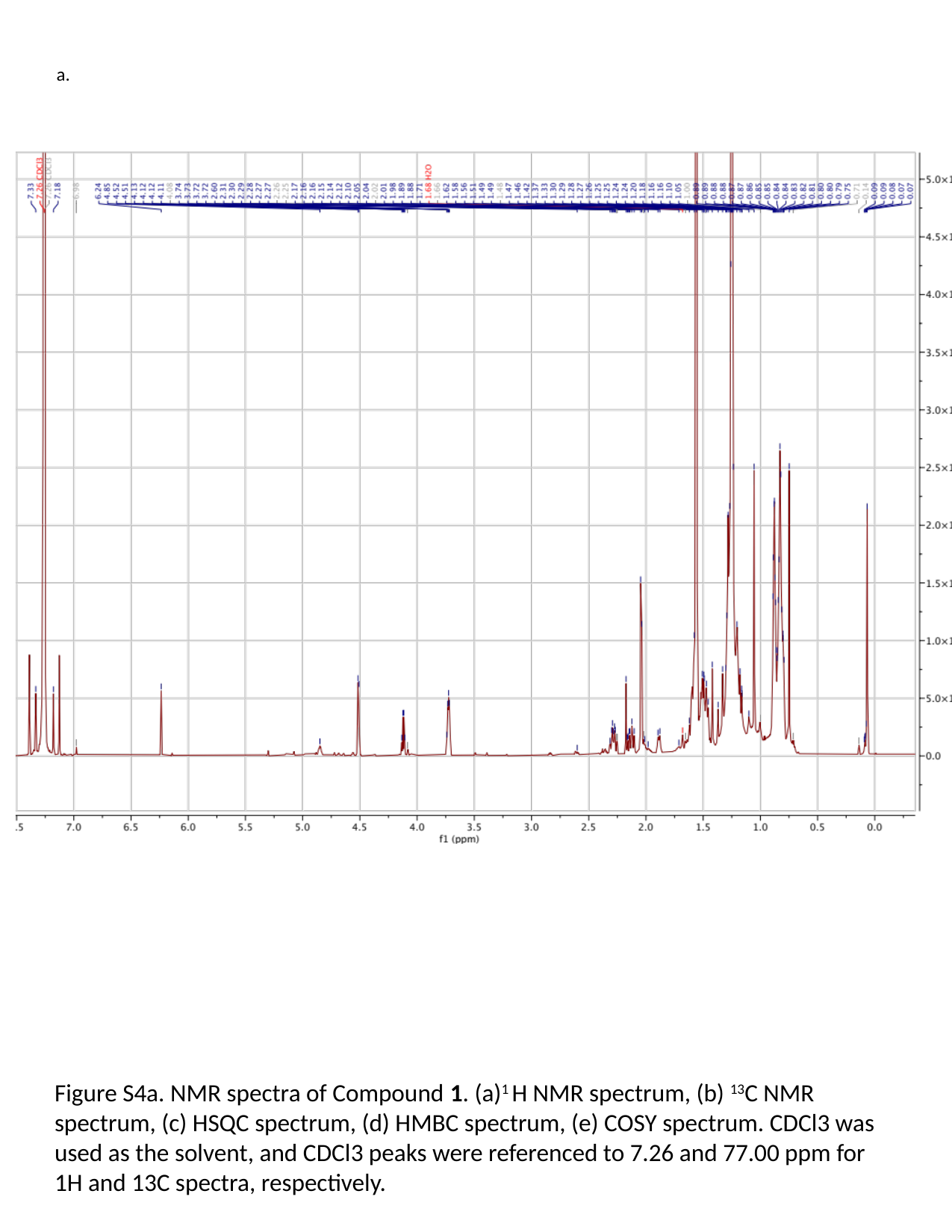

a.
Figure S4a. NMR spectra of Compound 1. (a)^1^ H NMR spectrum, (b) ^13^C NMR spectrum, (c) HSQC spectrum, (d) HMBC spectrum, (e) COSY spectrum. CDCl3 was used as the solvent, and CDCl3 peaks were referenced to 7.26 and 77.00 ppm for 1H and 13C spectra, respectively.

### Slide 6
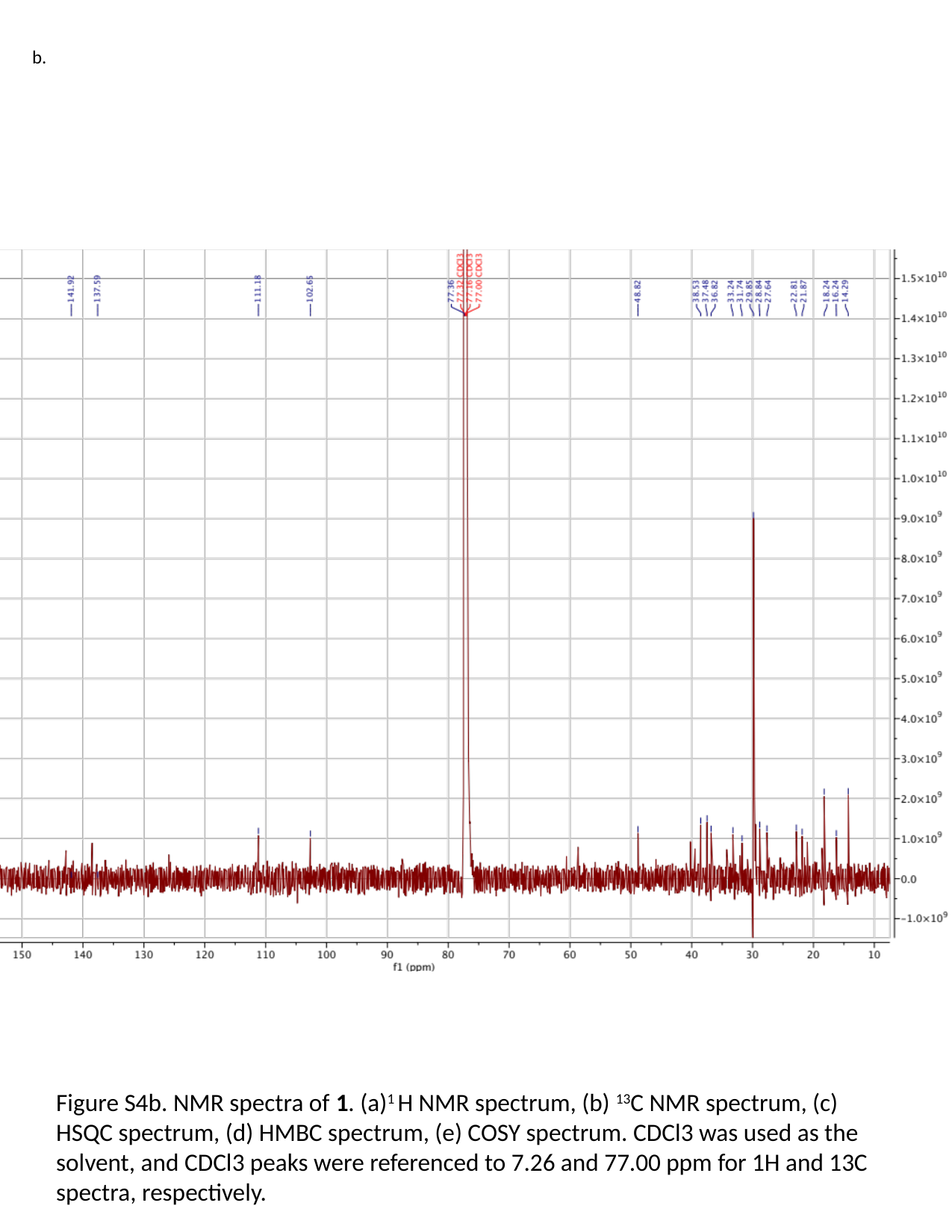

b.
Figure S4b. NMR spectra of 1. (a)^1^ H NMR spectrum, (b) ^13^C NMR spectrum, (c) HSQC spectrum, (d) HMBC spectrum, (e) COSY spectrum. CDCl3 was used as the solvent, and CDCl3 peaks were referenced to 7.26 and 77.00 ppm for 1H and 13C spectra, respectively.

### Slide 7
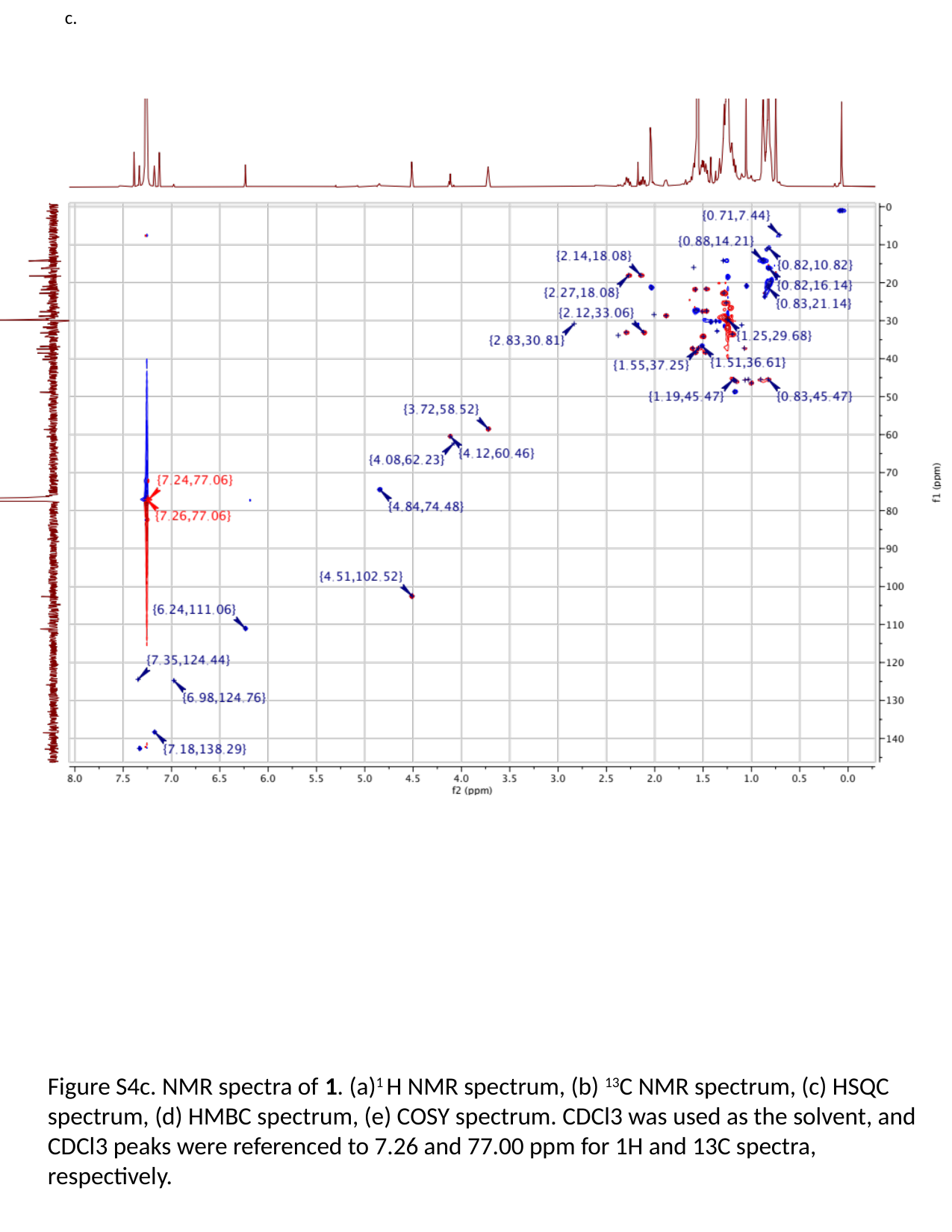

c.
Figure S4c. NMR spectra of 1. (a)^1^ H NMR spectrum, (b) ^13^C NMR spectrum, (c) HSQC spectrum, (d) HMBC spectrum, (e) COSY spectrum. CDCl3 was used as the solvent, and CDCl3 peaks were referenced to 7.26 and 77.00 ppm for 1H and 13C spectra, respectively.

### Slide 8
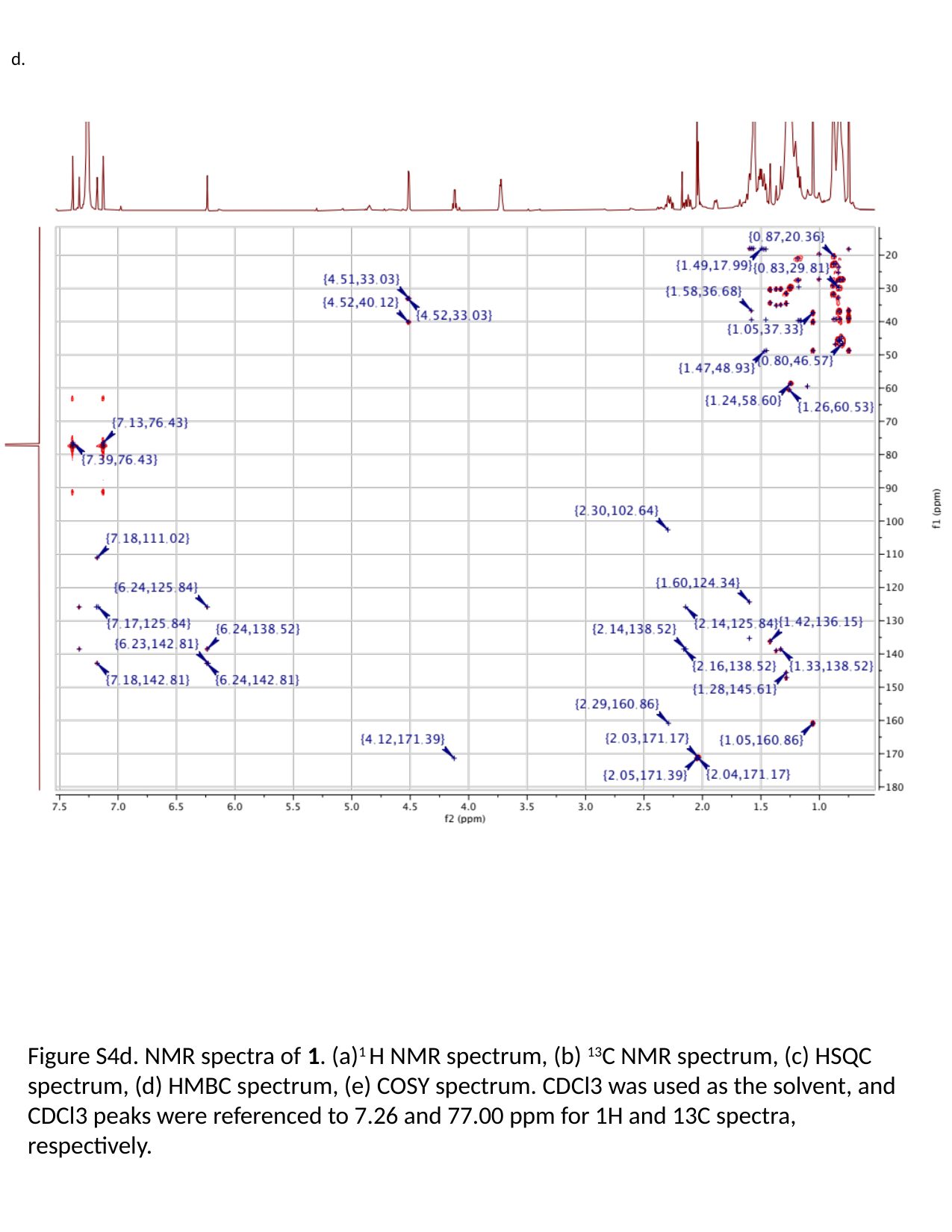

d.
Figure S4d. NMR spectra of 1. (a)^1^ H NMR spectrum, (b) ^13^C NMR spectrum, (c) HSQC spectrum, (d) HMBC spectrum, (e) COSY spectrum. CDCl3 was used as the solvent, and CDCl3 peaks were referenced to 7.26 and 77.00 ppm for 1H and 13C spectra, respectively.

### Slide 9
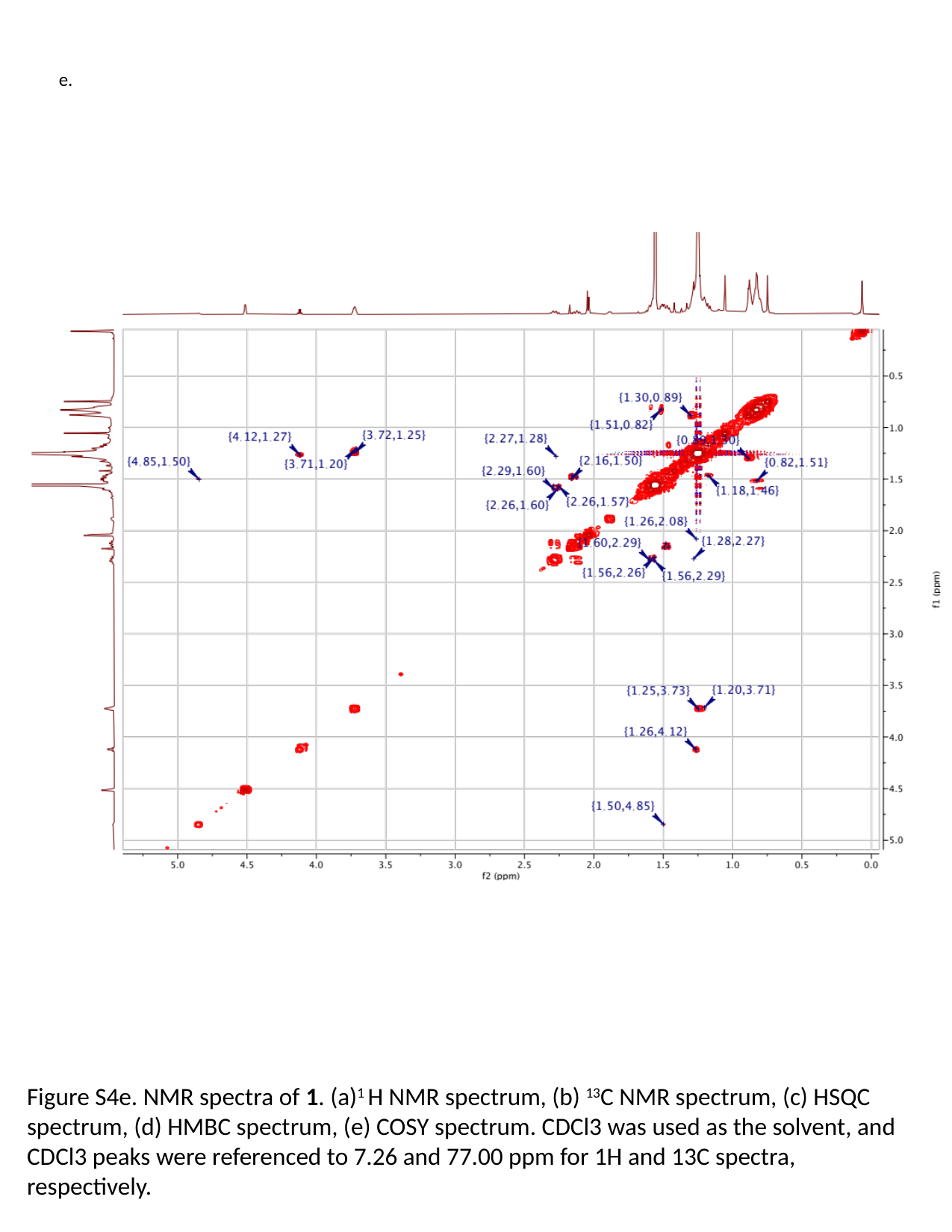

e.
Figure S4e. NMR spectra of 1. (a)^1^ H NMR spectrum, (b) ^13^C NMR spectrum, (c) HSQC spectrum, (d) HMBC spectrum, (e) COSY spectrum. CDCl3 was used as the solvent, and CDCl3 peaks were referenced to 7.26 and 77.00 ppm for 1H and 13C spectra, respectively.

### Slide 10
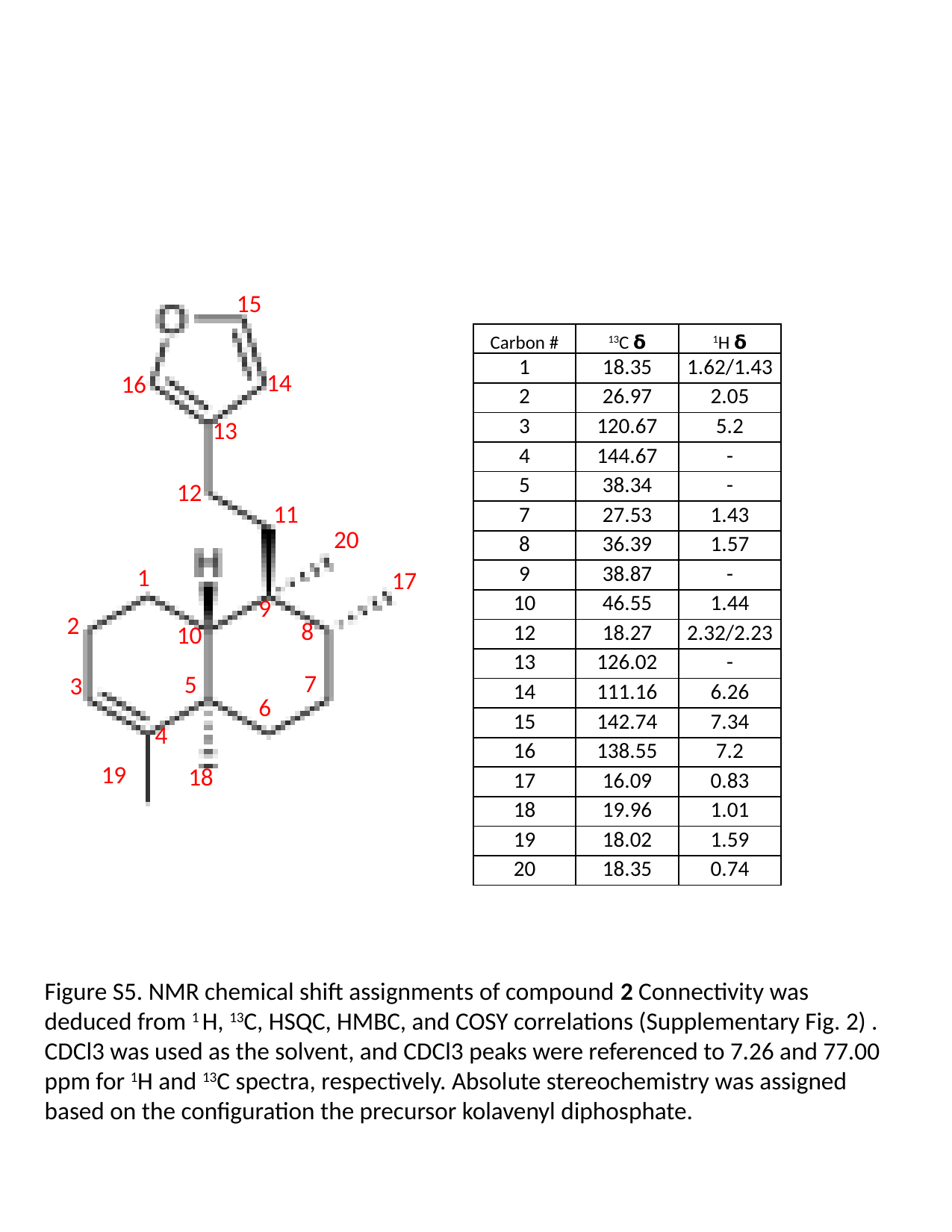

15
14
16
13
12
11
20
1
17
9
2
8
10
7
5
3
6
4
19
18
| Carbon # | ^13^C 𝝳 | ^1^H 𝝳 |
| --- | --- | --- |
| 1 | 18.35 | 1.62/1.43 |
| 2 | 26.97 | 2.05 |
| 3 | 120.67 | 5.2 |
| 4 | 144.67 | - |
| 5 | 38.34 | - |
| 7 | 27.53 | 1.43 |
| 8 | 36.39 | 1.57 |
| 9 | 38.87 | - |
| 10 | 46.55 | 1.44 |
| 12 | 18.27 | 2.32/2.23 |
| 13 | 126.02 | - |
| 14 | 111.16 | 6.26 |
| 15 | 142.74 | 7.34 |
| 16 | 138.55 | 7.2 |
| 17 | 16.09 | 0.83 |
| 18 | 19.96 | 1.01 |
| 19 | 18.02 | 1.59 |
| 20 | 18.35 | 0.74 |
Figure S5. NMR chemical shift assignments of compound 2 Connectivity was deduced from ^1^ H, ^13^C, HSQC, HMBC, and COSY correlations (Supplementary Fig. 2) . CDCl3 was used as the solvent, and CDCl3 peaks were referenced to 7.26 and 77.00 ppm for ^1^H and ^13^C spectra, respectively. Absolute stereochemistry was assigned based on the configuration the precursor kolavenyl diphosphate.

### Slide 11
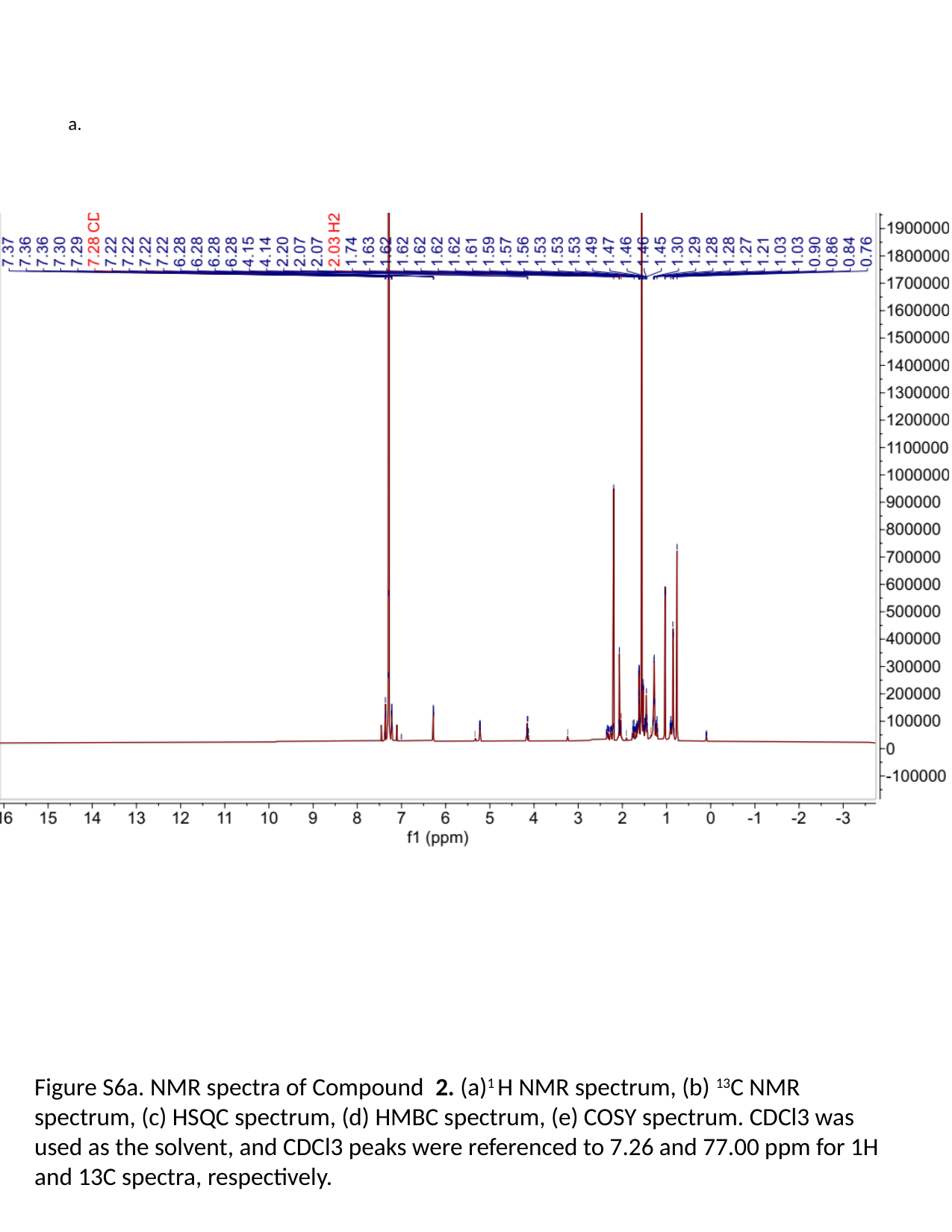

a.
Figure S6a. NMR spectra of Compound 2. (a)^1^ H NMR spectrum, (b) ^13^C NMR spectrum, (c) HSQC spectrum, (d) HMBC spectrum, (e) COSY spectrum. CDCl3 was used as the solvent, and CDCl3 peaks were referenced to 7.26 and 77.00 ppm for 1H and 13C spectra, respectively.

### Slide 12
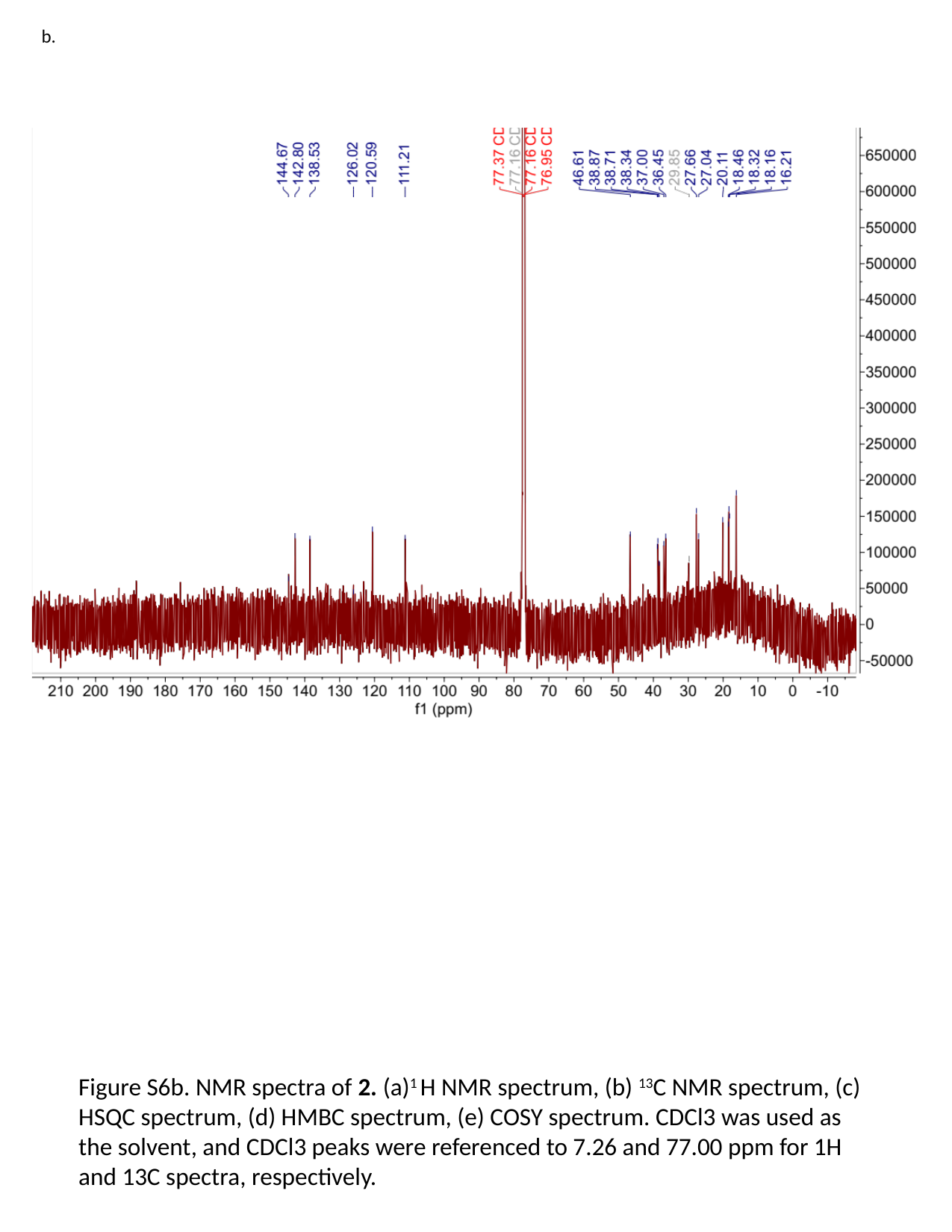

b.
Figure S6b. NMR spectra of 2. (a)^1^ H NMR spectrum, (b) ^13^C NMR spectrum, (c) HSQC spectrum, (d) HMBC spectrum, (e) COSY spectrum. CDCl3 was used as the solvent, and CDCl3 peaks were referenced to 7.26 and 77.00 ppm for 1H and 13C spectra, respectively.

### Slide 13
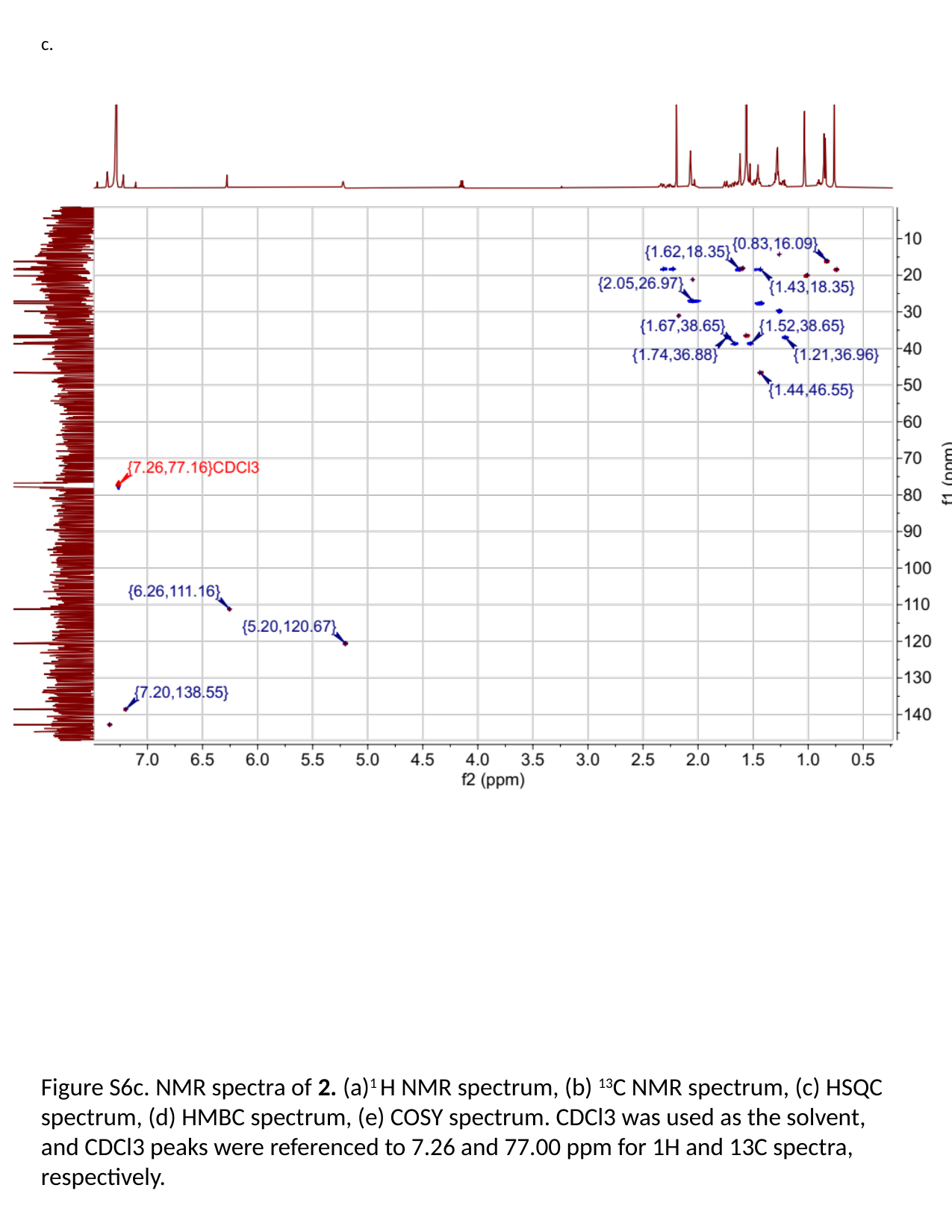

c.
Figure S6c. NMR spectra of 2. (a)^1^ H NMR spectrum, (b) ^13^C NMR spectrum, (c) HSQC spectrum, (d) HMBC spectrum, (e) COSY spectrum. CDCl3 was used as the solvent, and CDCl3 peaks were referenced to 7.26 and 77.00 ppm for 1H and 13C spectra, respectively.

### Slide 14
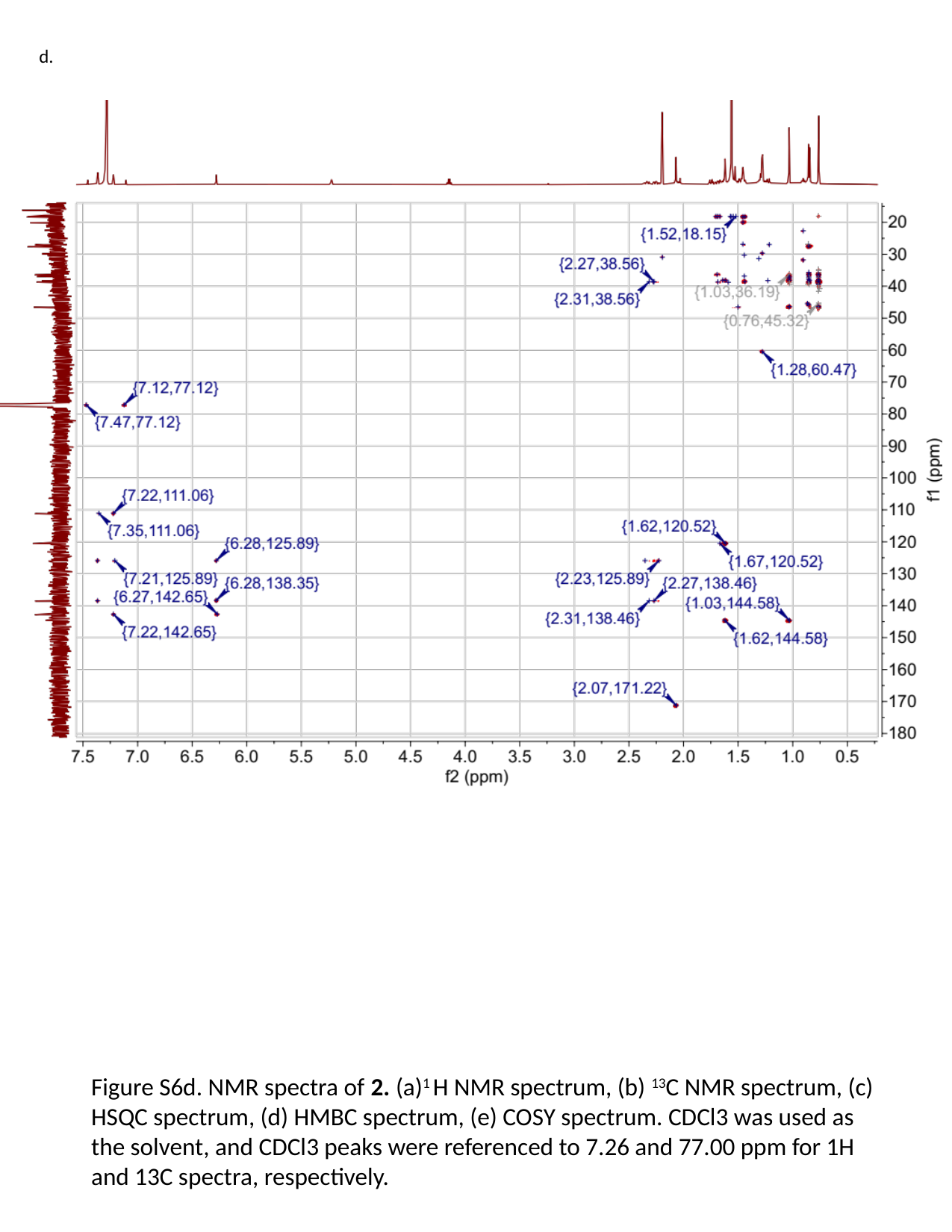

d.
Figure S6d. NMR spectra of 2. (a)^1^ H NMR spectrum, (b) ^13^C NMR spectrum, (c) HSQC spectrum, (d) HMBC spectrum, (e) COSY spectrum. CDCl3 was used as the solvent, and CDCl3 peaks were referenced to 7.26 and 77.00 ppm for 1H and 13C spectra, respectively.

### Slide 15
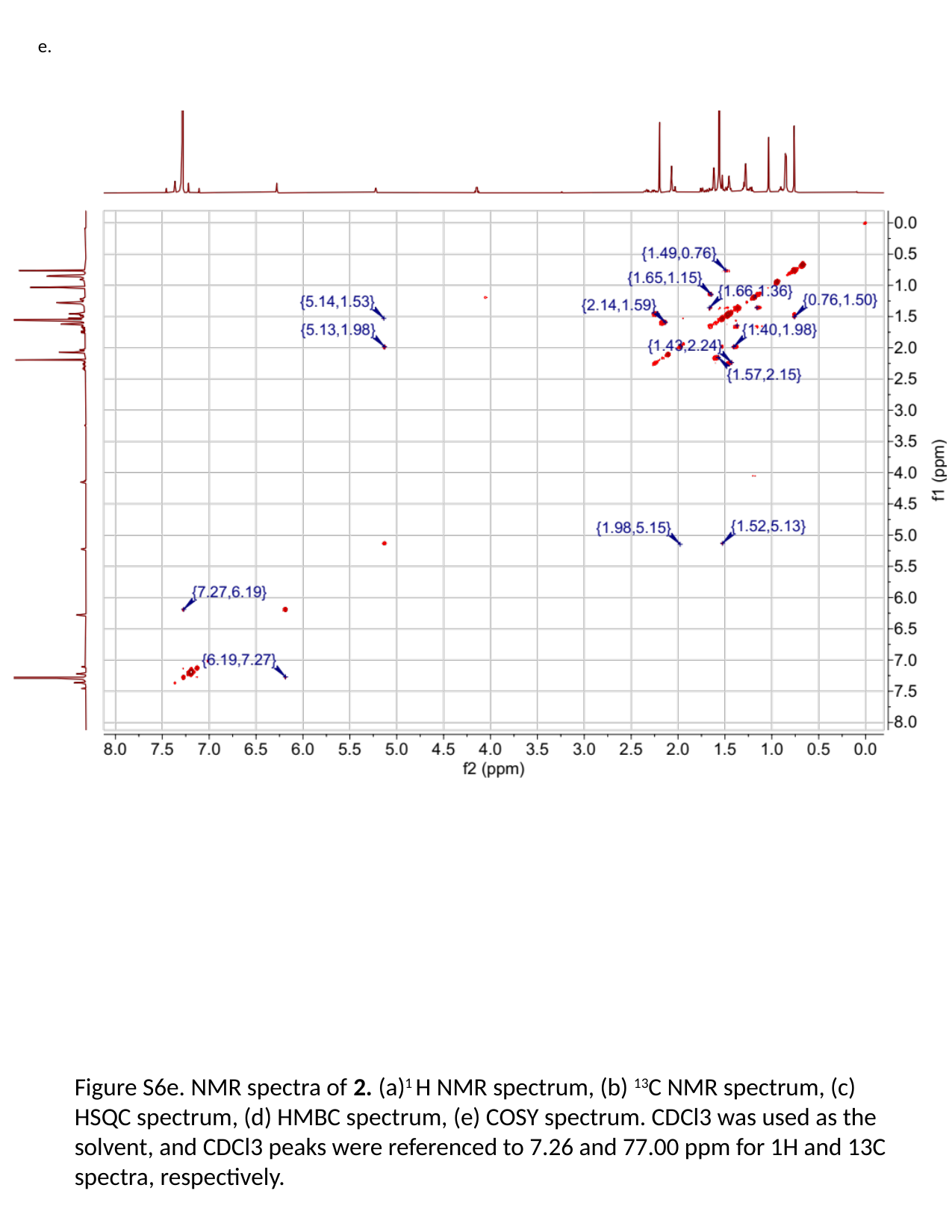

e.
Figure S6e. NMR spectra of 2. (a)^1^ H NMR spectrum, (b) ^13^C NMR spectrum, (c) HSQC spectrum, (d) HMBC spectrum, (e) COSY spectrum. CDCl3 was used as the solvent, and CDCl3 peaks were referenced to 7.26 and 77.00 ppm for 1H and 13C spectra, respectively.

### Slide 16
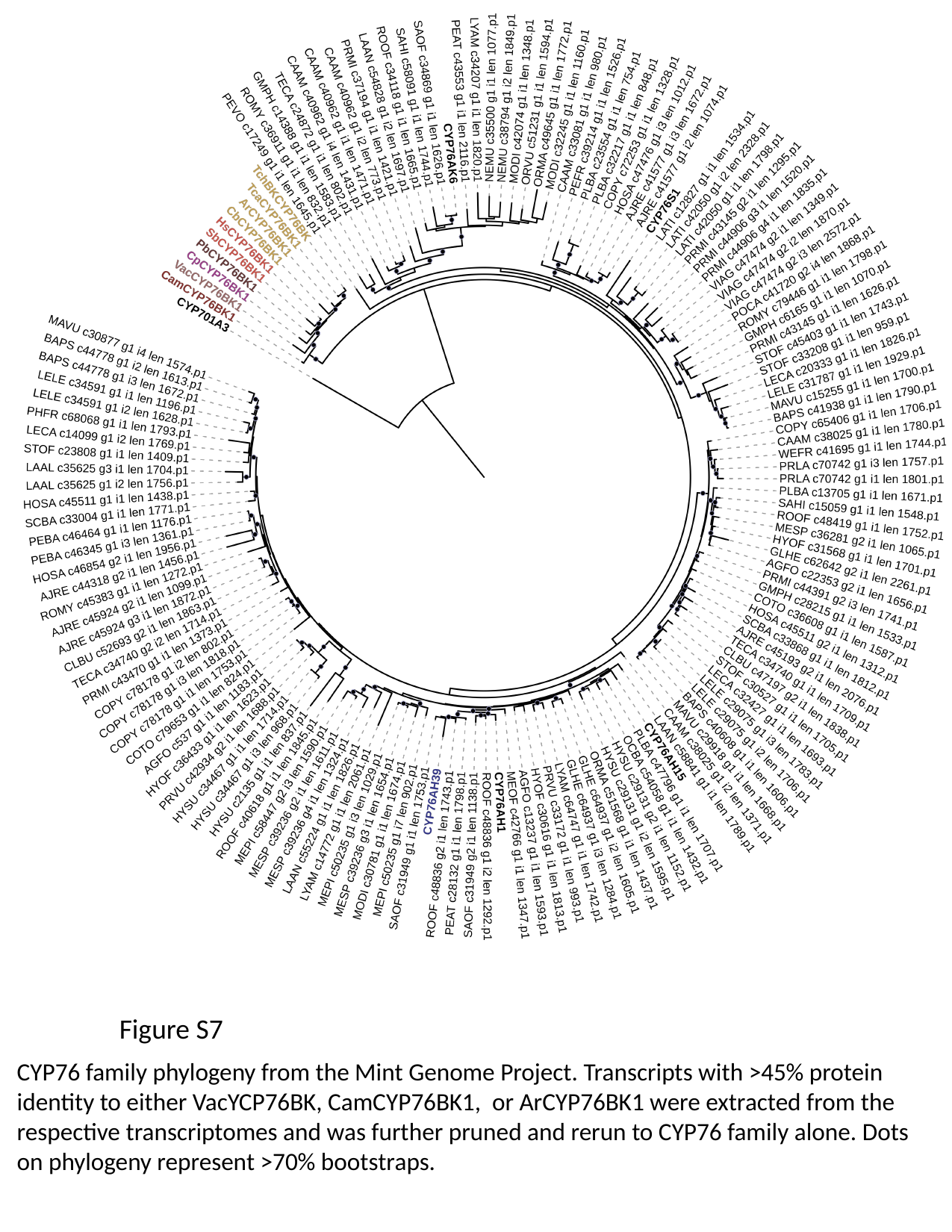

Figure S7
CYP76 family phylogeny from the Mint Genome Project. Transcripts with >45% protein identity to either VacYCP76BK, CamCYP76BK1, or ArCYP76BK1 were extracted from the respective transcriptomes and was further pruned and rerun to CYP76 family alone. Dots on phylogeny represent >70% bootstraps.

### Slide 17
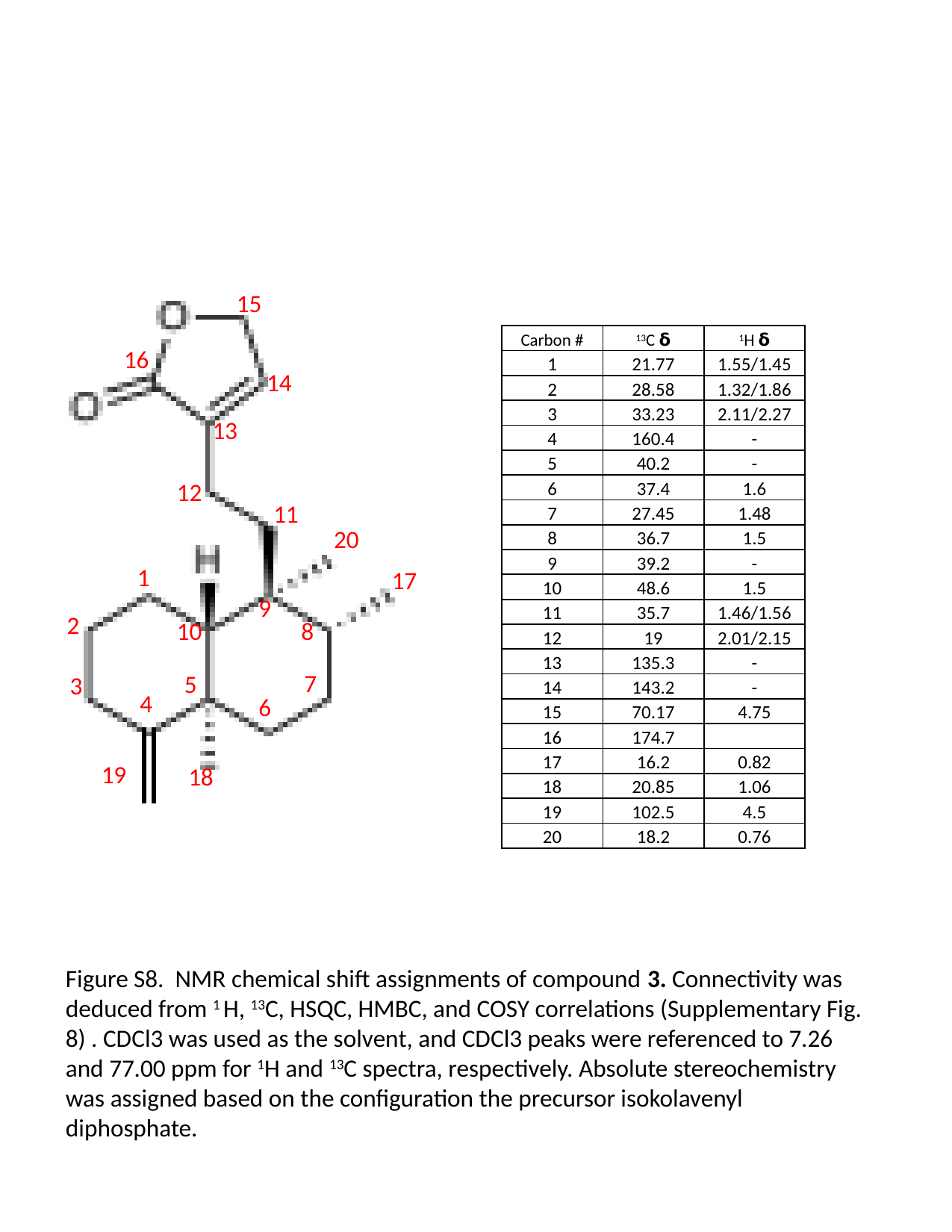

15
16
14
13
12
11
20
1
17
9
2
8
10
7
5
3
4
6
19
18
| Carbon # | ^13^C 𝝳 | ^1^H 𝝳 |
| --- | --- | --- |
| 1 | 21.77 | 1.55/1.45 |
| 2 | 28.58 | 1.32/1.86 |
| 3 | 33.23 | 2.11/2.27 |
| 4 | 160.4 | - |
| 5 | 40.2 | - |
| 6 | 37.4 | 1.6 |
| 7 | 27.45 | 1.48 |
| 8 | 36.7 | 1.5 |
| 9 | 39.2 | - |
| 10 | 48.6 | 1.5 |
| 11 | 35.7 | 1.46/1.56 |
| 12 | 19 | 2.01/2.15 |
| 13 | 135.3 | - |
| 14 | 143.2 | - |
| 15 | 70.17 | 4.75 |
| 16 | 174.7 | |
| 17 | 16.2 | 0.82 |
| 18 | 20.85 | 1.06 |
| 19 | 102.5 | 4.5 |
| 20 | 18.2 | 0.76 |
Figure S8. NMR chemical shift assignments of compound 3. Connectivity was deduced from ^1^ H, ^13^C, HSQC, HMBC, and COSY correlations (Supplementary Fig. 8) . CDCl3 was used as the solvent, and CDCl3 peaks were referenced to 7.26 and 77.00 ppm for ^1^H and ^13^C spectra, respectively. Absolute stereochemistry was assigned based on the configuration the precursor isokolavenyl diphosphate.

### Slide 18
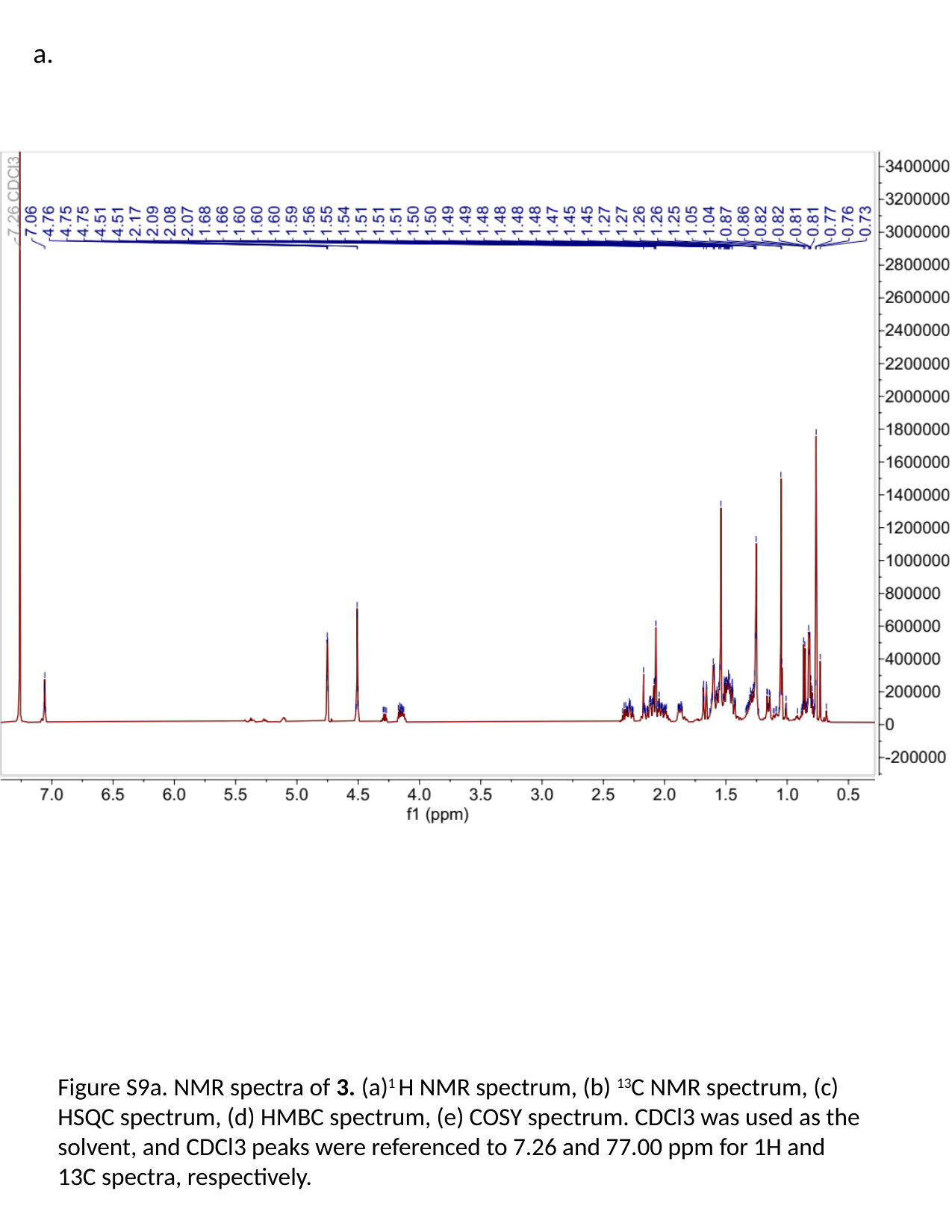

a.
Figure S9a. NMR spectra of 3. (a)^1^ H NMR spectrum, (b) ^13^C NMR spectrum, (c) HSQC spectrum, (d) HMBC spectrum, (e) COSY spectrum. CDCl3 was used as the solvent, and CDCl3 peaks were referenced to 7.26 and 77.00 ppm for 1H and 13C spectra, respectively.

### Slide 19
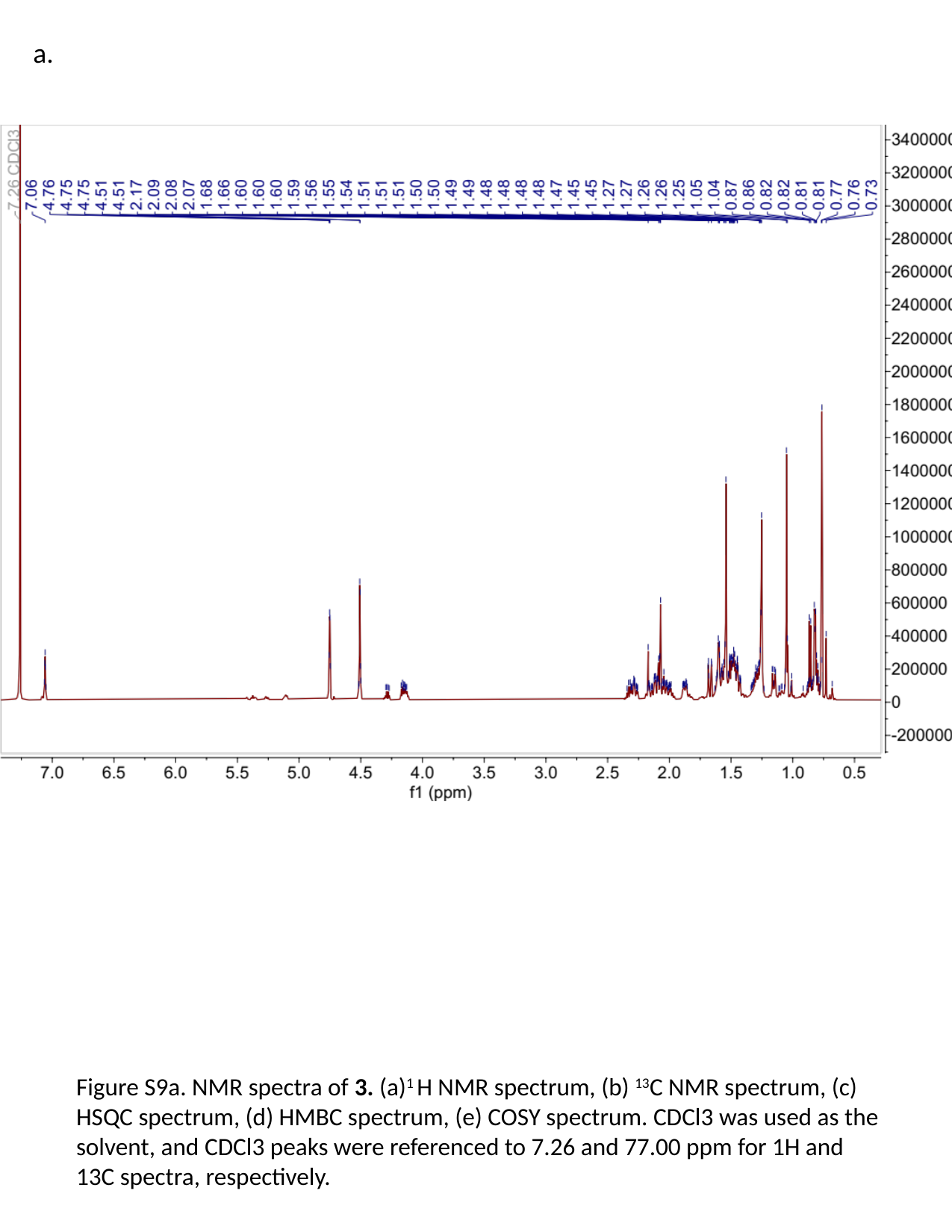

a.
Figure S9a. NMR spectra of 3. (a)^1^ H NMR spectrum, (b) ^13^C NMR spectrum, (c) HSQC spectrum, (d) HMBC spectrum, (e) COSY spectrum. CDCl3 was used as the solvent, and CDCl3 peaks were referenced to 7.26 and 77.00 ppm for 1H and 13C spectra, respectively.

### Slide 20
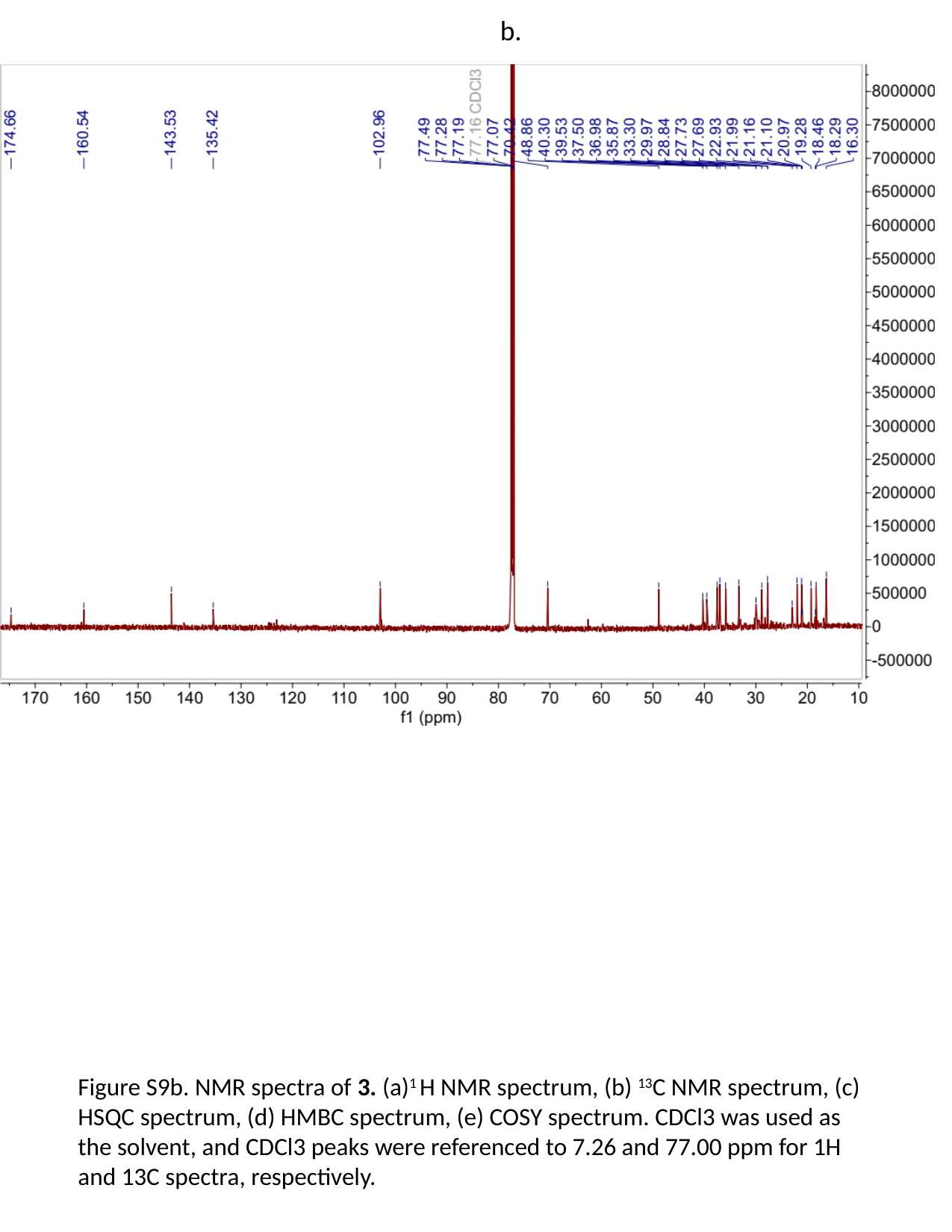

b.
Figure S9b. NMR spectra of 3. (a)^1^ H NMR spectrum, (b) ^13^C NMR spectrum, (c) HSQC spectrum, (d) HMBC spectrum, (e) COSY spectrum. CDCl3 was used as the solvent, and CDCl3 peaks were referenced to 7.26 and 77.00 ppm for 1H and 13C spectra, respectively.

### Slide 21
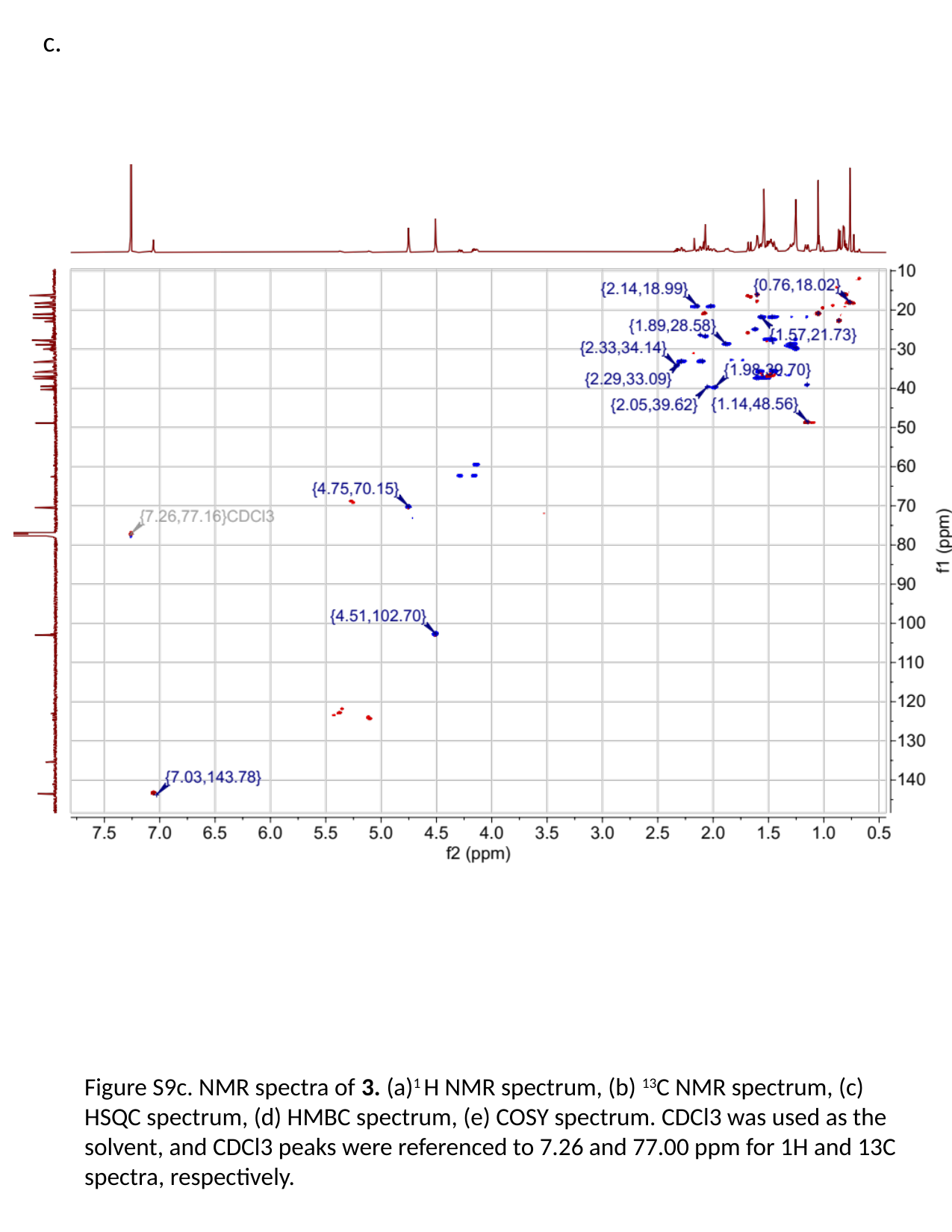

c.
Figure S9c. NMR spectra of 3. (a)^1^ H NMR spectrum, (b) ^13^C NMR spectrum, (c) HSQC spectrum, (d) HMBC spectrum, (e) COSY spectrum. CDCl3 was used as the solvent, and CDCl3 peaks were referenced to 7.26 and 77.00 ppm for 1H and 13C spectra, respectively.

### Slide 22
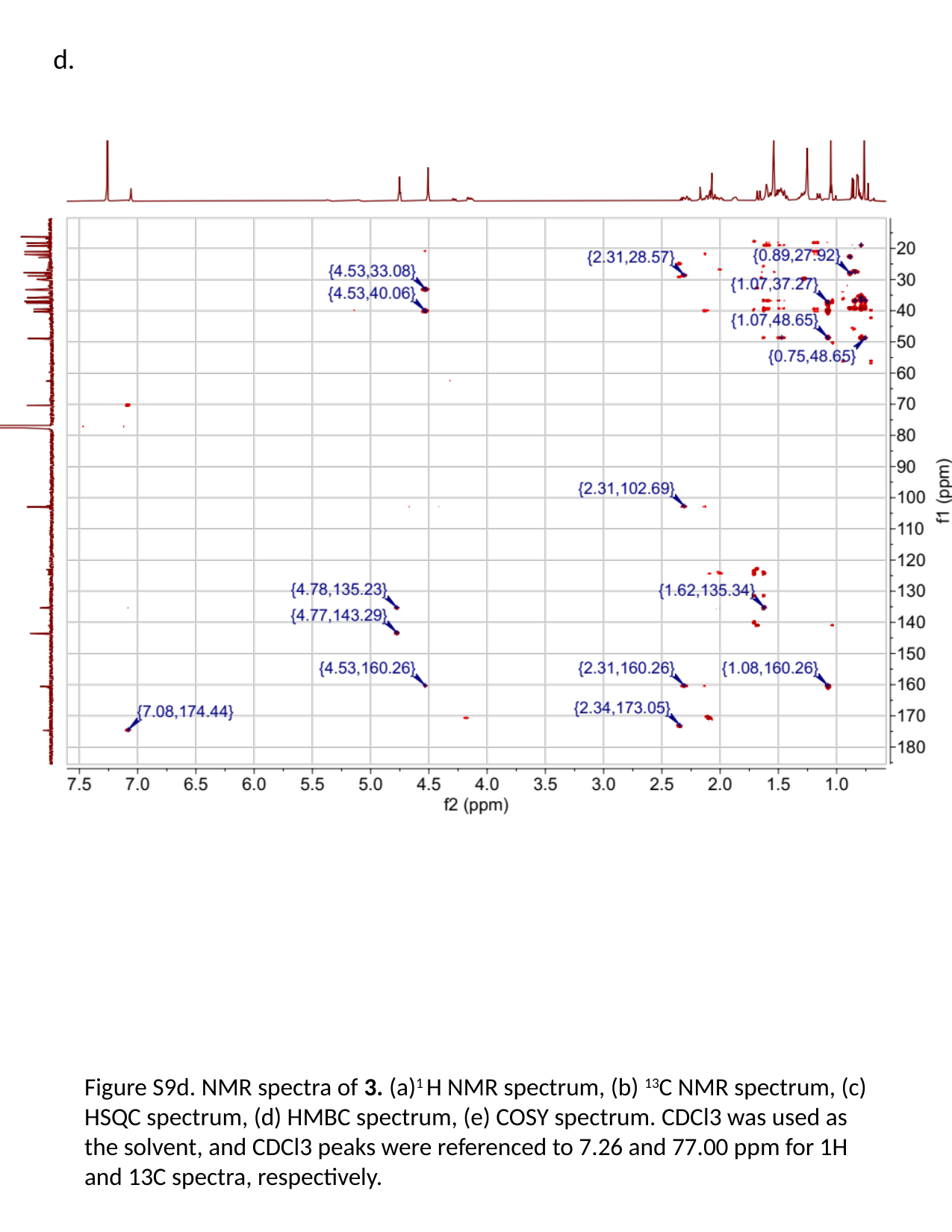

d.
Figure S9d. NMR spectra of 3. (a)^1^ H NMR spectrum, (b) ^13^C NMR spectrum, (c) HSQC spectrum, (d) HMBC spectrum, (e) COSY spectrum. CDCl3 was used as the solvent, and CDCl3 peaks were referenced to 7.26 and 77.00 ppm for 1H and 13C spectra, respectively.

### Slide 23
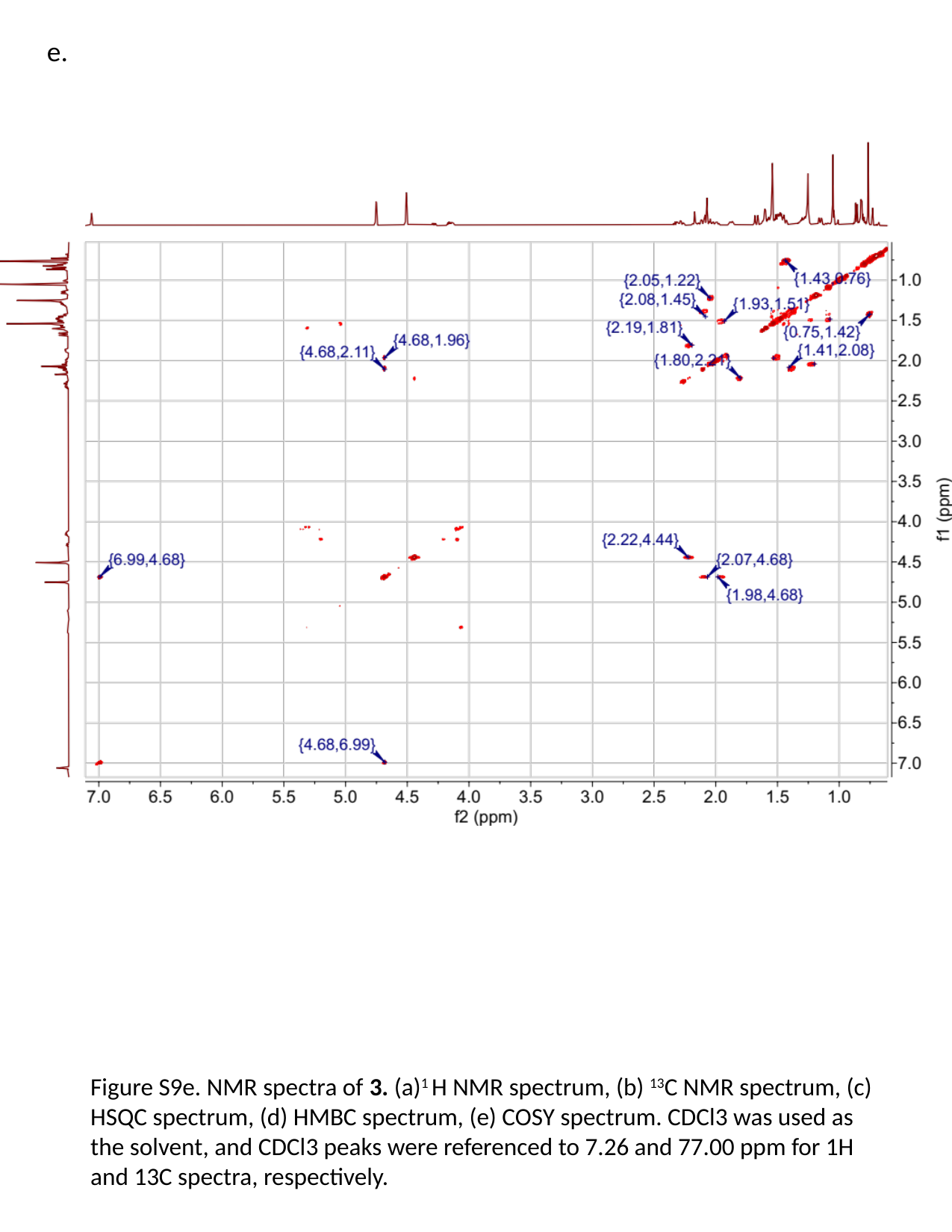

e.
Figure S9e. NMR spectra of 3. (a)^1^ H NMR spectrum, (b) ^13^C NMR spectrum, (c) HSQC spectrum, (d) HMBC spectrum, (e) COSY spectrum. CDCl3 was used as the solvent, and CDCl3 peaks were referenced to 7.26 and 77.00 ppm for 1H and 13C spectra, respectively.

### Slide 24
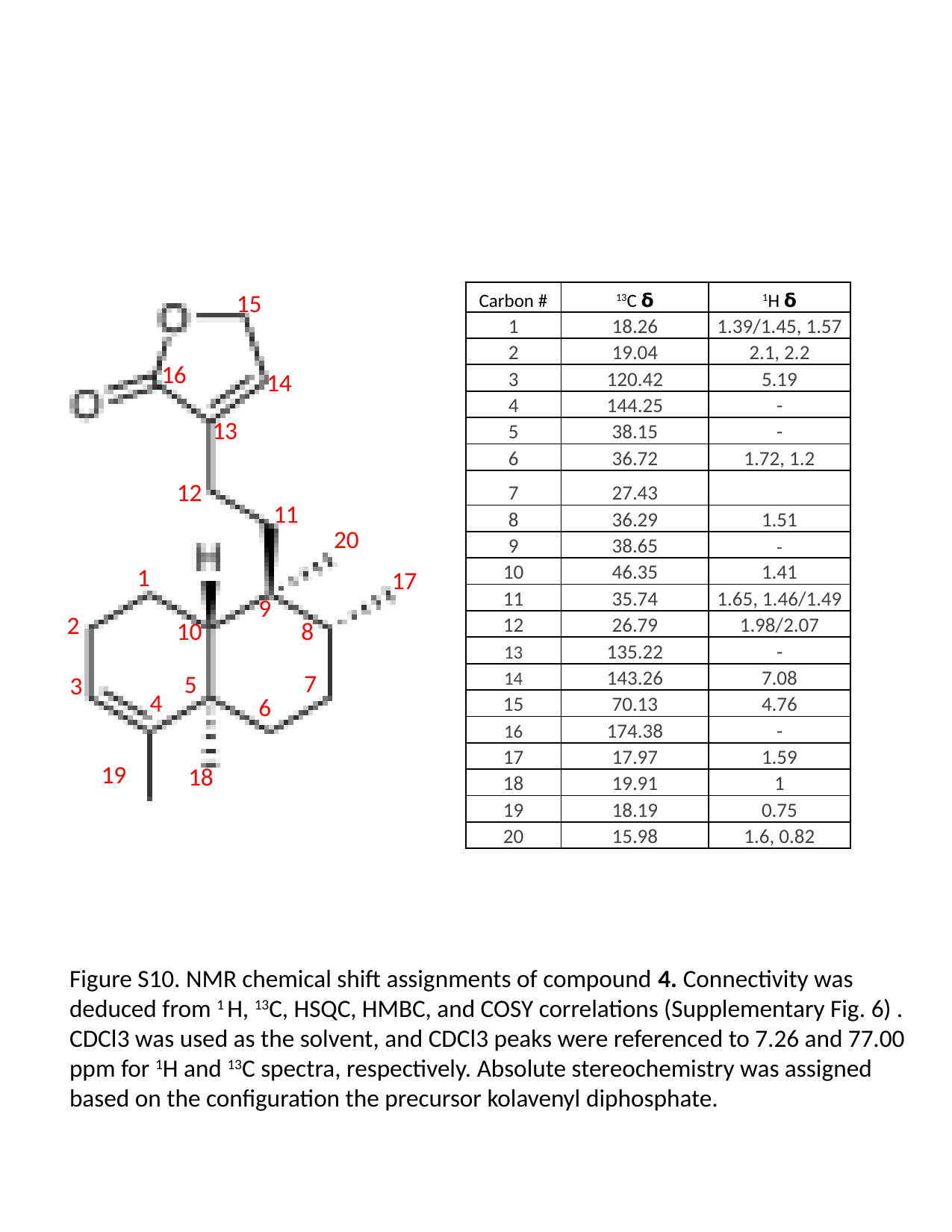

15
16
14
13
12
11
20
1
17
9
2
8
10
7
5
3
4
6
19
18
| Carbon # | ^13^C 𝝳 | ^1^H 𝝳 |
| --- | --- | --- |
| 1 | 18.26 | 1.39/1.45, 1.57 |
| 2 | 19.04 | 2.1, 2.2 |
| 3 | 120.42 | 5.19 |
| 4 | 144.25 | - |
| 5 | 38.15 | - |
| 6 | 36.72 | 1.72, 1.2 |
| 7 | 27.43 | |
| 8 | 36.29 | 1.51 |
| 9 | 38.65 | - |
| 10 | 46.35 | 1.41 |
| 11 | 35.74 | 1.65, 1.46/1.49 |
| 12 | 26.79 | 1.98/2.07 |
| 13 | 135.22 | - |
| 14 | 143.26 | 7.08 |
| 15 | 70.13 | 4.76 |
| 16 | 174.38 | - |
| 17 | 17.97 | 1.59 |
| 18 | 19.91 | 1 |
| 19 | 18.19 | 0.75 |
| 20 | 15.98 | 1.6, 0.82 |
Figure S10. NMR chemical shift assignments of compound 4. Connectivity was deduced from ^1^ H, ^13^C, HSQC, HMBC, and COSY correlations (Supplementary Fig. 6) . CDCl3 was used as the solvent, and CDCl3 peaks were referenced to 7.26 and 77.00 ppm for ^1^H and ^13^C spectra, respectively. Absolute stereochemistry was assigned based on the configuration the precursor kolavenyl diphosphate.

### Slide 25
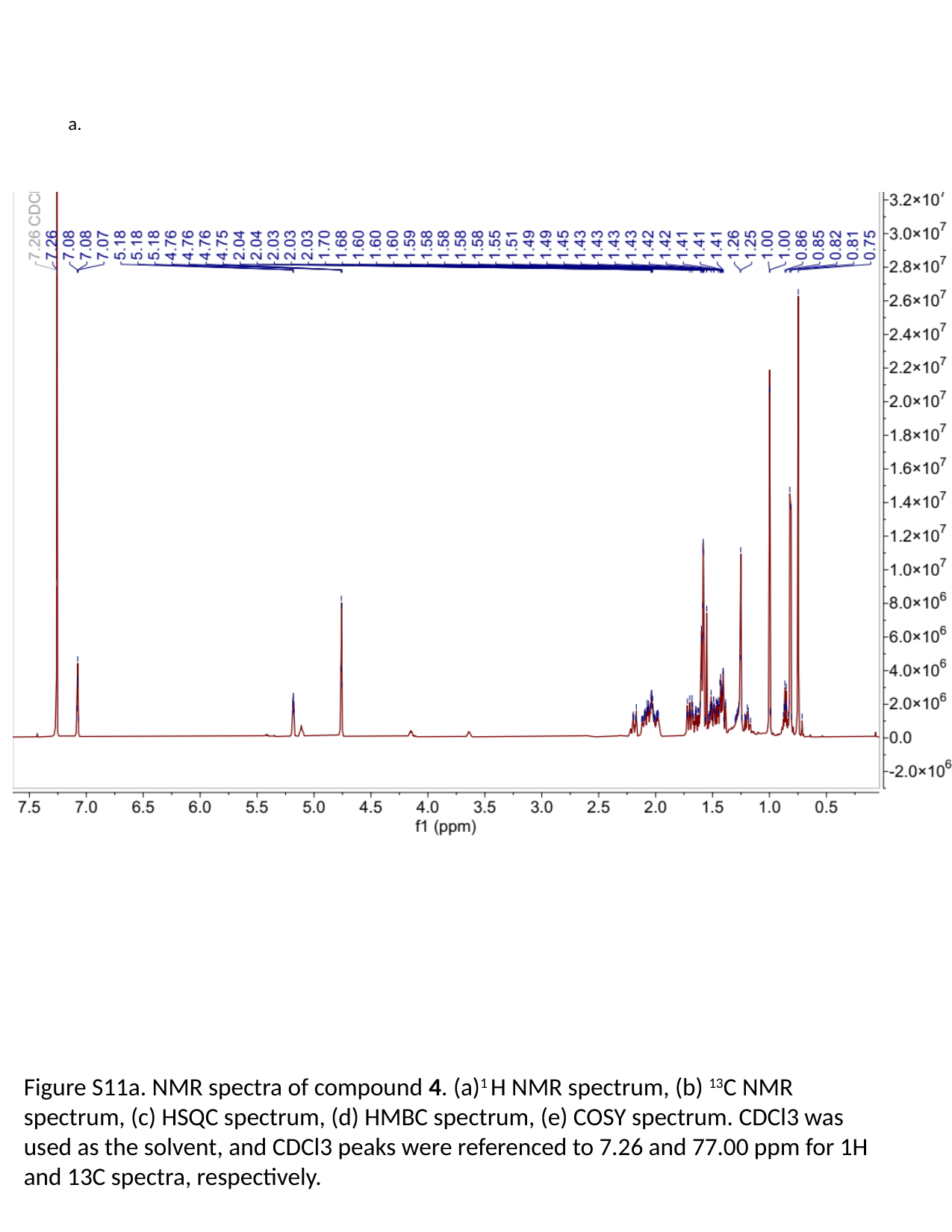

a.
Figure S11a. NMR spectra of compound 4. (a)^1^ H NMR spectrum, (b) ^13^C NMR spectrum, (c) HSQC spectrum, (d) HMBC spectrum, (e) COSY spectrum. CDCl3 was used as the solvent, and CDCl3 peaks were referenced to 7.26 and 77.00 ppm for 1H and 13C spectra, respectively.

### Slide 26
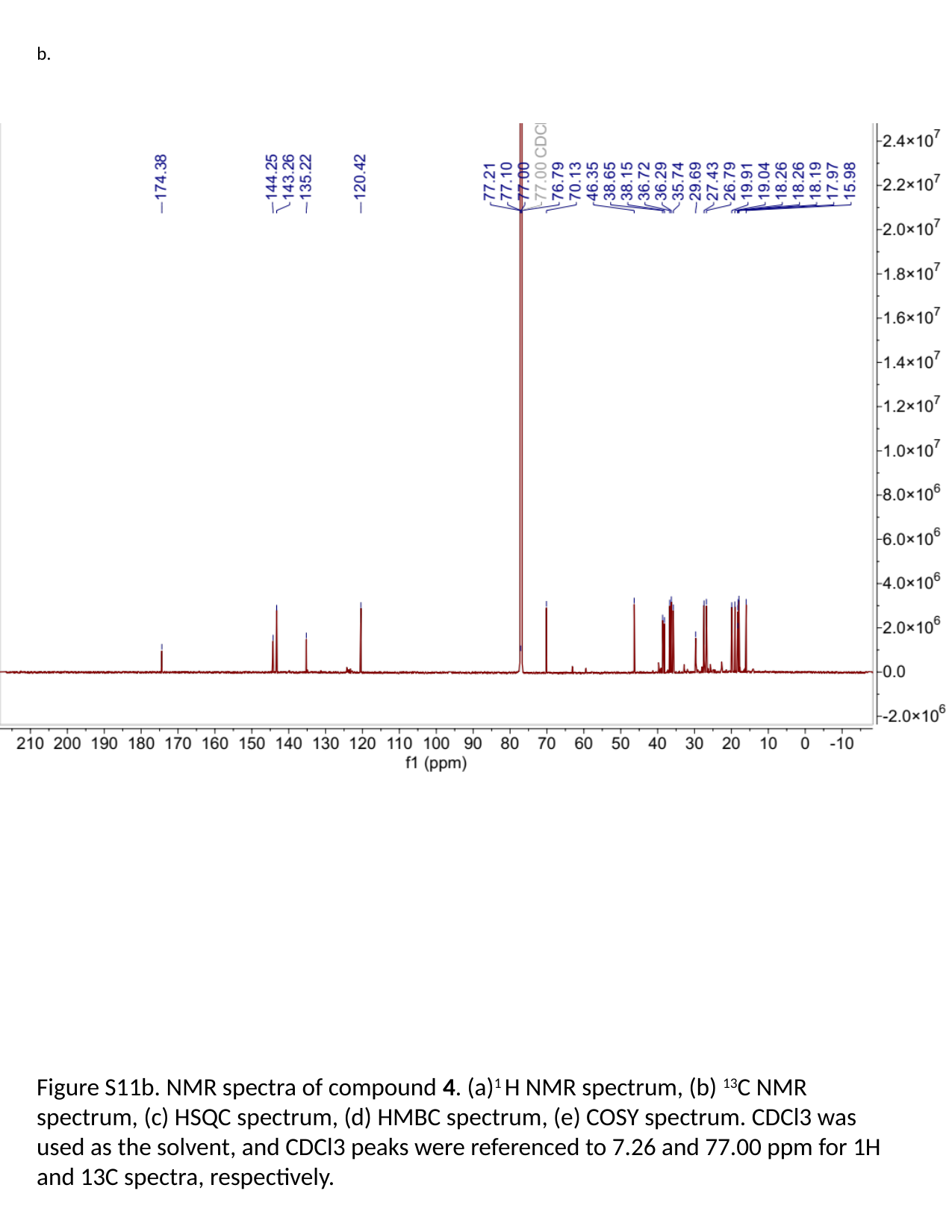

b.
Figure S11b. NMR spectra of compound 4. (a)^1^ H NMR spectrum, (b) ^13^C NMR spectrum, (c) HSQC spectrum, (d) HMBC spectrum, (e) COSY spectrum. CDCl3 was used as the solvent, and CDCl3 peaks were referenced to 7.26 and 77.00 ppm for 1H and 13C spectra, respectively.

### Slide 27
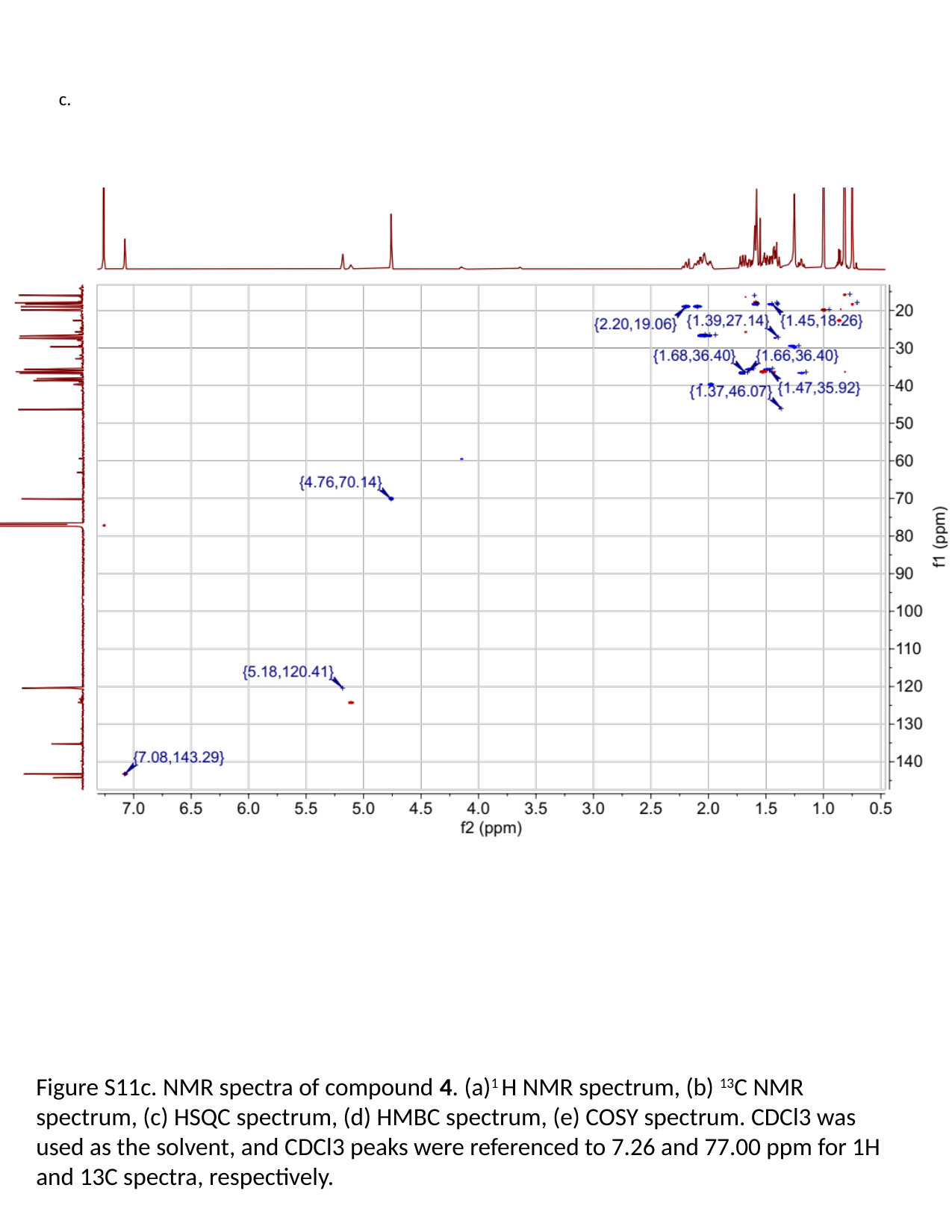

c.
Figure S11c. NMR spectra of compound 4. (a)^1^ H NMR spectrum, (b) ^13^C NMR spectrum, (c) HSQC spectrum, (d) HMBC spectrum, (e) COSY spectrum. CDCl3 was used as the solvent, and CDCl3 peaks were referenced to 7.26 and 77.00 ppm for 1H and 13C spectra, respectively.

### Slide 28
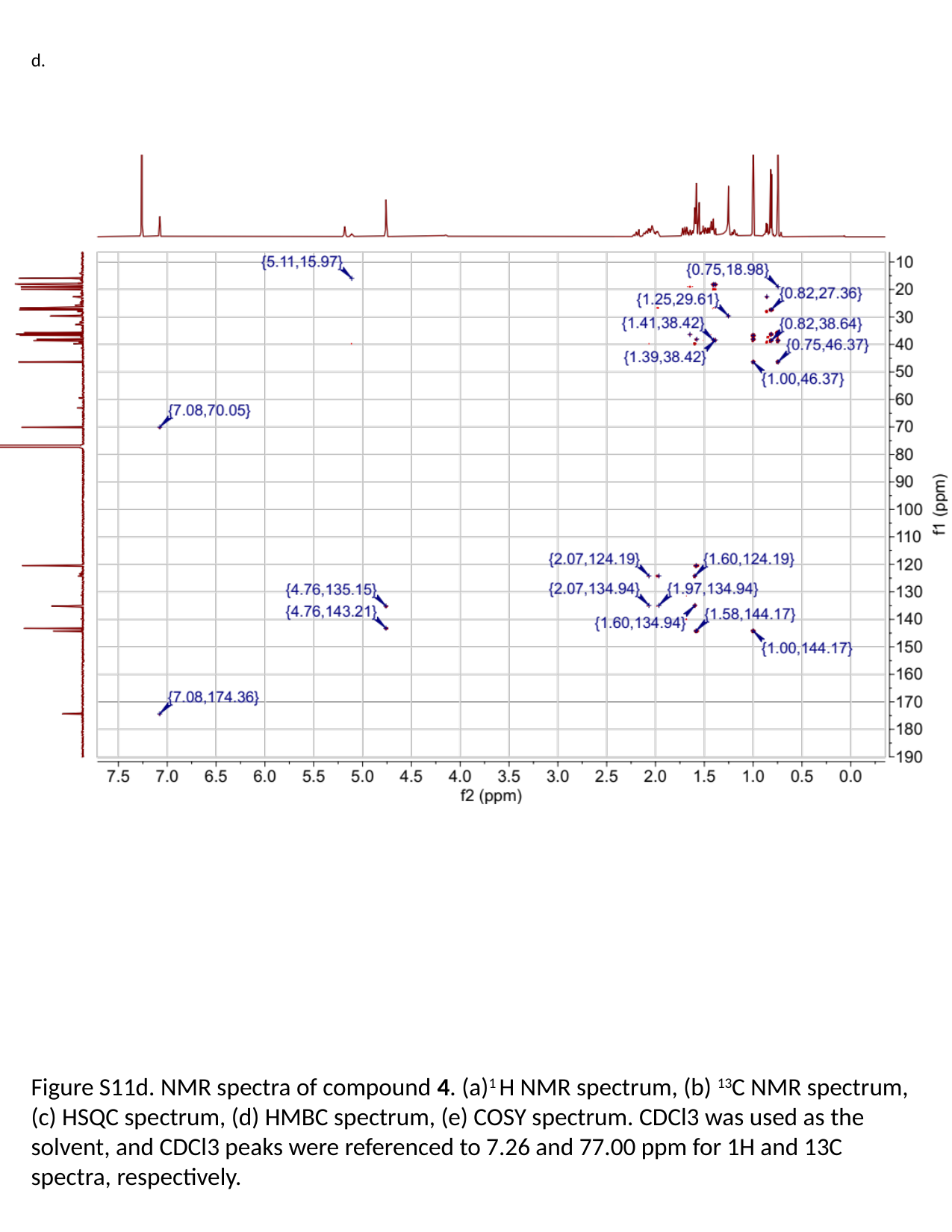

d.
Figure S11d. NMR spectra of compound 4. (a)^1^ H NMR spectrum, (b) ^13^C NMR spectrum, (c) HSQC spectrum, (d) HMBC spectrum, (e) COSY spectrum. CDCl3 was used as the solvent, and CDCl3 peaks were referenced to 7.26 and 77.00 ppm for 1H and 13C spectra, respectively.

### Slide 29
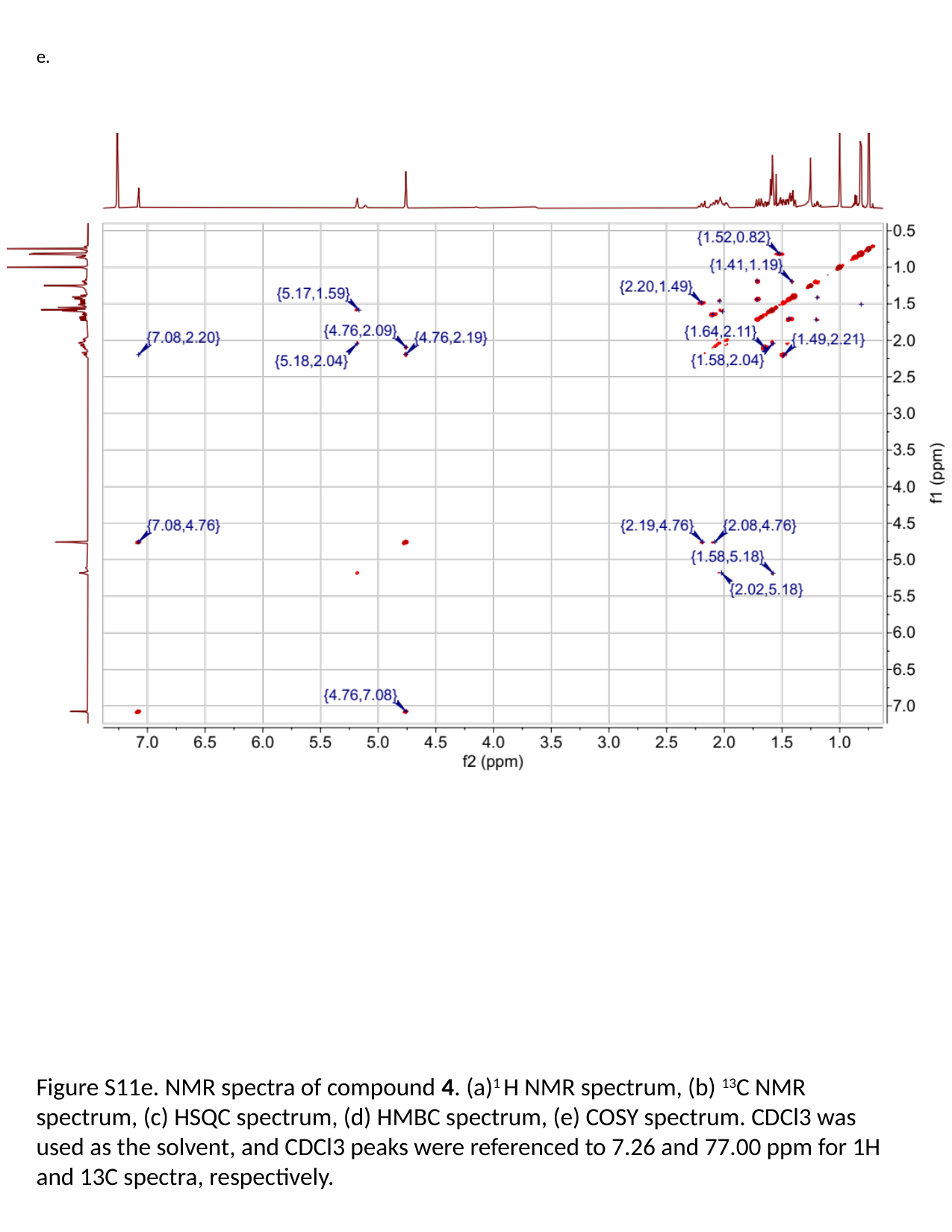

e.
Figure S11e. NMR spectra of compound 4. (a)^1^ H NMR spectrum, (b) ^13^C NMR spectrum, (c) HSQC spectrum, (d) HMBC spectrum, (e) COSY spectrum. CDCl3 was used as the solvent, and CDCl3 peaks were referenced to 7.26 and 77.00 ppm for 1H and 13C spectra, respectively.

### Slide 30
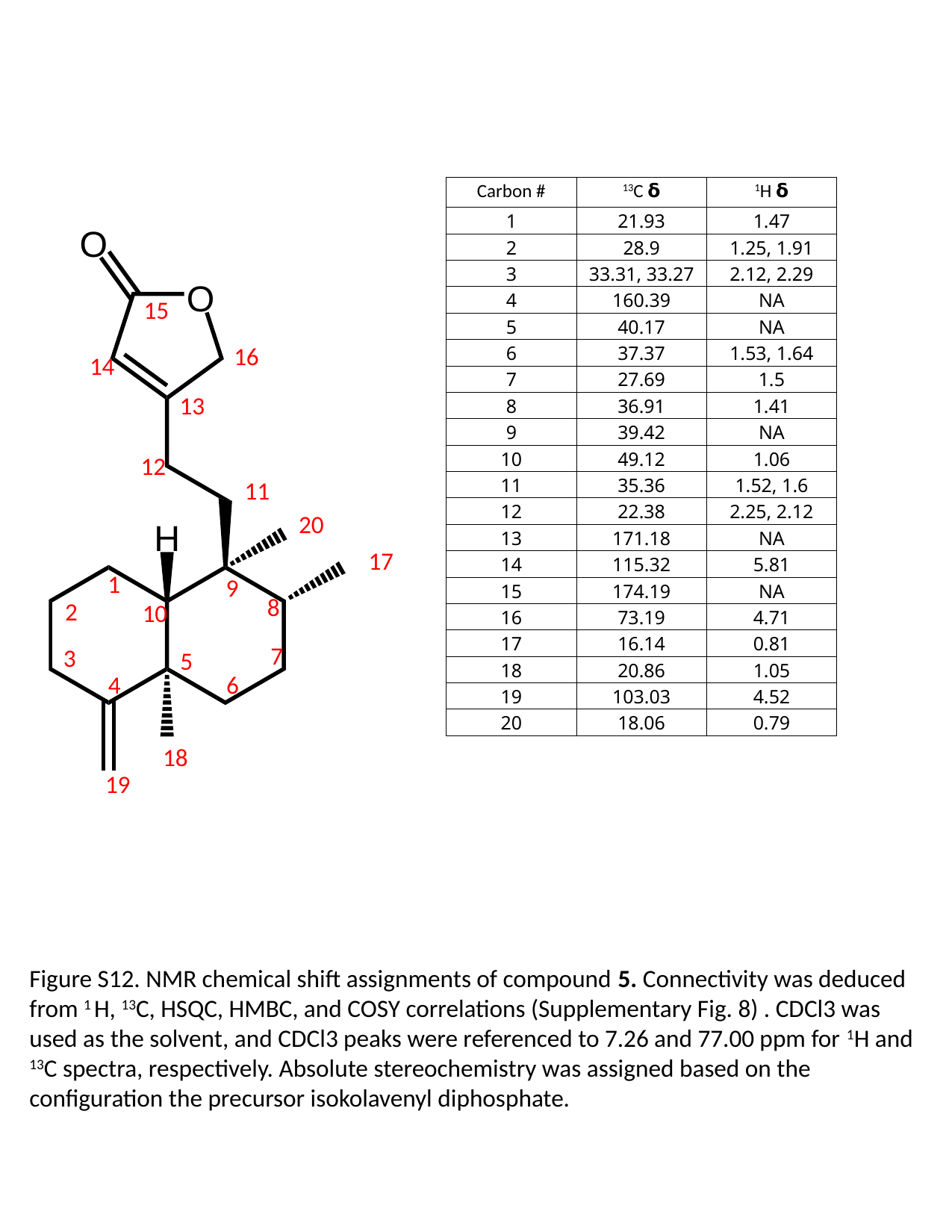

| Carbon # | ^13^C 𝝳 | ^1^H 𝝳 |
| --- | --- | --- |
| 1 | 21.93 | 1.47 |
| 2 | 28.9 | 1.25, 1.91 |
| 3 | 33.31, 33.27 | 2.12, 2.29 |
| 4 | 160.39 | NA |
| 5 | 40.17 | NA |
| 6 | 37.37 | 1.53, 1.64 |
| 7 | 27.69 | 1.5 |
| 8 | 36.91 | 1.41 |
| 9 | 39.42 | NA |
| 10 | 49.12 | 1.06 |
| 11 | 35.36 | 1.52, 1.6 |
| 12 | 22.38 | 2.25, 2.12 |
| 13 | 171.18 | NA |
| 14 | 115.32 | 5.81 |
| 15 | 174.19 | NA |
| 16 | 73.19 | 4.71 |
| 17 | 16.14 | 0.81 |
| 18 | 20.86 | 1.05 |
| 19 | 103.03 | 4.52 |
| 20 | 18.06 | 0.79 |
15
16
14
13
12
11
20
17
1
9
8
2
10
7
3
5
4
6
18
19
Figure S12. NMR chemical shift assignments of compound 5. Connectivity was deduced from ^1^ H, ^13^C, HSQC, HMBC, and COSY correlations (Supplementary Fig. 8) . CDCl3 was used as the solvent, and CDCl3 peaks were referenced to 7.26 and 77.00 ppm for ^1^H and ^13^C spectra, respectively. Absolute stereochemistry was assigned based on the configuration the precursor isokolavenyl diphosphate.

### Slide 31
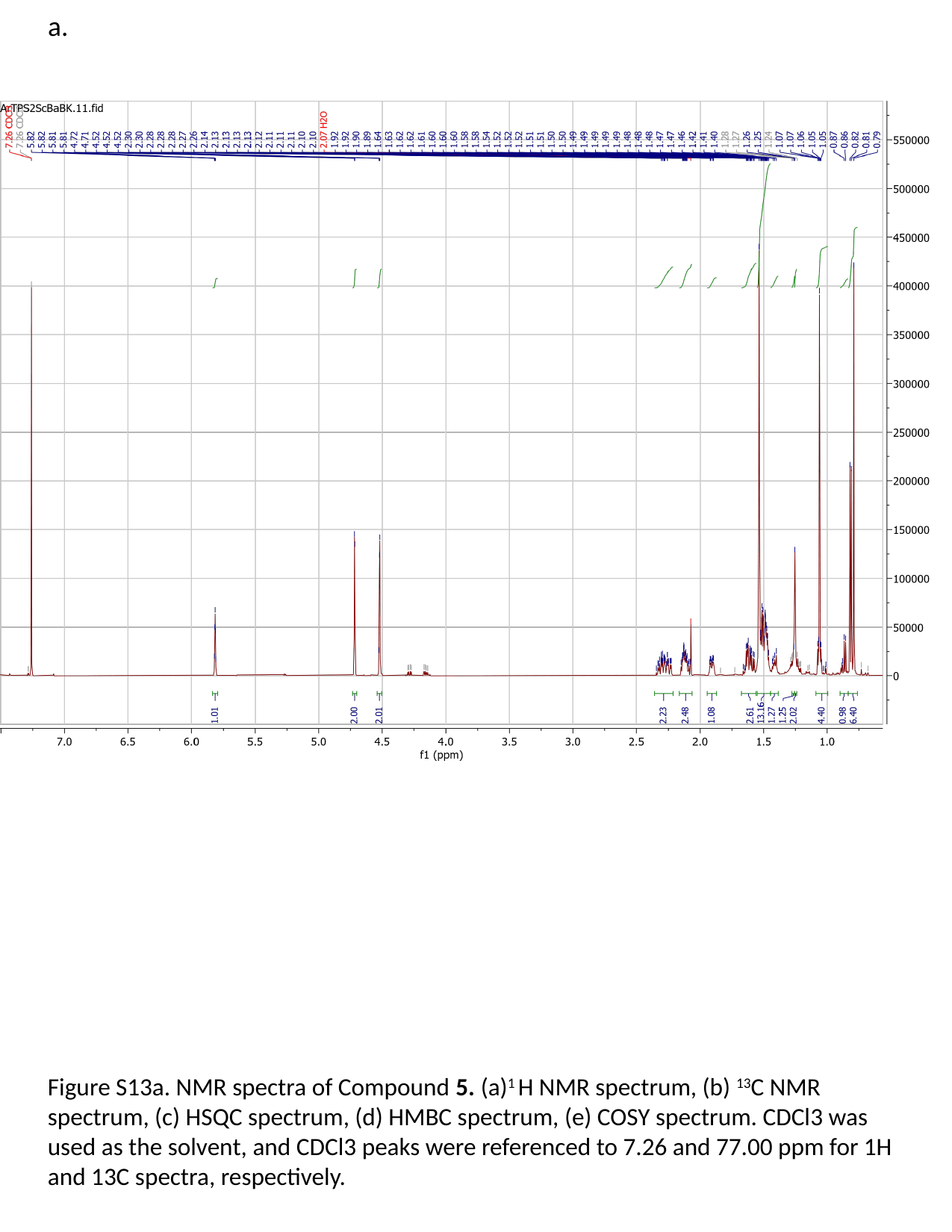

a.
Figure S13a. NMR spectra of Compound 5. (a)^1^ H NMR spectrum, (b) ^13^C NMR spectrum, (c) HSQC spectrum, (d) HMBC spectrum, (e) COSY spectrum. CDCl3 was used as the solvent, and CDCl3 peaks were referenced to 7.26 and 77.00 ppm for 1H and 13C spectra, respectively.

### Slide 32
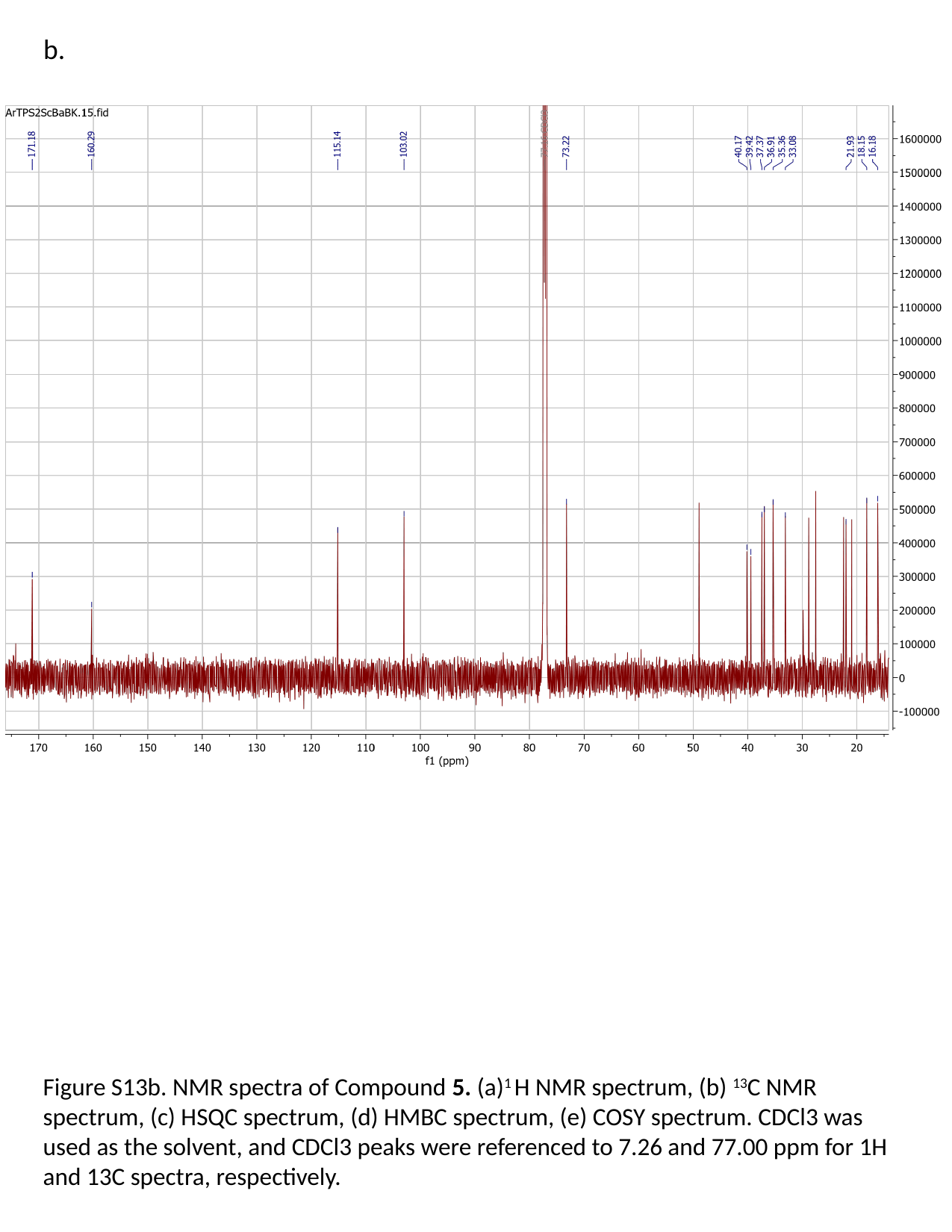

b.
Figure S13b. NMR spectra of Compound 5. (a)^1^ H NMR spectrum, (b) ^13^C NMR spectrum, (c) HSQC spectrum, (d) HMBC spectrum, (e) COSY spectrum. CDCl3 was used as the solvent, and CDCl3 peaks were referenced to 7.26 and 77.00 ppm for 1H and 13C spectra, respectively.

### Slide 33
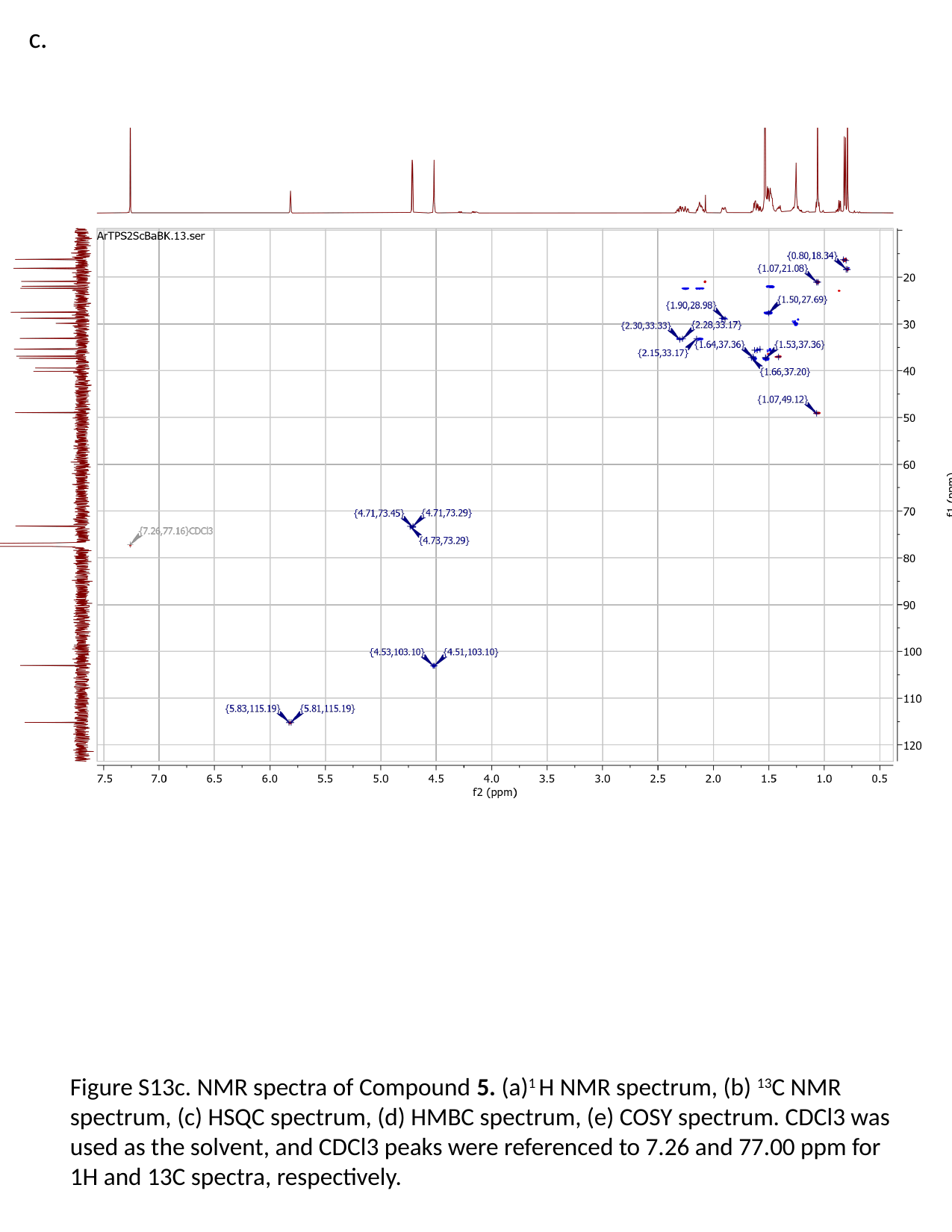

c.
Figure S13c. NMR spectra of Compound 5. (a)^1^ H NMR spectrum, (b) ^13^C NMR spectrum, (c) HSQC spectrum, (d) HMBC spectrum, (e) COSY spectrum. CDCl3 was used as the solvent, and CDCl3 peaks were referenced to 7.26 and 77.00 ppm for 1H and 13C spectra, respectively.

### Slide 34
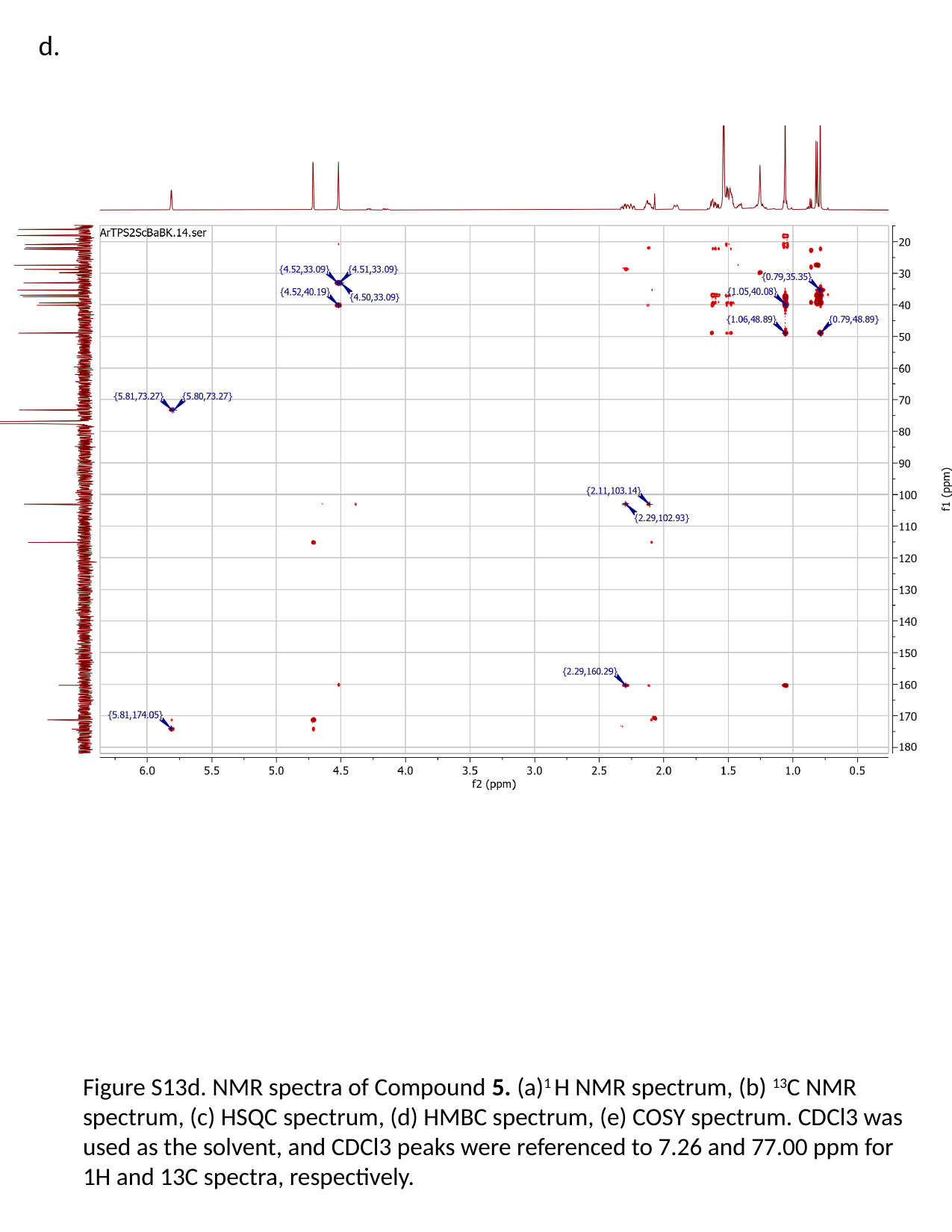

d.
Figure S13d. NMR spectra of Compound 5. (a)^1^ H NMR spectrum, (b) ^13^C NMR spectrum, (c) HSQC spectrum, (d) HMBC spectrum, (e) COSY spectrum. CDCl3 was used as the solvent, and CDCl3 peaks were referenced to 7.26 and 77.00 ppm for 1H and 13C spectra, respectively.

### Slide 35
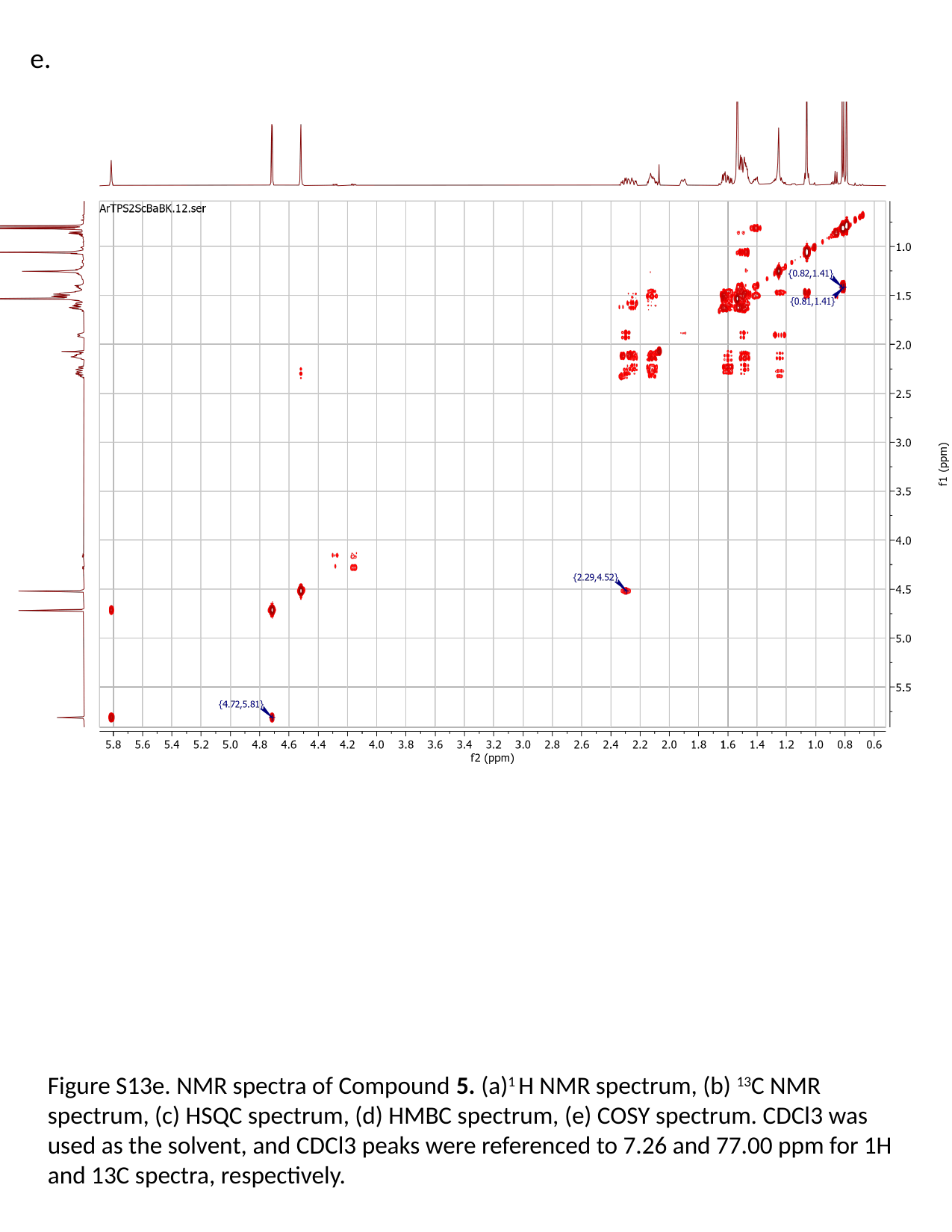

e.
Figure S13e. NMR spectra of Compound 5. (a)^1^ H NMR spectrum, (b) ^13^C NMR spectrum, (c) HSQC spectrum, (d) HMBC spectrum, (e) COSY spectrum. CDCl3 was used as the solvent, and CDCl3 peaks were referenced to 7.26 and 77.00 ppm for 1H and 13C spectra, respectively.

### Slide 36
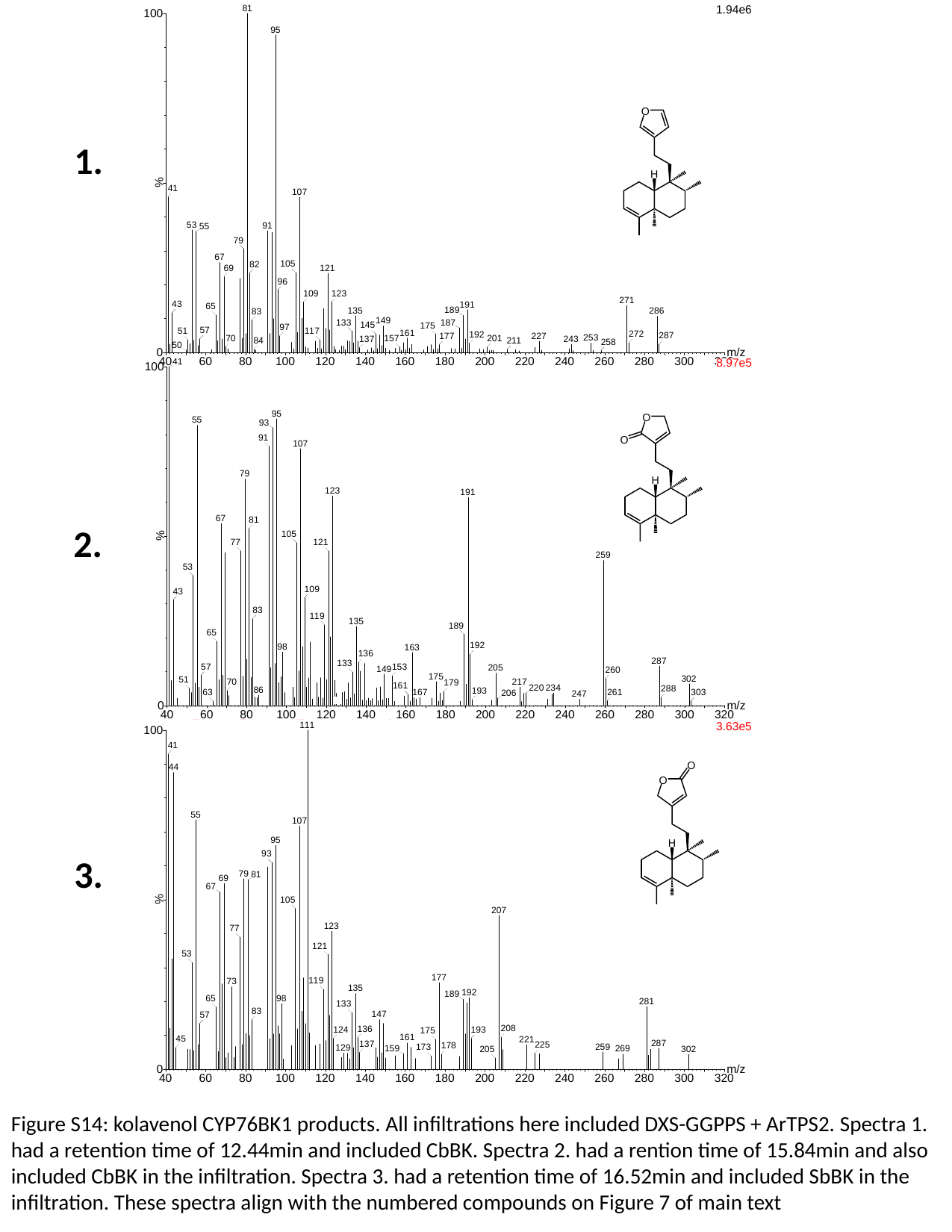

1.
2.
3.
Figure S14: kolavenol CYP76BK1 products. All infiltrations here included DXS-GGPPS + ArTPS2. Spectra 1. had a retention time of 12.44min and included CbBK. Spectra 2. had a rention time of 15.84min and also included CbBK in the infiltration. Spectra 3. had a retention time of 16.52min and included SbBK in the infiltration. These spectra align with the numbered compounds on Figure 7 of main text

### Slide 37
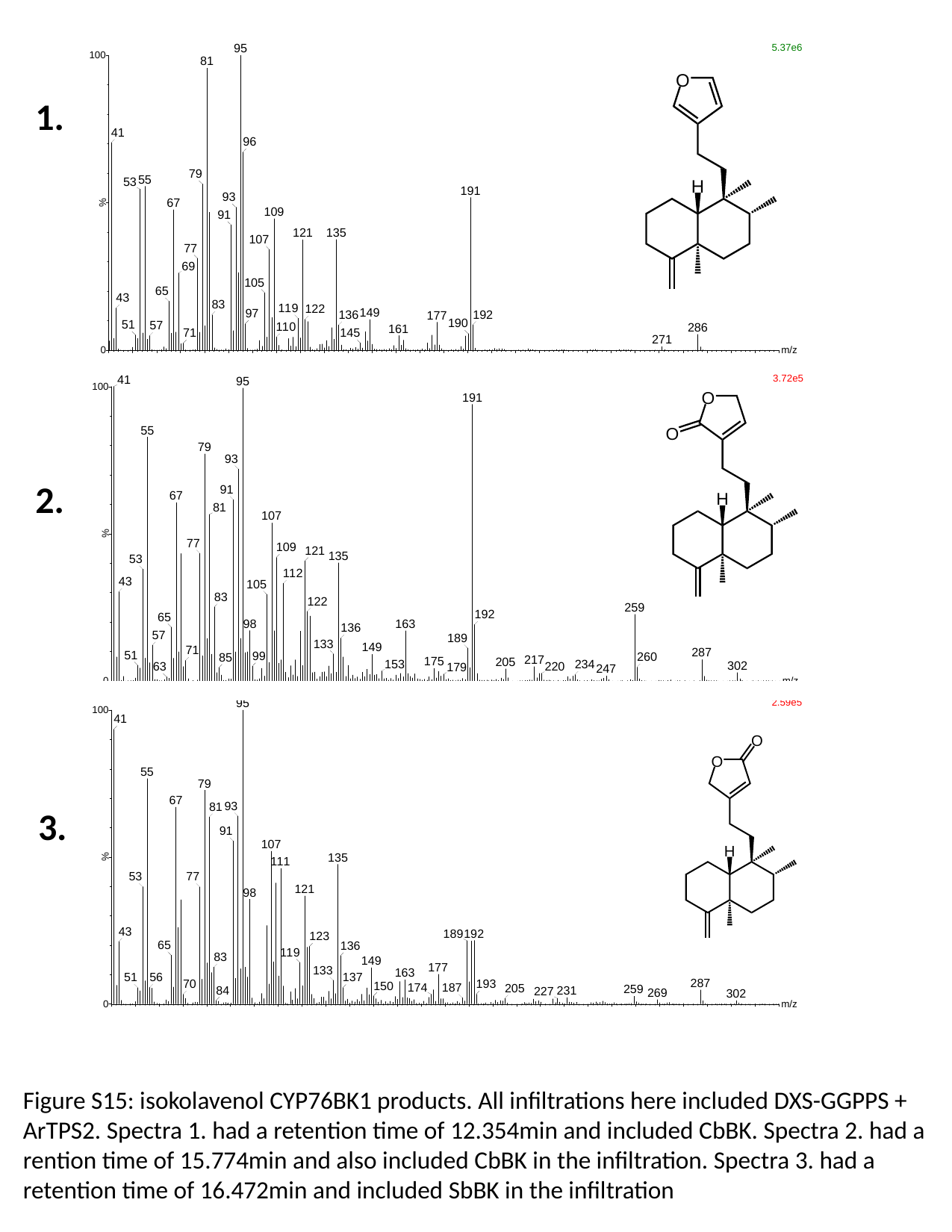

1.
2.
3.
Figure S15: isokolavenol CYP76BK1 products. All infiltrations here included DXS-GGPPS + ArTPS2. Spectra 1. had a retention time of 12.354min and included CbBK. Spectra 2. had a rention time of 15.774min and also included CbBK in the infiltration. Spectra 3. had a retention time of 16.472min and included SbBK in the infiltration

### Slide 38
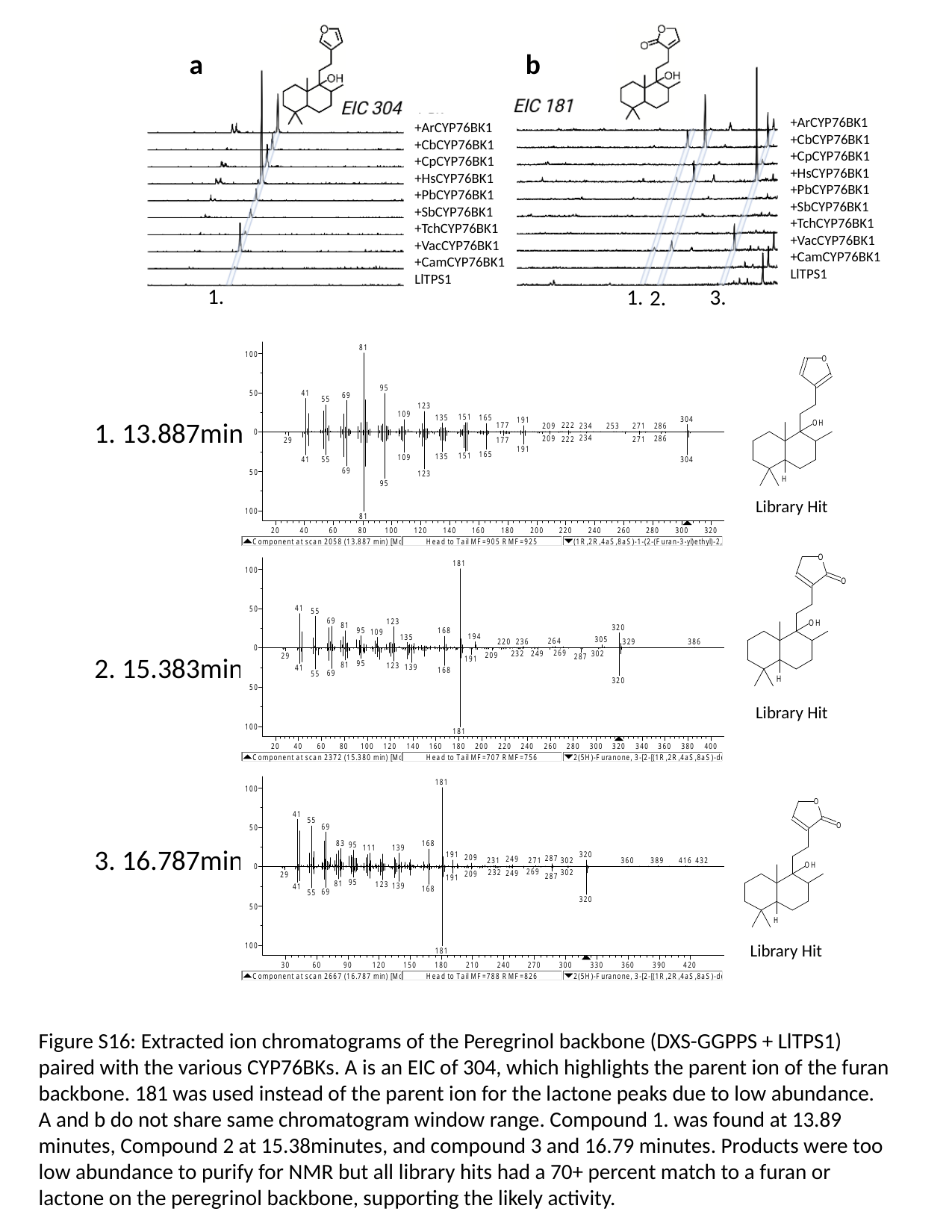

a
b
+ArCYP76BK1
+CbCYP76BK1
+CpCYP76BK1
+HsCYP76BK1
+PbCYP76BK1
+SbCYP76BK1
+TchCYP76BK1
+VacCYP76BK1
+CamCYP76BK1
LlTPS1
+ArCYP76BK1
+CbCYP76BK1
+CpCYP76BK1
+HsCYP76BK1
+PbCYP76BK1
+SbCYP76BK1
+TchCYP76BK1
+VacCYP76BK1
+CamCYP76BK1
LlTPS1
1.
1.
3.
2.
1. 13.887min
Library Hit
2. 15.383min
Library Hit
3. 16.787min
Library Hit
Figure S16: Extracted ion chromatograms of the Peregrinol backbone (DXS-GGPPS + LlTPS1) paired with the various CYP76BKs. A is an EIC of 304, which highlights the parent ion of the furan backbone. 181 was used instead of the parent ion for the lactone peaks due to low abundance. A and b do not share same chromatogram window range. Compound 1. was found at 13.89 minutes, Compound 2 at 15.38minutes, and compound 3 and 16.79 minutes. Products were too low abundance to purify for NMR but all library hits had a 70+ percent match to a furan or lactone on the peregrinol backbone, supporting the likely activity.

### Slide 39
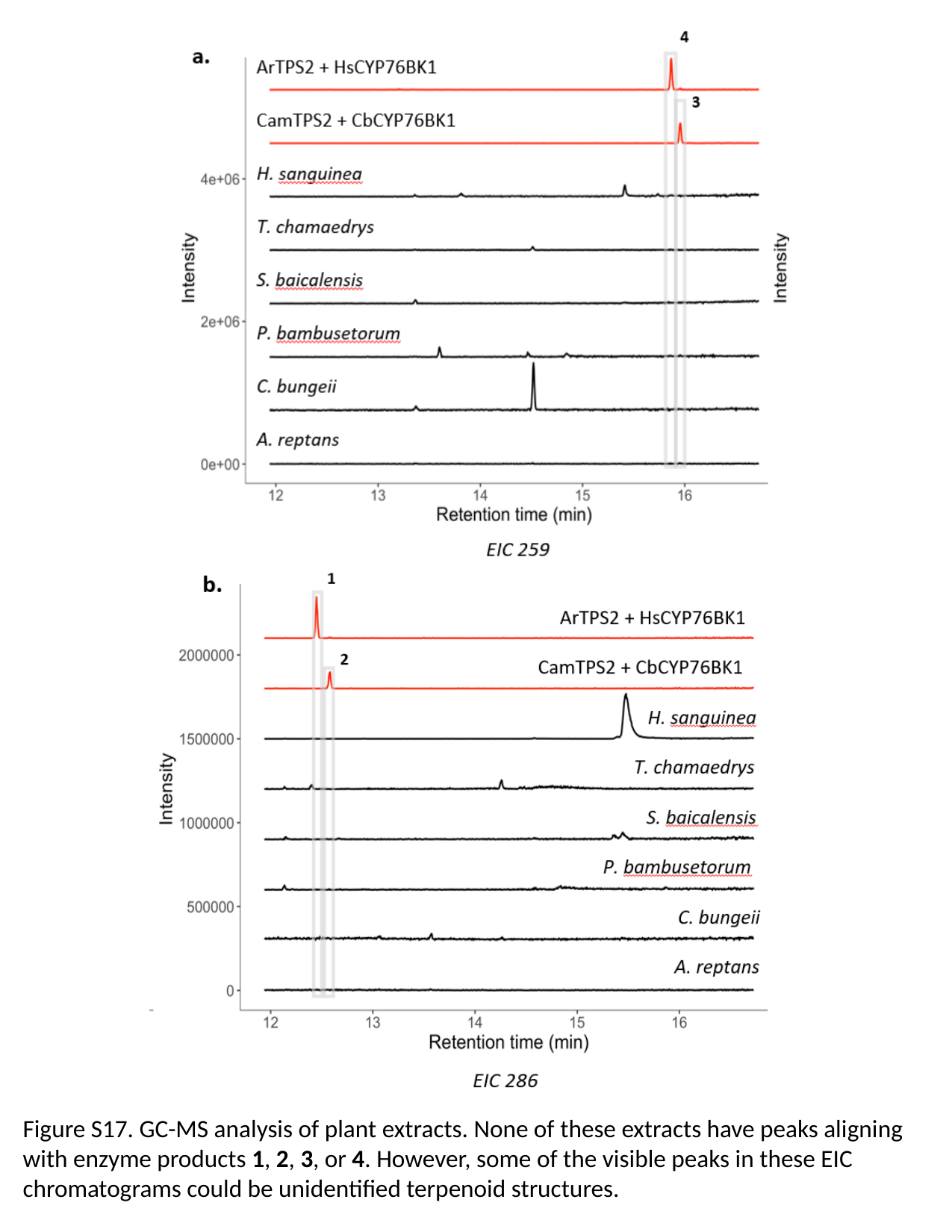

Figure S17. GC-MS analysis of plant extracts. None of these extracts have peaks aligning with enzyme products 1, 2, 3, or 4. However, some of the visible peaks in these EIC chromatograms could be unidentified terpenoid structures.

### Slide 40
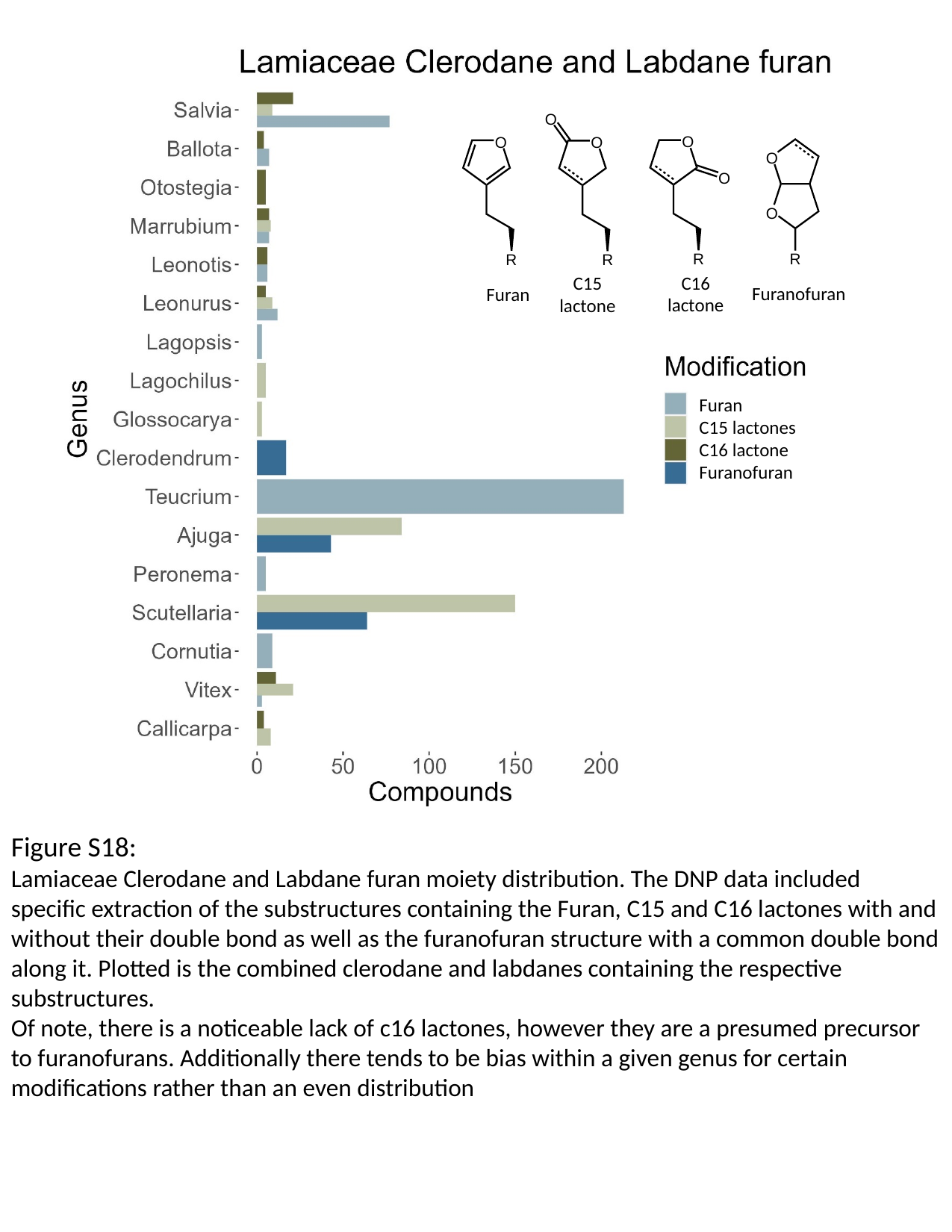

C16 lactone
C15 lactone
Furanofuran
Furan
Furan
C15 lactones
C16 lactone
Furanofuran
Figure S18:
Lamiaceae Clerodane and Labdane furan moiety distribution. The DNP data included specific extraction of the substructures containing the Furan, C15 and C16 lactones with and without their double bond as well as the furanofuran structure with a common double bond along it. Plotted is the combined clerodane and labdanes containing the respective substructures.
Of note, there is a noticeable lack of c16 lactones, however they are a presumed precursor to furanofurans. Additionally there tends to be bias within a given genus for certain modifications rather than an even distribution

### Slide 41
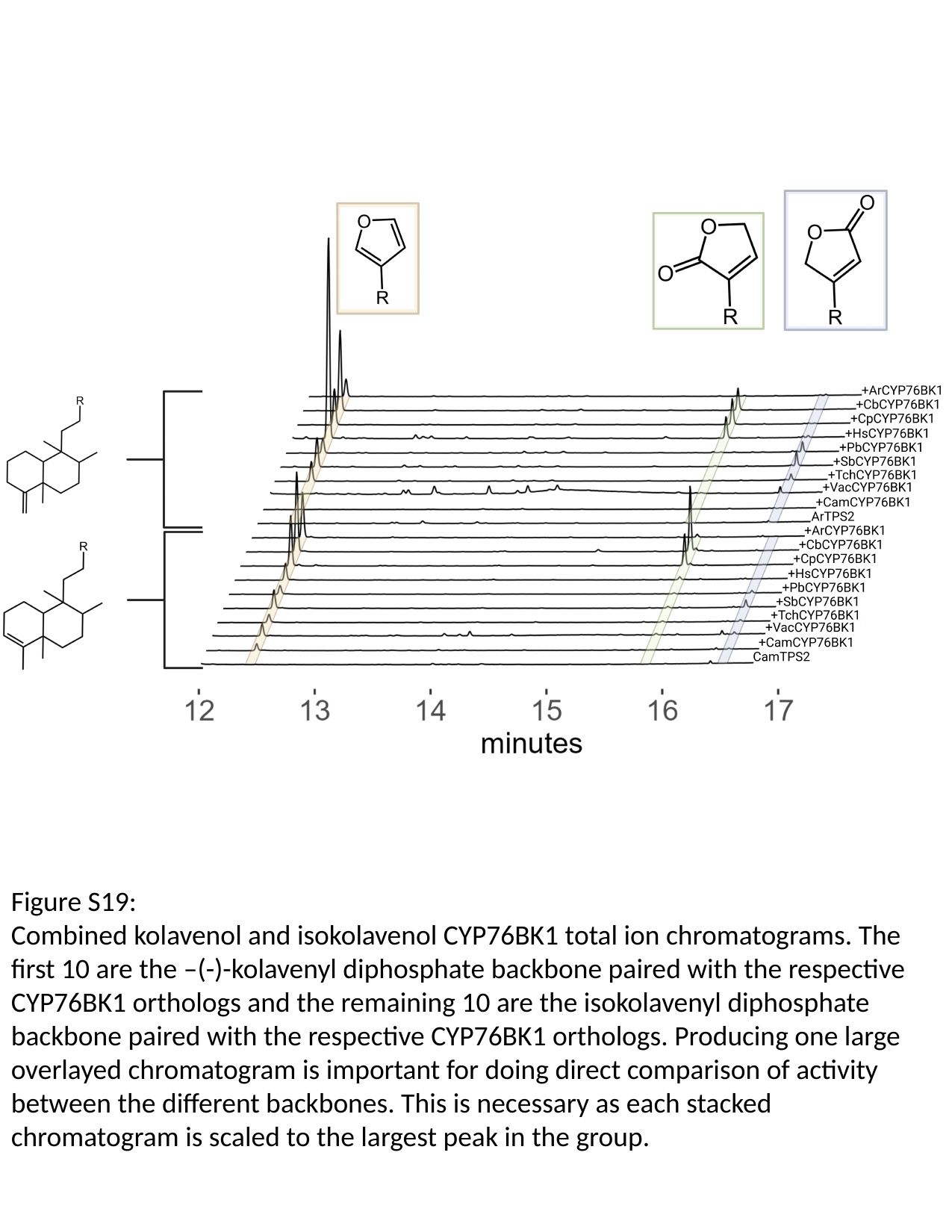

Figure S19:
Combined kolavenol and isokolavenol CYP76BK1 total ion chromatograms. The first 10 are the –(-)-kolavenyl diphosphate backbone paired with the respective CYP76BK1 orthologs and the remaining 10 are the isokolavenyl diphosphate backbone paired with the respective CYP76BK1 orthologs. Producing one large overlayed chromatogram is important for doing direct comparison of activity between the different backbones. This is necessary as each stacked chromatogram is scaled to the largest peak in the group.

### Slide 42
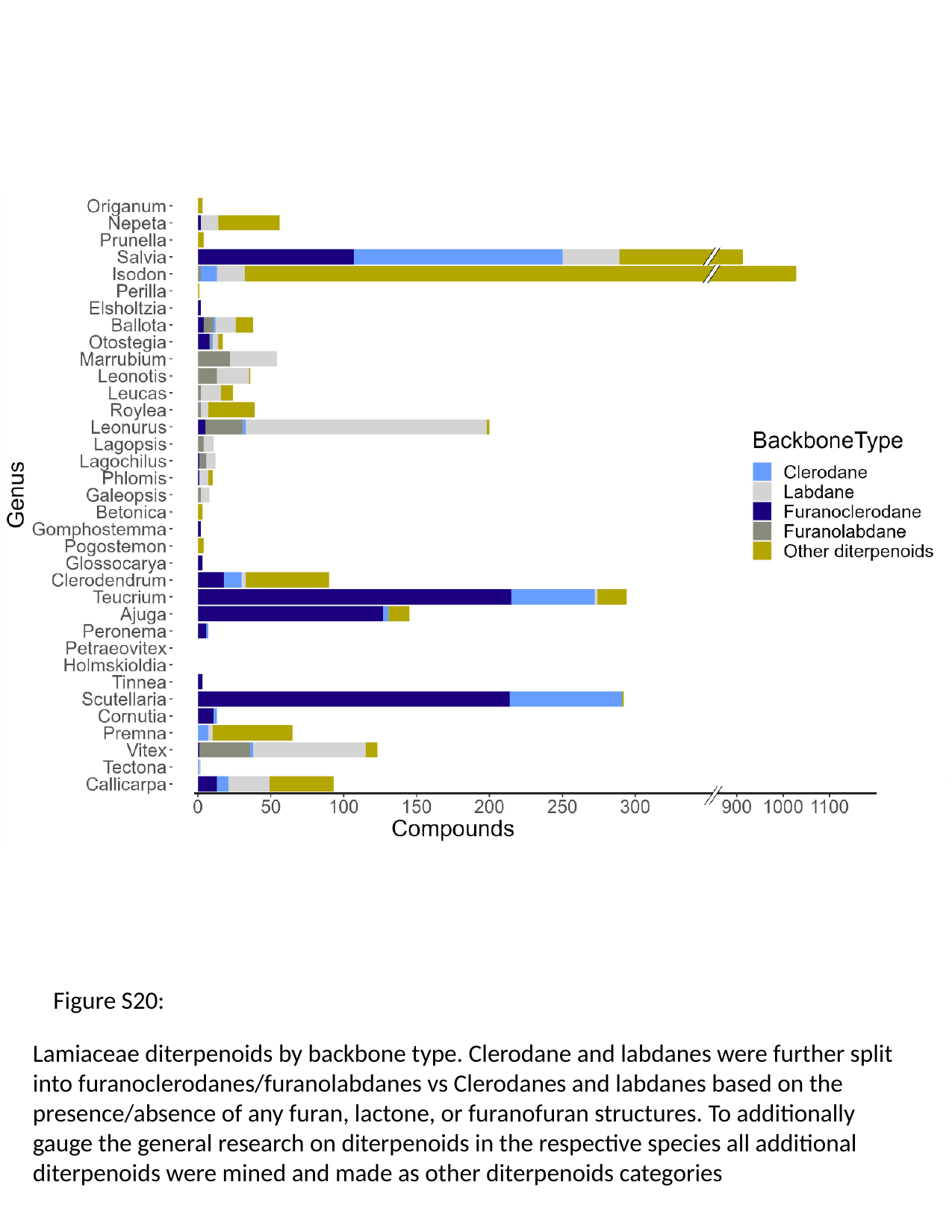

Figure S20:
Lamiaceae diterpenoids by backbone type. Clerodane and labdanes were further split into furanoclerodanes/furanolabdanes vs Clerodanes and labdanes based on the presence/absence of any furan, lactone, or furanofuran structures. To additionally gauge the general research on diterpenoids in the respective species all additional diterpenoids were mined and made as other diterpenoids categories

### Slide 43
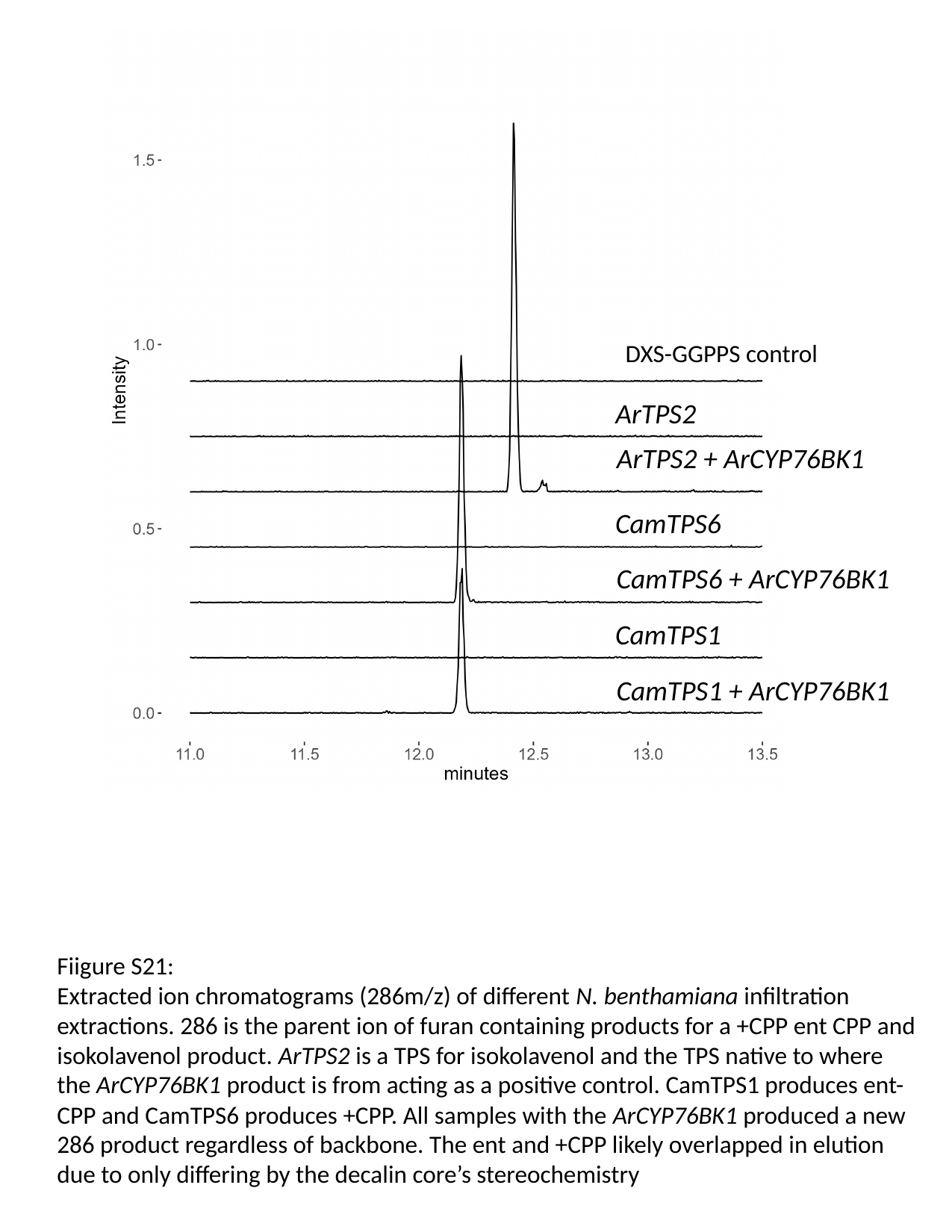

DXS-GGPPS control
ArTPS2
ArTPS2 + ArCYP76BK1
CamTPS6
CamTPS6 + ArCYP76BK1
CamTPS1
CamTPS1 + ArCYP76BK1
Fiigure S21:
Extracted ion chromatograms (286m/z) of different N. benthamiana infiltration extractions. 286 is the parent ion of furan containing products for a +CPP ent CPP and isokolavenol product. ArTPS2 is a TPS for isokolavenol and the TPS native to where the ArCYP76BK1 product is from acting as a positive control. CamTPS1 produces ent-CPP and CamTPS6 produces +CPP. All samples with the ArCYP76BK1 produced a new 286 product regardless of backbone. The ent and +CPP likely overlapped in elution due to only differing by the decalin core’s stereochemistry
